# Coupled nucleoplasmic flows and envelope expansion govern the mechanical resistance of the nucleus

**DOI:** 10.64898/2026.06.12.731963

**Authors:** Romain Rollin, Michele Nava, Alice Williart, Solène Ludwig, Camille Plancke, Marie Aude Plamont, Mabel San Roman, Guilherme P. F Nader, Zoher Gueroui, Jean-François Joanny, Daniel J. Müller, Damien Cuvelier, Pierre Sens, Matthieu Piel

**Affiliations:** Institut Pierre Gilles de Gennes, PSL Research University, 6 rue Jean Calvin, 75005 Paris, France; Institut Curie, PSL Research University, CNRS UMR 144, Paris, France; University of Geneva (UNIGE), Department of Genetics and Evolution, Bd d’Yvoy 4, 1205 Geneve, Switzerland; Eidgenössische Technische Hochschule (ETH) Zurich, Department of Biosystems Science and Engineering, Klingelbergstrasse 48, Basel, Switzerland; CPCV, Department of Chemistry, ENS, PSL University, Sorbonne Université, CNRS, Paris, France; Institute Curie, UMR3215, 75005 Paris, France; Department of Pathology and Laboratory Medicine, Children’s Hospital of Philadelphia & Perelman School of Medicine, University of Pennsylvania, Philadelphia, PA, USA; Institut Curie Physics of Cells and Cancer/Collège de France Chair Soft Matter and Biophysics, Paris Cedex 05, France; Physique of Cells and Cancer, Sorbonne Université, Institut Curie, Université PSL, CNRS UMR168, 75005 Paris, France

## Abstract

In recent year, it became clear that the cell nucleus can undergo large deformations, during immune cell migration and tumor growth. These deformations generate signals that allow cells to sense their environment and adapt to it. How cells cope with and respond to large deformations thus strongly depends on the nuclear mechanics, but our understanding of the physical properties of the nucleus remains incomplete. In particular, it is not clear how the nuclear volume responds to deformation. Here we combine controlled confinement assays, high-resolution imaging and atomic force microscopy with theoretical modelling to propose a physical model of the cell nucleus that accounts for its surface and bulk properties and addresses both steady-state and transient regimes. Our results establish the nucleus as a poroelastic body in which mechanics are dominated by the envelope and dynamics by the chromatin, and suggest that regulating water permeability may be as important as softening the envelope for cells migrating rapidly through dense tissues.

## Introduction

Cells have long been known to respond to mechanical cues (Vogel et al., 2006; Jaalouk et al., 2009; Iskratsch et al., 2014;). The first interface between cells and their physical environment is the plasma membrane, focal adhesions and their coupling with the cytoskeleton (Dupont et al., 2011; Wickström et al., 2018; Dupont et al., 2022;), but, in recent years, the nucleus, as well as other intracellular organelles such as mitochondria (Kirby et al., 2018; Helle et al., 2017; Romani et al., 2022), have emerged as additional mechanosensors, with the ability to trigger cellular responses when they are subjected to mechanical constraints. Nuclear mechanosensing relies on modulation of nucleocytoplasmic transport (Elosegui-Artola et al., 2017; Andreu et al., 2022;), chromatin structure (Nava et al., 2020;) and enzymatic activities (Kidiyoor et al., 2020; Enyedi et al., 2016;) and induces short term cellular responses such as contractility (Venturini et al., 2020; Lomakin et al., 2020;), as well as transcriptional reprogramming (Alraies et al., 2024; McCreery et al., 2025; Tajik et al., 2016; Dupont et al., 2022;).

Force transmission from the surface of the cell to the chromatin via the actomyosin cytoskeleton and the LINC complex was proposed and investigated for several decades (Maniotis et al., 1997; Ingber et al., 1997; Wang et al., 2009; Lombardi et al., 2011; Denais et al., 2014; Vahabikashi et al., 2022;). For example, cyclic cell stretching was shown to induce heterochromatinisation of specific loci by acting on the Polycomb complex in an emerin-dependent manner (Le et al., 2016). More recently, it has been shown that nuclear membrane tension is an important player in nuclear mechanosensing. Nuclear envelope tension can be modulated by the classical force transmission from the cell surface, via the cytoskeleton, or directly upon large cell shape changes, in response to osmotic shock or confinement (Nava et al., 2020). Tension can have a direct effect on enzymatic activities (*e.g.,* cPLA2, Enyedi et al., 2016; Venturini et al., 2020; Lomakin et al., 2020; or Kidiyoor et al., 2020;), open nuclear pores (Elosegui-Artola et al., 2017; Andreu et al., 2022;) and induce chromatin remodeling (Nava et al., 2020; Hsia et al., 2022). Nuclear envelope tension, mostly sustained by the lamina, was also proposed to increase intranuclear pressure, eventually leading to the formation of nuclear blebs and envelope ruptures (Denais et al. 2016; Raab et al. 2016; Srivastava et al., 2021;). The balance of forces at the envelope (osmotic versus hydrostatic pressure) can in principle modulate the volume of the nucleus (Kim et al., 2015; Rollin et al., 2023). Changes in nuclear volume would impact chromatin organization, nucleoplasmic crowding (Szórádi et al., 2021) and the concentration and potential phase separation of nuclear proteins, which could all impact gene transcription and thus cellular behavior and fate (Zhao et al., 2024; McCreery et al., 2025).

Although nuclear volume appears an important feature of cell physiology (McCreery et al., 2025), it has not been studied in detail and its potential modulation remains a subject of controversy. Some articles report large changes of volume upon deformation, giving the image of an object with a stiff envelop but a very small bulk osmotic modulus (Rowat et al., 2006, Finan et al., 2009, Stöberl et al., 2024), while others report a constant volume, leading to the conception of an incompressible object, at least in the physiological range of forces relevant for cells (McKee et al., 2025;). In these studies, the methods applied to deform and measure the nuclear volume, the cell types, as well as the extent and the timescale of the deformation, all differ, making it difficult to reach a clear consensus on this important question.

To account for these various observations, several physical models have been proposed, emphasizing the importance of specific nuclear components, leading to different predictions. A classical model for the nucleus represents the lamina as an elastic shell, encompassing a constant volume (Dickinson et al., 2024). Conversely, the nucleus, because of its large pores and its chromatin content, has also been described as a polymer hydrogel losing volume upon deformation like a sponge (Szórádi et al., 2021, Mazumder et al., 2008). More recently, several experimental and theoretical studies, propose that the nuclear volume is conserved by an osmotic equilibrium dominated by nuclear proteins (Deviri et al., 2022; Lemière et al., 2022, Rollin et al., 2023), explaining the long known nuclear/cytoplasmic volume ratio and the experiments showing that nuclear volume depends on import/export rates (*e.g.*, Levy et al., 2010; Neumann et al., 2007; Cantwell et al., 2019; Pennacchio et al., 2024). These studies suggest that the important notion determining volume conservation and the level of envelope tension is the bulk osmotic modulus of the nucleus. In this view, an increase in nuclear envelope tension could balance a difference in osmotic pressure between the nucleus and the cytoplasm, which would correspond to a loss of nuclear volume, an increased concentration of nuclear proteins and thus an increased intranuclear pressure (Rollin et al., 2023). Although attractive and compatible with prior experimental observations such as the formation of nuclear blebs upon large nuclear deformations (Raab et al., 2016; Nader et al., 2021; Baird et al., 2026;), this model has not been tested experimentally so far.

Here we combined controlled nuclear deformation with simultaneous force and volume measures to better understand what dominates nuclear mechanics. By doing so, we validate the mechano-osmotic model proposed by Rollin et al. (Rollin et al., 2023). We first show that the nucleus can deform either at constant volume and zero envelop tension, or loose volume when the envelope gets tensed, which reconciliates seemingly contradictory observations and clarifies the contexts in which nuclear mechanosensing, driven by envelope tension or volume loss, can be triggered. We then introduce a theoretical framework to account for the dynamics of nuclear volume changes. This allows us to account also for the transient regimes and give access to new parameters of nuclear mechanics such as an effective bulk friction. It also reveals that the nucleus changes volume slowly (at the minute scale), similar to a dense hydrogel with a nanometric pore size (Mollenkopf et al., 2025). This suggests that nuclear volume relaxation —and consequently, force relaxation—could limit the migration of fast-moving cells such as leukocyte through dense tissues, since they must be able to translate their entire body through micron-sized constrictions at the minute timescale (Rowat et al., 2013; Lam et al., 2013).

## Results

To systematically investigate how the volume of the nucleus changes upon deformation, we first used a static multi-well confiner device (Le Berre et al., 2014). We imposed a range of controlled uniaxial deformations on rounded HeLa cells platted between non-adhesive parallel surfaces coated with PLL-PEG, with spacers from h=20 µm, serving as non-confined controls down to h=3 µm of height (Fig. 1A). Nuclei of cells expressing EGFP-LAP2b, a nuclear envelope protein, were imaged using confocal microscopy 10-30 minutes post-confinement. Three-dimensional reconstructions show that significant nuclear deformation occurs for confinement heights below 12 µm (Fig. 1B), consistent with a population-averaged homeostatic nuclear volume of 870 µm³, corresponding to an average nuclear diameter of approximately 12 µm (Fig. 1C). Consistent with previous studies, between 20 µm and 12 µm, nuclei were only mildly deformed, resembling crumpled spheres, whereas at lower heights they adopted a flattened, pancake-like morphology (Fig. 1B; Supplementary Fig. 1A; Supplementary Video 1).

**Figure 1:**
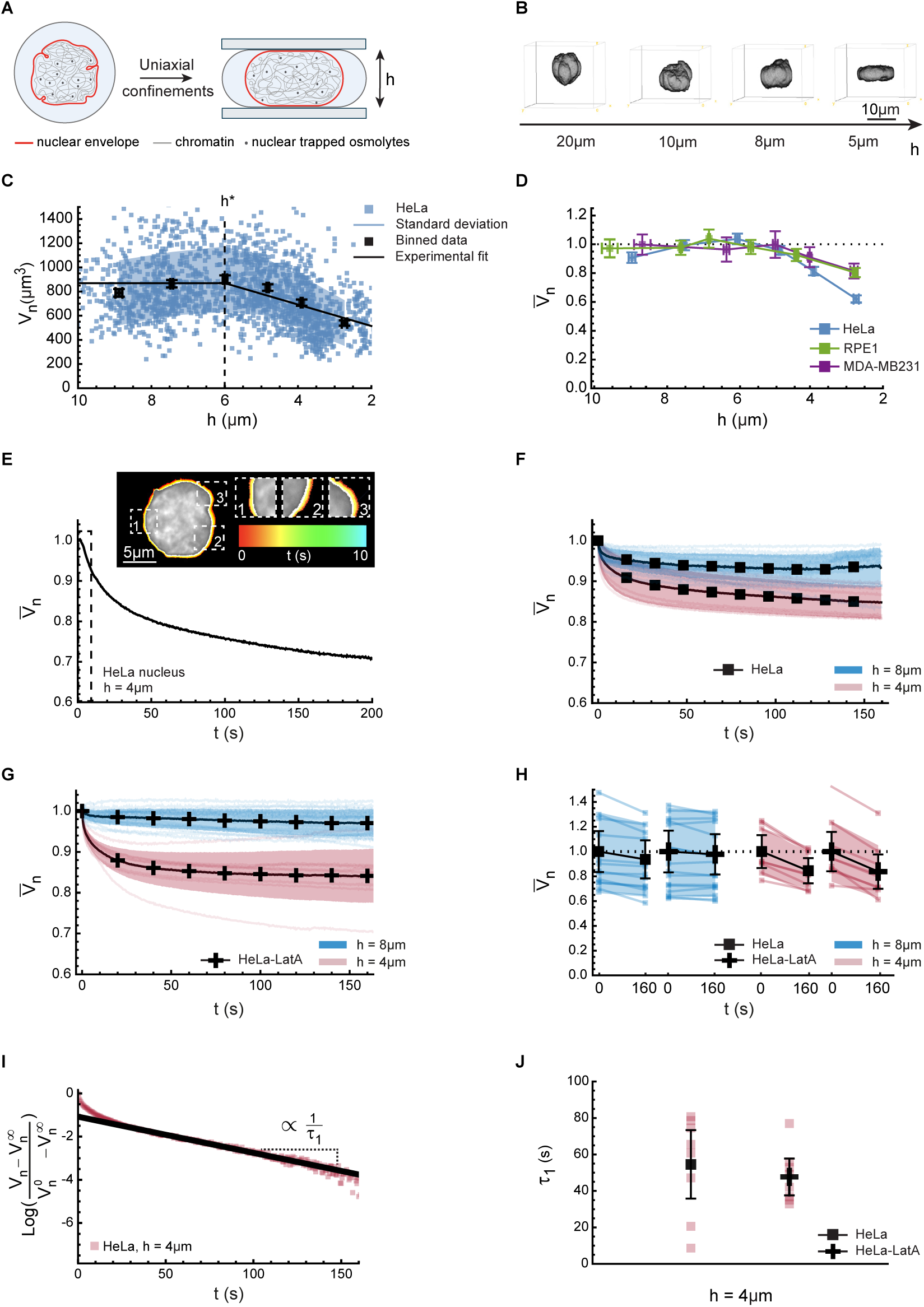
Nuclear volume response to uniaxial confinement on timescales from seconds to minutes. **A.** Schematic of the experimental setup. Cells and their nuclei are confined between two flat surfaces at varying heights (*h*). Key nuclear components are illustrated: nucleoplasm (blue), nuclear envelope (red), chromatin (grey filaments), and trapped nuclear osmolytes, such as proteins (grey dots). Further details are provided in Supplementary Information Section III.G. **B.** 3D confocal reconstructions of representative Hoechst-stained HeLa nuclei at varying confinement heights *h* (see Supplementary Video 1). **C.** Nuclear volume (V*_n_*) of individual HeLa cells (blue squares) as a function of confinement height (*h*). Nuclear volume loss is observed below a threshold confinement height (ℎ^∗^) which falls within the *5-6 µm* range. Data are binned into intervals of equal sample size (*N* = 7 independent experiments, *n* = 1,654 cells; 275 cells per bin), with black squares denoting the binned mean ± 95% confidence interval (CI). Error bars are smaller than the symbol size. A piecewise linear fit (black line) is provided as a visual guide, with the standard deviation indicated in light blue. Volume measurement details are provided in Supplementary Information Section III.I. **D.** Normalized nuclear volume relative to the unconfined state across three cell lines. Squares and whiskers denote mean ± 95% CI. Data are binned into intervals of equal sample size for HeLa (*N* = 7 independent experiments, *n* = 1,654 cells; 275 cells per bin), RPE1 (*N* = 4, *n* = 1,233; 205 per bin), and MDA-MB231 (*N* = 5, *n* = 550; 110 per bin). **E.** Temporal evolution of normalized nuclear volume (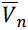) for a representative HeLa cell following a 4 μm confinement step using a dynamic confiner (Supplementary Information Section III.G). The nuclear envelope is visualized via EGFP-LAP2β expression, imaged at 2 frames per second. Inset: Color-coded temporal overlay of confocal images for the initial 10 s post-confinement, featuring magnified regions of the nuclear envelope (corresponding to the dotted box). Time *t* = 0 corresponds to the first frame recorded post-confinement. **F,G.** Dynamics of normalized nuclear volume (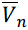) for single cells following confinement at 4 μm (light red) and 8 μm (light blue) using a dynamic confiner. Data are shown for control HeLa cells (**F**; *N* = 2 independent experiments, *n* = 8 cells at 4 μm, *n* = 14 at 8 μm) and cells treated with Latrunculin A (LatA; 2 μM in DMSO, added 20 min before the experiment;) (**G;** *N* = 2, *n* = 9 at 4 μm, *n* = 17 at 8 μm). The nuclear envelope was visualized via EGFP-LAP2β expression (acquired at 2 frames per second). Individual cell traces are plotted in their respective colors, while solid black lines and shaded regions indicate the mean and standard deviation. Volumes are normalized to the first acquired time point, and time *t* = 0 is referenced to the first frame immediately following confinement. **H.** Normalized nuclear volume (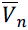) of single nuclei evaluated immediately post-confinement (*t* = 0) and at the end of the recording (*t*^∞^= 160s) for the same cell. Data are derived from the same cells analyzed in **F** and **G**, comparing control HeLa cells (black squares) and LatA-treated cells (black crosses) confined at 8 μm (light blue) and 4 μm (light red). Black symbols denote the mean ± 95% CI. Light-colored lines represent individual nuclei traces, and shaded regions indicate the standard deviation. Volumes of individual nuclei are normalized to the population’s mean nuclear volume at *t* = 0. **I.** Log-linear time course of the nuclear volume (V*_n_*) for a single, representative HeLa nucleus confined at 4 μm (dataset from **F**–**H**). Red dots represent individual time points. The y-axis 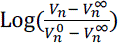 displays the logarithm of the relative nuclear volume change between *t* = 0 and *t*^∞^= 160s. The solid black line indicates a linear regression fit from 20 s to 150 s post-confinement, used to extract the characteristic timescale (τ1) of the slow, mono-exponential volume loss kinetics. **J.** Characteristic timescales (τ1) of nuclear volume loss for individual HeLa cells under 4 μm confinement, extracted using the method described in **I**. The plot compares control HeLa cells (squares; *n* = 8 cells) and LatA-treated cells (crosses; *n* = 9 cells), utilizing the same dataset as in **F**–**I** (representing *N* = 2 independent experiments per condition). Black symbols denote the mean ± 95% CI, with red squares indicating individual data points.

## A threshold behavior in nuclear volume response to confinement

We next measured nuclear volume from our confocal images at different confinement heights (Supplementary Fig. 1B; Supplementary information, Section III.I). We observed a threshold behavior: down to approximately 6 µm in height, i.e. ∼50% of vertical strains, nuclei maintained a nearly constant average volume at the population level. Conversely, below 6 µm, the average nuclear volume decreased, with up to 40% volume loss observed at 3 µm confinement (Fig. 1C). To assess the generality of this threshold behavior, we repeated the experiment on RPE1 and MDA-MB-231 cell lines (Supplementary Fig. 1C,D). All tested cell lines showed similar trends, with threshold heights for volume loss in the ∼5–6 µm range and nuclear volume loss of up to ∼20% (Fig. 1D). While this threshold behavior upon uniaxial confinement is broadly conserved, the precise height at which volume loss occurs is not universal; exact values vary slightly across cell lines and remain subject to experimental approximation. We also emphasize that this conserved threshold behavior exists despite nuclear volumes that differ substantially amongst different cell lines ∼870µm³ (HeLa), 630µm³ (RPE1), and 400µm³ (MDA), corresponding to diameters of ∼12, ∼11 and ∼9 µm.

Because the previous population-level data showed high variability — partly due to asynchronous cells distributed throughout the cell division cycle — we tracked the volume changes of individual nuclei. We used a dynamic confiner device (Le Berre et al., 2014) to impose confinement on single HeLa cell nuclei, reducing the height from unconfined to either 8 µm or 4 µm. Confocal images were acquired at high frame rate (∼500 ms) to capture the nuclear volume dynamics for several minutes following confinement (Fig. 1E-J). Consistent with our population-level measurements, there was no detectable volume change after a confinement at 8 µm, but a volume loss of ∼15% at 4 µm confinement (Fig. 1H). Importantly, these measurements allowed us to observe the dynamics of nuclear volume loss. We found that the typical timescale of nuclear volume loss was ∼50s (Fig. 1I,J), while the confinement itself occurred in less than a second. This suggests that nuclei lost a negligible volume during the confinement step —consistent with our model (detailed below), which predicts a reduction of ∼1%— but rather relaxed to a lower volume in the minutes following confinement.

Together these results show that nuclear volume follows two distinct regimes in response to deformation. Above a cell-type-specific threshold height, nuclei maintain a constant volume despite compression. Below this threshold, they lose up to ∼30% of their volume over a characteristic timescale of ∼50 s (in HeLa cells), corresponding to an average volume efflux rate of ∼100 µm³/min.

Importantly, we found that nuclei begin to lose volume within a range of confinement heights (6 to 4 µm) where nuclear envelope (NE) rupture and blebbing remain negligible (Supplementary Fig. 1E,F), ruling out a role of NE ruptures. In addition, depolymerizing the actin cytoskeleton with Latrunculin A did not change the timescale nor the amount of volume loss compared to control conditions (Fig. 1H,J), ruling out a role of processes activated by tension in the cell cortex upon deformation (*e.g.*, stretch activated ion channels or other volume regulatory mechanisms that depend on the actin cytoskeleton, Venkova et al., 2022), and suggesting a mechanism more directly related to nuclear deformation itself.

## Theoretical framework and validation: A quasi-static mechano-osmotic model for nuclear volume regulation

The biophysical regulation of nuclear volume has attracted growing theoretical and experimental interest in recent years (*e.g.*, Finan et al., 2009; Kim et al., 2015; Deviri et al., 2022; Lemière et al., 2022; Rollin et al., 2023). Increasing evidence supports that osmotic forces govern nuclear volume over minute timescales, primarily through the requirement that the mechanical pressure difference across the nuclear envelope is balanced by the osmotic pressure difference, and some of the existing models would predict a loss of volume upon compression. Building on these insights, we adapted existing models—including the mechano-osmotic framework from our previous work (Rollin et al., 2023;)—to explain the two regimes of volume loss upon confinement (Fig. 1C). Our model relies on two key assumptions. First, inspired by studies on hypo-osmotic shocks in chondrocyte nuclei (Finan et al., 2009;) and compression experiments on cultured cells (Lomakin et al., 2019; Venturini et al., 2019;), we propose that nuclear envelope tension exhibits a threshold behavior, remaining negligible until the nuclear surface area reaches a critical value *S*^∗^(Supplementary information, Sec. I.B). This is further supported by observations that nuclear wrinkles form and unfold at constant volume during cell rounding and spreading (McKee et al., 2025;), consistent with a deformation regime where the envelope remains tensionless. Second, unlike previous models of cell volume regulation that neglect the role of Laplace pressure at the cell cortex, we propose that Laplace pressure at the nuclear envelope plays a critical role due to the high permeability of the nuclear envelope (via nuclear pores) to small osmolytes, that strongly reduces the osmotic modulus of the nucleus. To capture this, we distinguish between two classes of osmolytes: (1) impermeant (“trapped”) osmolytes, which are confined within the nucleus and cannot freely diffuse through nuclear pores, though they are mobile within the nuclear compartment. These include large macromolecules such as large proteins, as well as non-condensed DNA counterions required to maintain electroneutrality. (2) permeant (“non-trapped”) osmolytes, which can freely diffuse across the nuclear envelope but are retained within the cell (at least on short timescales and in the absence of membrane ruptures, see Venkova et al for a discussion of cell volume changes upon large deformations). These include smaller molecules such as ions, amino acids and small proteins (below ∼50 kD for human cells – the relative composition of the nucleus and cytoplasm and the resulting osmotic pressures have been discussed in several studies e.g. Biswas et al., 2025; Rob Philipps et al., 2015;). For simplicity, and given the limited knowledge regarding the electric potential across the nuclear envelope, we neglect electrostatic effects in our model. Consequently, the balance of water chemical potential over timescale of minutes dictates the equilibration of concentrations of freely diffusible solutes between the nucleoplasm and cytoplasm.

To test our hypothesis that tension build-up drives nuclear volume loss, we explicitly modeled nuclear geometry and its deformation under compression (Fig. 2A; Supplementary Fig. 2A). We predict nuclear shape under compression by solving a mechanical force balance at the nuclear envelope, in the spirit of previous work (Kim et al., 2015). For simplicity and to isolate the role of the nuclear envelope, our model considers only the contribution of the nuclear envelope tension in this balance and does not include potential mechanical forces from the cytoplasm or the chromatin (see Discussion for further considerations and Supplementary information, Section II. B). The predicted shapes range from nearly spherical geometries under weak confinement to flattened “pancake-like” configurations under strong confinement. We verified that the model qualitatively captures the diversity of nuclear shapes observed experimentally (Fig. 2B; Supplementary Fig. 2B). Notably, in the regime where nuclear volume loss occurs (height < 6 µm), nuclei are well described by the pancake-like geometry predicted by the model (Supplementary Fig. 2C-E; Supplementary information, Section III. I).

**Figure 2:**
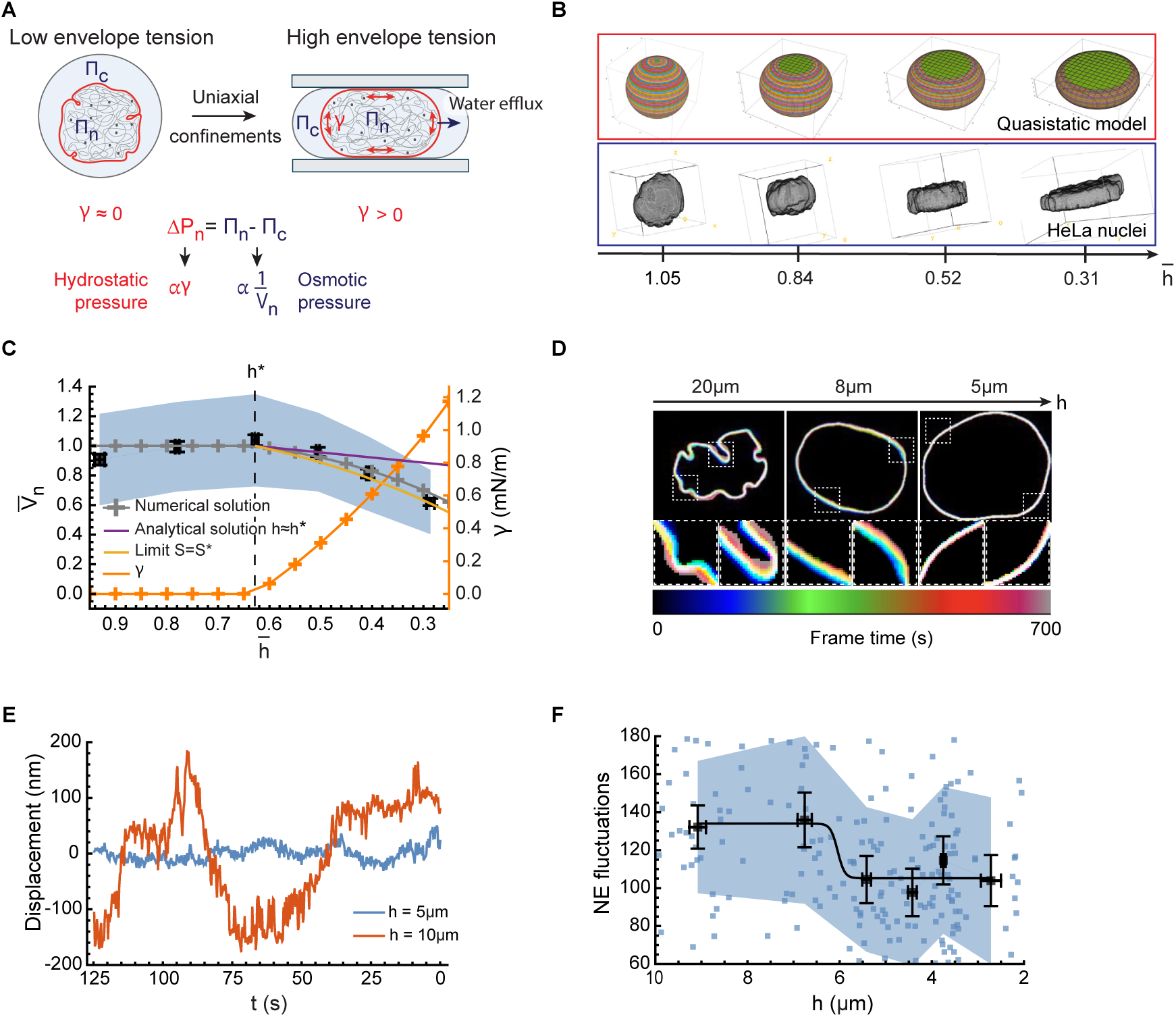
The quasistatic mechano-osmotic model of the nucleus reproduces the threshold behavior of nuclear volume under uniaxial confinement on the minute timescale. **A.** Schematic of the quasistatic model of nuclear volume loss upon confinement (nucleoplasm, blue; nuclear envelope, red; chromatin, grey filaments; trapped nuclear osmolytes such as proteins, grey dots). The equation describes the balance of hydrostatic and osmotic pressure difference (respectively Δ*P_n_* and Π*_n_* – Π*_c_*) at the nuclear envelope, establishing the relationship between nuclear envelope tension (γ) and nuclear volume (*V_n_*) (Supplementary Information Section I). **B,C.** A tension-driven nuclear volume loss model quantitatively reproduces the nuclear response to confinement. **B.** Qualitative comparison between the model-predicted nuclear geometry under confinement (Supplementary Information Section I.C) and 3D reconstructions of representative Hoechst-stained HeLa cell nuclei. **C.** Population-averaged HeLa nuclear volume as a function of confinement height (data from Fig. 1c). Solid lines represent model predictions using parameters independently sourced from the literature (Supplementary Information Section I.G). The limit *S* = *S*^∗^, corresponds to deformation under constant area, and the limit ℎ ≈ ℎ^∗^, represents the linear response regime at the onset of tension γ build-up (orange line; Supplementary Information Section I.D). **D, E, F.** Nuclear envelope fluctuations decrease below a height threshold (see Supplementary Video 2). **D.** Representative overlay of time-lapse confocal images of the EGFP-LAP2β-stained nuclear envelope at varying confinement heights. **E.** Examples of nuclear envelope displacement trajectories used to calculate the fluctuations. **F.** Quantification of the nuclear envelope fluctuation amplitude in HeLa cells under confinement (*N* = 3 independent experiments, *n* = 221 cells; 37 cells per bin; Supplementary Information Section III.H). Colored squares indicate individual data points; black squares and whiskers represent the mean ± 95% CI, respectively, and the shaded region denotes the standard deviation.

To determine the parameters controlling the amount of nuclear volume loss to confinement in our model, we solved the non-linear model perturbatively close to the threshold where the envelope becomes taut (Supplementary information, Section I.D). Our analysis revealed that the amount of volume loss close to the threshold depends on the initial osmotic pressure exerted by trapped nuclear osmolytes, Π_*t*_, divided by a mechanical pressure 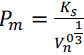, that is proportional to the stiffness of the nuclear envelope *K*_s_, scaled by a nucleus length scale 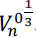. In other terms, if Π*_t_*<*P_m_*, the model predicts that the stretching of the nuclear envelope is not sufficient to press water out of the nucleus at the threshold; water efflux only begins when confinement reaches the length scale 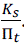. By contrast, when Π*_t_*<*P_m_*, the nucleus undergoes volume loss immediately beyond the threshold at nearly constant surface area, corresponding to the stiff-surface regime.

We next compared the experimentally measured slope of nuclear volume loss near the deformation threshold with predictions from our model. To quantitatively match this slope, the model must operate in a regime where volume loss occurs at approximately constant surface area after relaxation —consistent with a stiff nuclear envelope and a low bulk modulus. We compared this near-threshold prediction with numerical solutions of the full model over the entire range of observed deformations (Fig. 2C). Because the nuclear volume data alone do not meaningfully constrain the fit, we fixed all model parameters based on literature measurements (Supplementary information, Section I.G). Using this parameter set, the model accurately reproduces the magnitude of the observed volume loss, lending support to our hypothesis.

Altogether, our model quantitatively reproduces the experimental observations through a mechanism of nuclear volume loss driven by tension increase at the nuclear envelope, under two key conditions: (1) the nuclear envelope tension should exhibit a threshold like behavior, consistent with prior work (Finan et al., 2009), and (2) the osmotic pressure from trapped nuclear osmolytes remains small with respect to the mechanical pressure 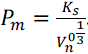. Under these two conditions, a key feature of our model is that the onset of nuclear envelope tension should coincide with the threshold for nuclear volume loss and that the bulk modulus of the nucleus should be in the kPa range (Supplementary Fig. 2F; Supplementary information, Section 1.E)., which is of the same order of magnitude than previous estimates in other systems (Biswas et al., 2025;).

## Quantifying nuclear envelope tension

We next sought to experimentally test and validate our theoretical predictions. To obtain a qualitative proxy for nuclear envelope (NE) tension, we first quantified NE fluctuations by measuring deviations of the envelope from its mean position. We observed that NE fluctuations are reduced when cells are confined below 6 μm (Fig. 2D-F; Supplementary Video 2), which coincides with the onset of nuclear volume loss, consistent with our model. Next, to quantitatively estimate nuclear envelope tension, we adapted the method developed by Fischer-Friedrich *et al*. (Fischer-Friedrich et al., 2014), to measure the force exerted by the cell on a flat wedged cantilever (Fig. 3A). We applied successive confinement steps to single cells at a constant cantilever descent velocity of 0.5 μm/s. After each step, the confinement height was held constant for 250 s to allow the nucleus to relax before proceeding to the next increment. At the end of each relaxation period, we acquired an image of the nuclear midplane, enabling simultaneous estimation of both nuclear volume and the exerted force following relaxation (Fig. 3B-D).

**Figure 3:**
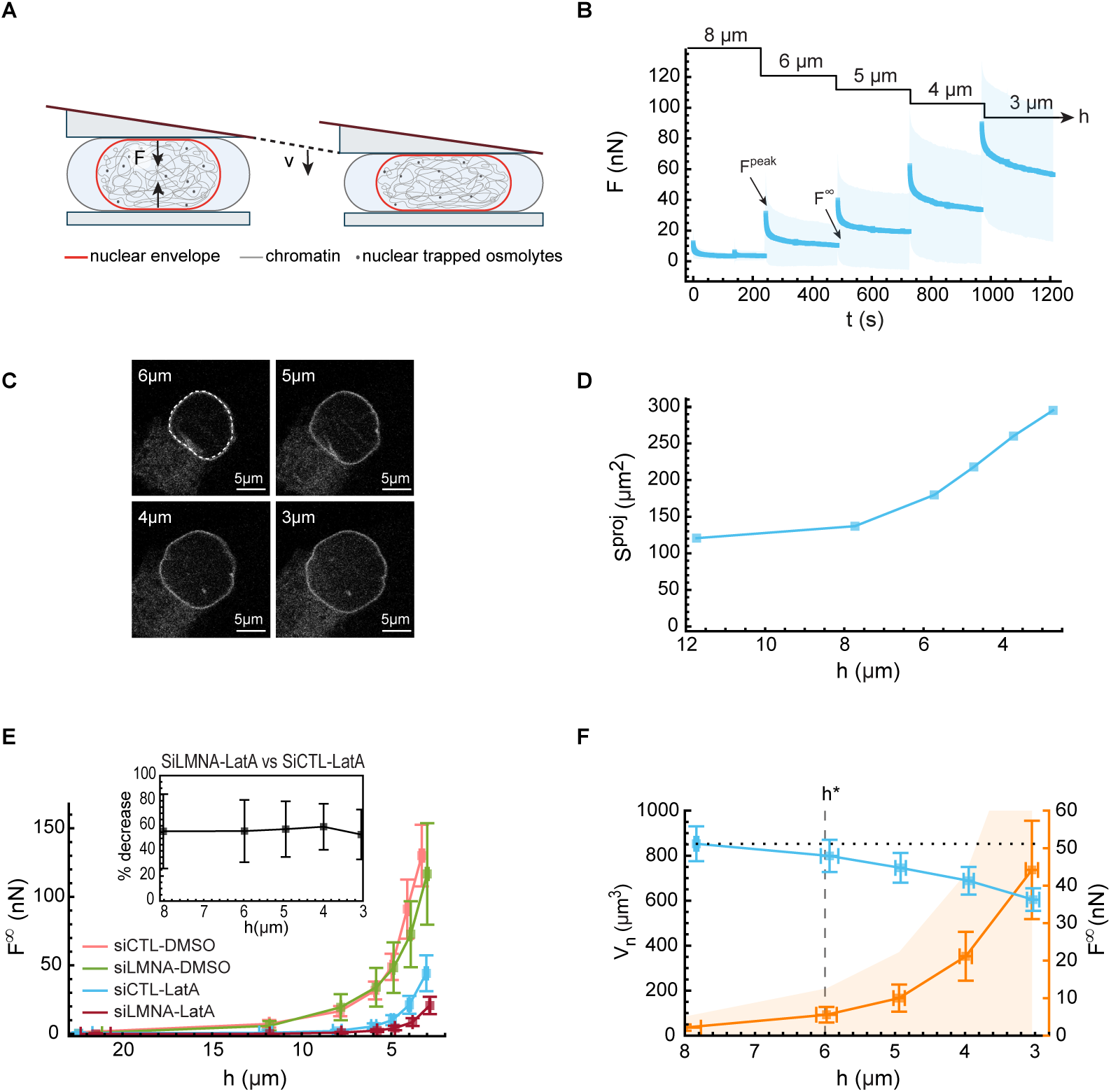
Increased nuclear envelope tension drives minute-scale nuclear volume loss. **A-D.** A combined AFM and confocal microscopy experimental setup to characterize single-nucleus mechanics across variable timescales and deformation strengths. **A.** Schematic of the experimental setup. Nuclei are uniaxially compressed by a wedge cantilever at a constant velocity *v*, allowing for precise control of both confinement height and deformation rate. **B.** Averaged force response of control siRNA Latrunculin A treated Hela nuclei (siCTL-LATA; *N=10*, independent experiments, *n=45* cells) during successive confinement increments. Nuclei were compressed at a constant velocity of *v* = 0.5 μ*m*. *s*^-1^to target heights (ℎ = 8,6,5,4 and 3μm), with 250 s relaxation periods between steps. Forces at the end of the compression increments and relaxation periods are denoted *F*^peak^ and *F*^∞^. The blue line and shaded region represent the mean and standard deviation, respectively. **C.** Representative confocal images of an EGFP-LAP2β-stained siCTL-LatA HeLa cell nucleus, acquired at the end of each relaxation period (confinement heights shown: ℎ = 6,5,4 and 3μm). **D.** Nuclear midplane area measurements derived from the images in **C** enable the estimation of nuclear volume at the end of each confinement step. **E.** Post-relaxation force (*F*^∞^, defined in Fig. 3b) across varying confinement heights in HeLa cells subjected to four perturbations: siCTL-DMSO (*N*=3, *n*=26), Lamin A/C knockdown DMSO (siLMNA-DMSO) (*N=4, n=15*), siCTL-LatA (*N*=10, *n*=45), and siLMNA-LatA (*N*=7, *n*=32). Squares and whiskers denote the mean ± 95% CI. **Inset**. Percentage of force reduction in siLMNA relative to siCTL cells (both LatA-treated). Uncertainty was calculated via error propagation: 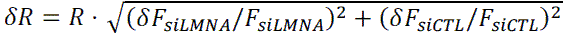, where *R* is the force ratio and *F* denotes the mean post-relaxation force. **F.** Post-relaxation nuclear volume and force for siCTL-LatA HeLa cells across varying confinement heights (*N=10, n=45*). Colored squares and whiskers represent the mean ± 95% CI; shaded regions denote the standard deviation. Note that the height thresholds ℎ^∗^for force increase and nuclear volume loss coincide.

We first performed biochemical perturbations of the actin cortex to distinguish the respective contributions of the nucleus and the cell to the observed force response curves (Fig. 3E; Supplementary Fig. 3A, A-H; Supplementary Fig. 3B, A-B). Inhibiting myosin contractility with Y-27632 reduced both the peak force (∼25%) and post-relaxation force (∼50%). Actin depolymerization induced by Latrunculin A (LatA) did not further reduce either the peak force or the post-relaxation force relative to non-contractile HeLa cells. In contrast, Lamin A knockout led to a further 60% reduction in the post-relaxation force (Fig. 3E, inset). Together, these results demonstrate that half of the force response is due to the cell surface tension, driven by myosin contractility, with nuclear mechanics dominating the residual force following myosin inhibition. Thus, to definitively exclude any residual mechanical contribution from the actin network itself, we treated cells with LatA, which altered neither the extent nor the kinetics of nuclear volume loss (Fig. 1H,J).

The force response of LatA-treated HeLa cells exhibited a threshold-like behavior, with almost no force down to 6μm and a gradual increase in the force needed to deform the cells below that height. Because this force, in LatA treated cells, as demonstrated with the Lamin A KD experiments, is mostly due to nuclear pressure, which is itself related to nuclear envelope tension via Laplace’s law (Supplementary information, Section I. B and discussion on the role of chromatin), these results confirm that nuclear envelope tension rises below 6 μm. We further used the projected nuclear surface area, measured at the end of each relaxation phase, to estimate nuclear volume based on the geometrical model (Supplementary information, Section III.I). This showed that volume loss follows the increase in nuclear envelope tension (Fig. 3F), consistent with our model prediction.

## Nuclear envelope folds: A reservoir of excess surface area and their role in mechanical adaptation to confinement

We next asked what mechanism could explain the existence of a regime of deformation with neither volume loss nor nuclear-envelope tension. Based on our and other’s observations (Lomakin et al., 2019; Venturini et al., 2019; McKee et al., 2025;), we propose that the NE possesses an excess of surface area stored into large folds, that can be opened upon compressive forces of a few nanonewtons. Multiple studies have reported the presence of nuclear envelope folds in various biological contexts: *in vivo* in the Drosophila embryo (Jackson et al., 2022), during hematopoiesis (Biedzinski et al., 2020), and *in vitro* in round, unspread cells (Lomakin et al., 2019;). Here, we also observe folded nuclear envelopes in round cells that are not subjected to external forces (Fig. 1B; Supplementary Fig. 1A; Supplementary Video 1 & 2). Electron microscopy of HeLa cell nuclei revealed that the nuclear envelope within folds contains nuclear pore complexes (NPCs) and does not exhibit any striking structural differences compared to the smooth regions of the envelope (see Supplementary Fig. 4A). This suggests that there is no force bearing structures holding the envelop folded and that the folds are just the result of an excess area of the envelope, resulting in buckling, explaining why they can open at low force.

Using population-level projected surface area measurements from the multiwell confiner experiment, we observed that the nuclear envelope progressively smoothens with increasing confinement in the HeLa cell line (Fig. 4A; Supplementary Fig. 1A; Supplementary Video 2). We confirmed this at the single-nucleus level using a dynamic confiner device, imaging the nuclear envelope immediately before and after confinement (Supplementary Fig. 4B). To systematically quantify the unfolding, we first performed a semi-manual segmentation of the nuclear envelope using thresholding and developed specific metrics to assess fold abundance in these segmented images (Supplementary, “Nuclear shape” Methods section and Fig. 4A & Supplementary Fig. 4C). This analysis revealed a significant reduction in nuclear envelope folds upon confinement — indicative of fold opening — with a characteristic inflection point at 6 μm of confinement, closely matching the threshold for nuclear volume loss. To further determine whether these folds could be opened not only by external constraints but also by internal active forces, we seeded HeLa cells on micropatterns of increasing area to increase cell spreading and thus, consequently, the actomyosin-mediated forces applied on the nucleus. We found that the extent of NE folding decreased as spreading area increased (Supplementary Fig. 4D,E, consistent with McKee et al., 2025), indicating that NE unfolding is a general response to both externally imposed and cell-generated mechanical stresses.

**Figure 4:**
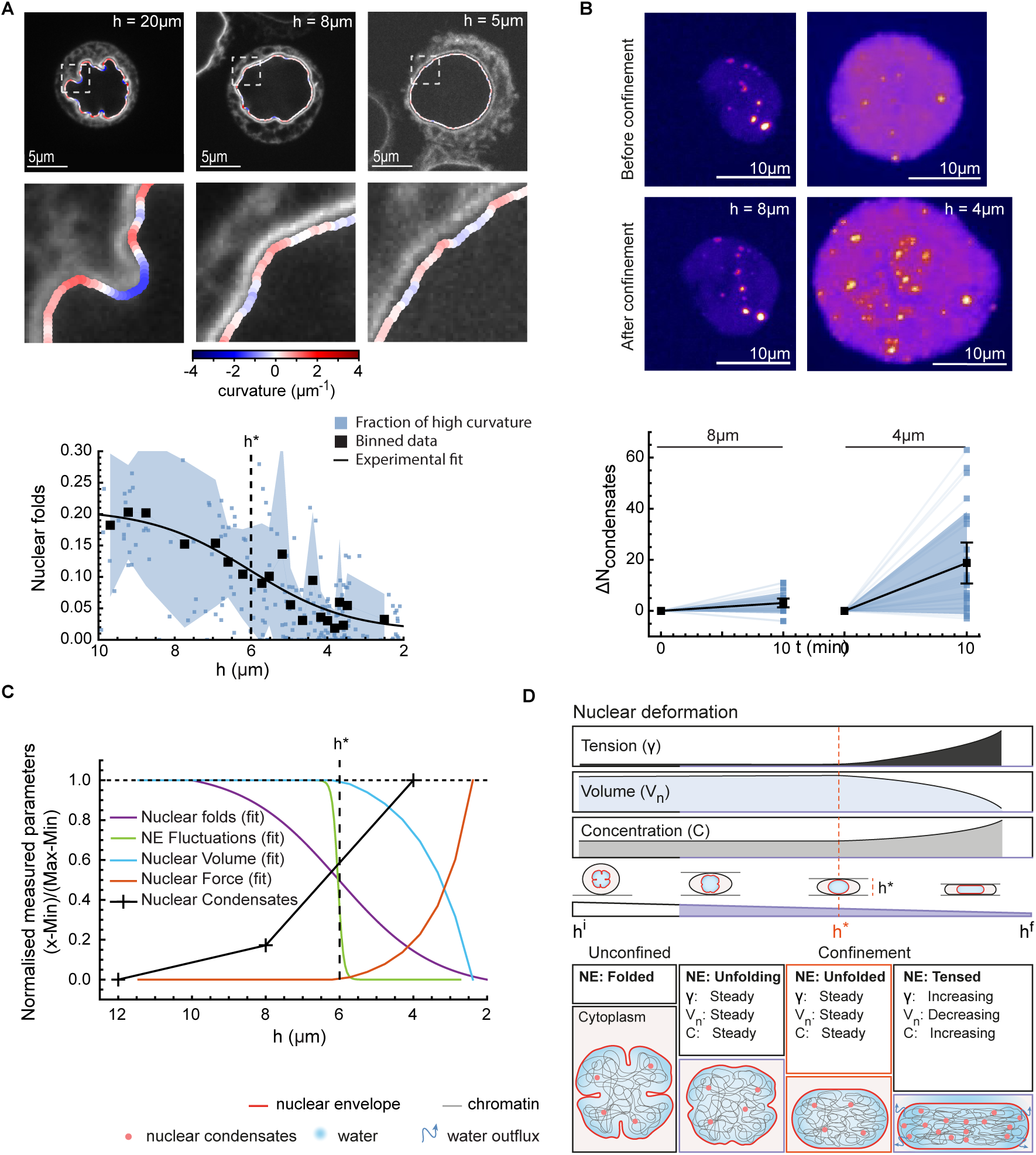
Nuclear envelope folds buUer tension until full unfolding triggers increased intranuclear pressure, volume loss, and increased trapped osmolyte concentration. **A.** Nuclear envelope smooths upon uniaxial confinement (see Supplementary Video 2). Representative EGFP-LAP2β-stained nuclear envelope shapes at varying confinement, overlaid with local curvature color-coded from negative (blue) to positive (red). Quantification of the “fraction of high curvature” metric (*N=3 independent experiments, n=285 nuclei;* Supplementary information, Section. III.J; Supplementary Fig. 4c). Colored squares represent individual data points; black squares denote binned data (*n=20* cells per bin). The shaded region represents the standard deviation. The solid line represents the sigmoidal least-squares fit. The inflection point of the sigmoid fit corresponds to the height threshold ℎ^∗^. **B.** Nuclear condensate abundance increases sharply below the nuclear volume-loss threshold (ℎ^∗^). Representative confocal images of Daxx-emGFP-expressing HeLa cell nuclei before and 10 minutes after compression to 8 μm and 4 μm. Quantification of the change in nuclear condensate number per nucleus 10 min post-confinement relative to baseline at 8 μm (*N=3* independent experiments, *n =22* nuclei) and 4 μm (*N=10, n =25*). Colored squares indicate individual nuclei. Black squares and whiskers denote the mean ± 95% CI. HeLa cells expressing Daxx-emGFP were previously described (Garcia-Jove Navarro et al., 2019). **C.** Overlay of normalized post-relaxation metrics across varying confinement heights. All metrics are rescaled to a range of 0–1 for comparison. Data are fitted via least-squares regression using either sigmoidal functions (nuclear folds and NE fluctuations) or the quasistatic model (nuclear volume and nuclear force). **D.** Schematic illustrating the two-regime response of the nucleus to uniaxial confinement at the minute timescale. An initial tension-buéered regime preserves constant volume, while the post-threshold regime—characterized by elevated tension, increased intranuclear pressure, interior concentration, and volume loss—occurs at a quasi-constant nuclear surface area.

Beyond this unfolding regime, our model interprets the volume loss regime as the result of a balance between an increased envelope tension and an increased intranuclear pressure of osmotic origin (Fig. 2A). Consistent with the timescale of volume decrease (minute), which is too short for a significant import/export of proteins, a direct prediction of our model would be that the increase in nuclear pressure corresponds to an increase in macromolecular concentration within the nucleus, that occurs specifically below the confinement threshold for the volume loss regime. To test this prediction, we used HeLa cells expressing Daxx-emGFP (Garcia-Jove Navarro et al., 2019). Cells were confined using a dynamic confiner at either 8 μm, a height in the constant volume regime, and 4 μm, a height inducing strong nuclear volume loss and negligible nuclear envelope rupture (Supplementary Fig.1F). Nuclei were then imaged at high temporal resolution to probe nuclear condensates number and size. We found that the number of nuclear condensates sharply increase at 4 μm of confinement both at the population level and single nucleus level, while it did not change at 8 μm confinement (Fig. 4B & Supplementary Fig. 4F,G). Importantly, consistent with recent reports of condensation during confined migration (Zhao et al., 2024), we found that while the number of condensates increased, their average size did not change (Supplementary Fig. 4H). This excludes fragmentation of pre-existing condensates and instead supports a mechanism in which new condensates nucleate in response to increasing macromolecular concentration.

Taken together, these results indicate that nuclei possess an excess of nuclear envelope (NE) surface area stored in folds when not subjected to mechanical stress. This excess surface can be mobilized during deformation induced either externally or internally, providing a range of ‘safe’ deformation at low tension and constant volume, consistent with reports from us and others (Lomakin et al., 2019; Venturini et al., 2019; McKee et al., 2025). Once nuclear deformation exceeds the threshold of full envelope unfolding, the nucleus enters a regime characterized by a concomitant increase in nuclear envelope tension, intranuclear pressure, trapped osmolyte concentration, and volume loss (Fig. 4C,D; Graphical Abstract).

## A poroelastic model of the dynamics of nuclear volume relaxation upon uniaxial confinement

The simple steady-state model we propose explains well the two regimes of deformation, but in the regime of volume loss, it does not capture the minute-long transient regime during which the nucleus loses volume (Fig. 1F,G). Prior models of nuclear volume regulation rely on the hypothesis that water has equilibrated (no water flows), which breaks down on second timescales (Deviri et al., 2022; Lemière et al., 2022, Rollin et al., 2023), or attribute the relaxation dynamics solely to the nucleoplasmic flow through the nuclear envelope (Kim et al., 2015; Hoffmann et al. 2025;), which fails to account for the multi-timescale relaxation occurring within the first tens of seconds (Fig. 1I). Thus, to understand this transient regime and the potentially new mechanisms that govern it, we developed a time-dependent model of nuclear volume changes tailored to the successive confinement steps applied in AFM experiments and the simple geometry imposed by the wedged cantilever at low confinement heights (Fig. 5A-C). We hypothesized that, on short timescales, the bulk of the nucleus behaves as a porous gel—a concept that has been shown for cells (Moeendarbary et al., 2013) and theoretically proposed for the nucleus (Mazumder et al., 2008).

**Figure 5:**
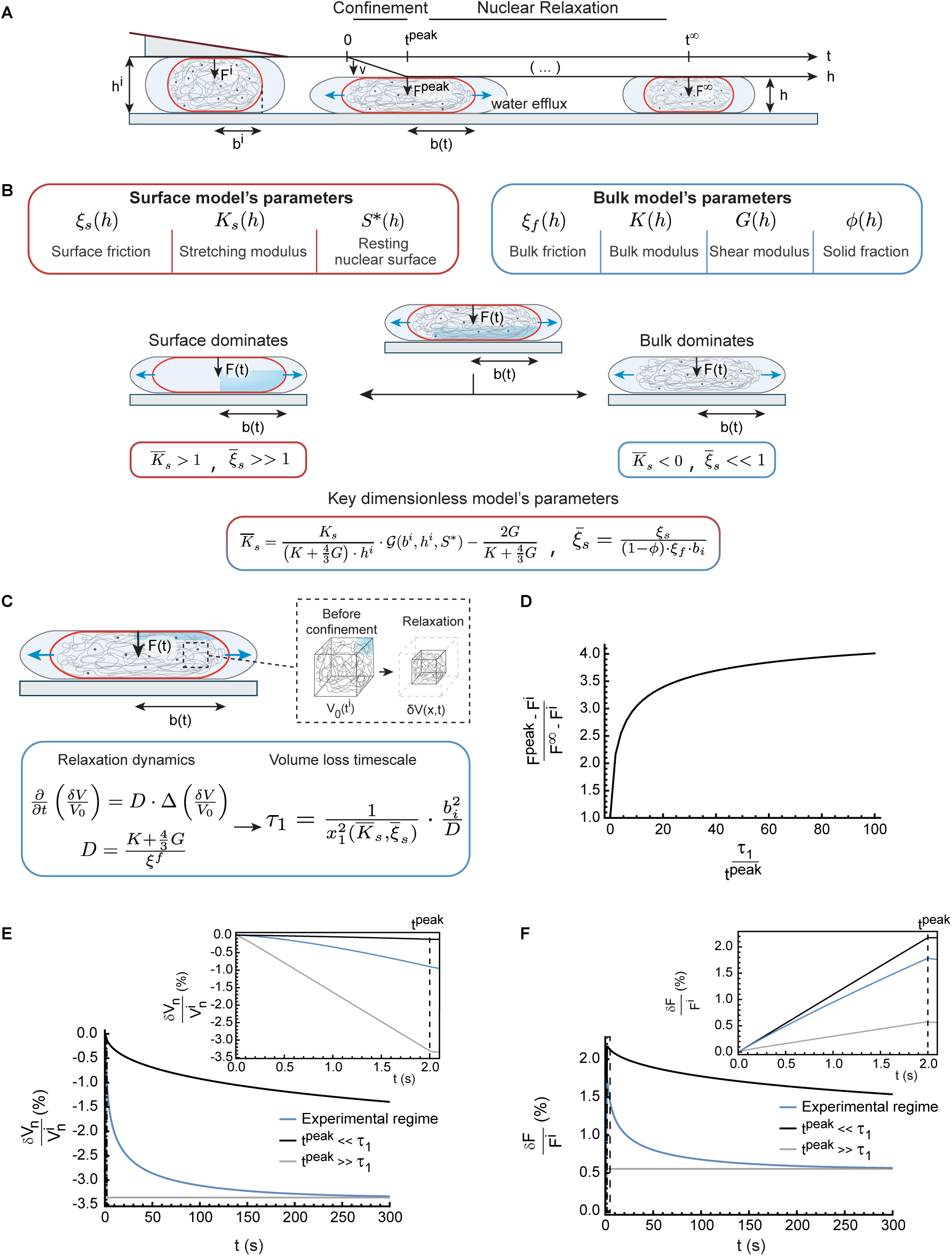
Phenomenological model of nuclear volume dynamics under uniaxial compression. **A.** Schematic of a single confinement step. A cell nucleus, initially confined at a height ℎ*^i^* by an applied force *F^i^* via a wedge cantilever, features an initial midplane radius *b^i^*. The nucleus is subsequently compressed at a constant velocity *v* for a duration of *t^peak^*, followed by a relaxation phase to allow the nucleus to reach its quasistatic volume. The time-dependent evolutions of the midplane radius and the compressive force are characterized by the parameters *b*(*t*)and *F*(*t*), respectively, where *F^peak^* denotes the peak force at the end of the compression step and *F*^9^ represents the asymptotic force after relaxation. The dynamical model predicts the evolving nuclear geometry (and by extension, its volume) and force response based on the intrinsic mechanical parameters of the nucleus. **B.** Nuclear model parameters governing nuclear volume and force dynamics upon uniaxial confinement. Parameters corresponding to the nuclear envelope and nuclear bulk are indicated in red and blue, respectively. The dynamics are controlled by specific combinations of these parameters forming dimensionless quantities 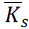, 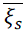, which scale the relative mechanical contributions of the envelope versus the bulk (Supplementary Information Section II). The envelope dominates the response in the limit 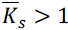 and 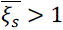, whereas bulk mechanics dominate in the opposite limit. **C.** Poroelastic basis of nuclear volume kinetics. The transient volume change is governed by a diéusion equation whose diéusion constant scales as the ratio of nuclear elastic parameters to bulk friction (Supplementary Information Section II.4.b). This process yields a spectrum of relaxation timescales τ*_n_* (Supplementary Fig. 5B), with τ_1_indicating the slowest mode. Structurally, these timescales scale with the square of the characteristic nuclear length scale *b^i^* divided by the diéusion coeéicient 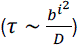, with a scaling factor dictated by the dimensionless parameters 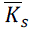, 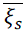. **D.** Nuclear strain-rate stiéening. Dynamic model predictions for the relative peak force **F*^peak^* during a 5 to 4 μm confinement step. The relative force response is evaluated as a function of the ratio of the nuclear volume loss τ_3_to the characteristic compression timescale *t^peak^*. Complete model parameters are detailed in Supplementary Information Section II.G.5, Table III. **E, F.** Dynamic model predictions for the relative nuclear volume loss and force response during a 5–4 μm confinement step across diéerent values of 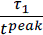. **Top panels:** Magnified views of the dynamics during the compression phase (duration *t*^peak^ = 2*s*). The experimental condition (blue curve) closely matches the regime characterized by negligible volume loss during compression (black curve, less than 1% volume loss), where the corresponding force prediction error remains below 1% (see Supplementary Fig. 5A). **Bottom panels:** The identical volume and force dynamics displayed over an extended 300 s timeline. Model parameters are detailed in Supplementary Information Section II.G.5, Table III.

We extended the stress-diffusion coupling model for poroelastic materials (Tanaka et al., 1979) to describe nuclear volume changes by explicitly coupling the bulk mechanics to an elastic nuclear envelope. In our model, the dynamic is governed by two generic dissipation mechanisms: a surface friction (ξ*_s_*) a bulk friction (ξ*_f_*). The mechanical response is determined by three elastic moduli: the isotropic and shear elastic moduli of the bulk, and the stretching elastic modulus of the envelope (Fig. 5B). We first focused our analysis on the volume loss regime, corresponding to confinement steps from 6 to 5 μm, 5 to 4 μm, and 4 to 3 μm (in HeLa cells, the nuclear envelope rupture regime becomes dominant below 3 μm) (Supplementary Fig.1F; Raab et al., 2016). In this regime of nuclear volume loss, the nuclear envelope is fully unfolded, and the nucleus adopts a smooth and flattened, pancake-like geometry (Fig. 1B; Supplementary information, Section III.I). Exploiting this simplified geometry, we can derive an analytical solution to the model. Moreover, the linear material assumption in our model does not limit its applicability, due to the design of the confinement step experiments. Indeed, while the absolute strains experienced by nuclei range from 50% to 80% relative to their initial sizes (∼12 μm for HeLa cells), each incremental compression step is small—only ∼10–20% relative to the preceding plateau height. This design allows the material response to be treated as linear for each confinement step while still capturing nonlinear behavior by allowing the model parameters to vary between plateaus. This approach enables us to assess the nonlinearity of the nuclear response to confinement without imposing a priori assumptions. Our analysis reveals two key dimensionless parameters: 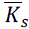, the scaled ratio of surface to bulk moduli, and 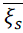, the scaled ratio of surface to bulk friction. Together, these parameters quantify the relative contributions of the bulk and the envelope to both the dynamics and the amplitude of the force relaxation response (Fig. 5B). In the limit where 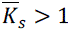 and 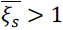, the envelope dominates both the mechanical response and the relaxation dynamics. Conversely, when 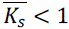 and 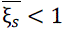, the bulk governs the nucleus behavior. Based on this framework, our model makes several testable predictions, which we experimentally validate.

A key prediction of the model is that nuclear volume dynamics follow a diffusion-like equation, with an effective diffusion coefficient D, as commonly found in poroelastic systems (Doi et al., 2009). Qualitatively, this implies that water escapes from the nuclear periphery, leading to a relaxation of displacement that propagates inward from the edges (Fig. 5C). The slowest characteristic relaxation timescale is set by the square of a relevant length scale (nuclei radius *b_i_*) divided by the diffusion coefficient, 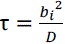. The fact that volume loss follows a diffusion equation provides a clear interpretation of the observed relaxation dynamics (Fig. 1I). If compression occurs much faster than the diffusion timescale, the nucleus deforms at nearly constant volume because water cannot escape rapidly enough. This leads to an initial increase in surface area, tension, and force. Subsequently, as water gradually escapes, the volume shrinks, reducing surface area, tension, and force. Conversely, if the compression timescale is much longer than the diffusion timescale, the process is quasistatic, volume is lost during the confinement, and no significant relaxation is observed post-confinement (Fig. 5D-F & Supplementary Fig. 5A-E).

The model further predicts a multi-timescale relaxation of nuclear volume, with timescales τ*_n_* proportional to the diffusion timescale 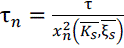. The scaling constants *x_n_* are dimensionless eigenvalues that depend non-trivially on both the relative stiffness 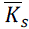 and the relative friction 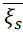 (Supplementary information, Section II.E). Importantly, higher-order modes (larger n) decay faster, such that the relaxation initially involves multiple timescales but eventually becomes dominated by the slowest mode τ_1_, leading to a single-exponential decay at times longer than a few tens of seconds (Supplementary Fig. 5B). Our model thus reproduces the transient multimodal dynamics followed by the single-mode relaxation regime, consistent with our experimental observations (Fig. 1I). Moreover, it recovers limiting regimes previously described in the literature, including classical poroelastic gel models and a surface-friction–limited regime for volume changes (Venkova et al., 2022) (Supplementary information, Section II. F,H).

## Experimental validation of the dynamic model: Force and volume relaxation kinetics

A second prediction of the model is that nuclear volume and force should exhibit the same characteristic relaxation timescales. To test this, we acquired nuclear images at high temporal resolution (5 s intervals) throughout the entire AFM confinement experiment, rather than imaging only at the end of the relaxation phase. This approach enabled us to simultaneously monitor nuclear volume and force dynamics under confinement. Consistent with our model’s prediction, we found that both nuclear force and volume relax with matching temporal profiles (Fig. 6A).

**Figure 6:**
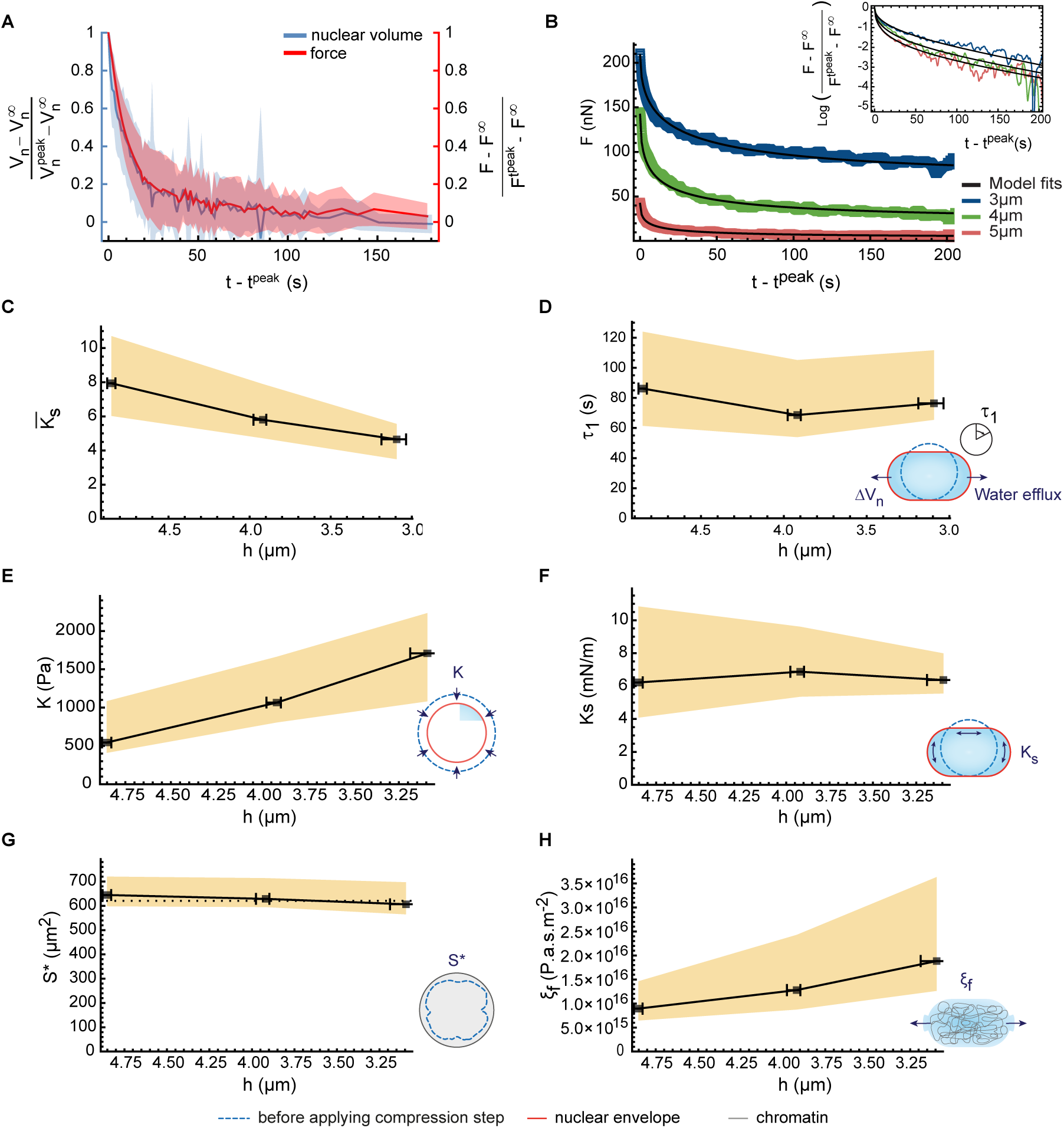
Inference from AFM relaxation curves reveals surface-dominated nuclear mechanics and bulk-controlled volume-loss timescales. **A.** Comparison of relative nuclear volume (blue) and force (red) relaxation dynamics during a 5–4 µm uniaxial confinement step for Latrunculin A treated Hela cells (HeLa-LatA) (*N=3* independent experiments, *n=26* cells). Solid lines and shaded regions represent the mean and standard deviation, respectively. **B.** Representative force relaxation dynamics for a HeLa-LatA cell following successive confinement steps at ℎ = 5μ*m* (red), 4μ*m* (green), 3μ*m* (blue). Black lines indicate the dynamical model fits to the data. Inset: The same dataset and model fits displayed on a log-linear scale. **C-H.** Inferred model parameters across successive confinement steps for HeLa-LatA cells (*N=3* independent experiments*, n=22* cells). Black squares denote the population median, while the whiskers and shaded region represent the 95% confidence intervals for the *x-* and *y-* errors, respectively. Error margins are computed by bootstrapping the median 10^D^times (Supplementary Information Section III.L). Schematic icons in the lower-right corner of the plots illustrate the respective model parameter (defined in Fig. 5B). Dimensional parameters were determined by combining experimental measurements of the midplane nuclear surface area after relaxation with the fitted values of the dimensionless parameters (Supplementary Information Section II.G).

To further quantitatively test our model’s prediction, we developed a pipeline to infer the model parameters from our dynamic measurements (Fig. 6 & Supplementary Fig. 6). Our analysis reveals that HeLa cell nuclei operate in a limiting regime characterized by 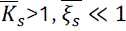 (Fig. 6C; Supplementary information, Section V.E) and that our experiments are done with compression times much shorter than the slowest relaxation timescale τ_1_ (Fig. 6D). In this experimentally relevant regime, the model simplifies and reduces to three fitting parameters (Supplementary information, Section II.G) that we can determine using (i) the peak force immediately after compression, (ii) the relaxed force at steady state, and (iii) τ_1_ by performing a least-squares fit of the force relaxation curve (Fig. 6B).

Our finding that 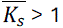 indicates that the nuclear envelope dominates the force response in the volume loss regime of large deformations. This result independently validates the core assumption of our previous quasistatic model and aligns with previous findings in the literature (*e.g.*, Stephens et al., 2017; Finan et al., 2009). While the envelope remains the primary mechanical contributor in the confinement range explored, we observe a progressive decrease in 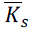, suggesting that the bulk contribution becomes increasingly significant as the nucleus loses volume (Fig. 6C). This trend is explained by the distinct behaviors of the two mechanical constituents: the envelope maintains a constant stretching modulus (Fig. 6F)—in the order of 6 mN/m—consistent with a linear elastic behavior, whereas the bulk exhibits a nonlinear response, with an effective modulus K that increases with compression and ranges on average from 500Pa at 5 µm confinement to 1.5 kPa at 3 µm confinement (Fig. 6E). Notably, the estimated values of these parameters are consistent in order of magnitude with those reported independently in the literature using other methods (*e.g.,* Biswas et al., 2025; Stephens et al., 2017; Dahl et al., 2004). Moreover, the threshold surface *S*^∗^, independently inferred from the dynamic model (*∼*700 µm^4^) (Fig. 6G & Supplementary information, Section I.F) agrees in magnitude (within less than 10%) with the threshold surface obtained from the quasistatic model fit (Fig. 2C and Supplementary information, Section II.G), further supporting the robustness of our characterization. Notably, *S*^∗^ is independent of the confinement step, suggesting that the nuclear envelope does not reorganize within the experimental timescale. This observation is again pointing to a linear-elastic response of the nuclear envelope, even in the high-compression regime, and supports our conclusion that the system operates in the “stiff-envelope” regime, where quasi-static volume loss occurs at approximately constant total surface area thus limiting the extension of the surface to a regime where it remains linear (at least within the tens-of-minutes timescale of our observations) (Fig. 2C). To further test this interpretation, we used the independently inferred mechanical parameters *K* and *K_s_* from the AFM pipeline, without invoking assumptions about their microscopic origins, and verified the criterion for the “stiff-envelope” regime of the previous quasistatic model, namely that the trapped osmolyte osmotic pressure is less than the mechanical pressure due to the envelope Π*_t_* < *P_m_* (Supplementary information, Section I.D.4.d). Together, this dynamic model, and the full mechanical characterization of the nucleus that it provides, reinforces our initial conclusion that, under strong confinement, nuclear volume loss is driven by nuclear-envelope tension.

In addition to the measures of *K*, *K*_&_ and *S*^∗^, our dynamic model gives access to an interpretation of the origin of the relaxation kinetics. While the envelope dominates the mechanics, the fact that 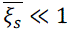 implies that the relaxation is dominated by bulk dissipations. This result is consistent with an order-of-magnitude estimate based on Poiseuille flow through nuclear pore complexes, which suggests that nucleoplasmic flows across the nuclear envelope are not rate-limiting for the observed volume changes (Supplementary information, section II.G.9). In addition, our approach not only allows to measure the relative importance but also absolute values of the frictions. We find that the bulk friction ξ*_f_* ranges between 10^36^Pa.s.m^-2^ at 5 μm confinement to 1.7 ⋅ 10^36^Pa.s.m^-2^ at 3 μm (Fig. 6 H). This value is coherent with the diffusive nuclear dynamics of swelling upon protease digestion reported in (Mazumder et al., 2008), and is 100 to 1000 times larger than the friction in the cytoplasm (Moeendarbary et al., 2013).

## Lamin A/C depletion and chromatin decompaction changes nuclear mechanics and bulk properties

Studies reporting measures and models for nuclear mechanics have been debating the relative importance of surface and bulk components of the nucleus, focusing on the relative roles of the nuclear lamina and of the chromatin (Stephens et al., 2017). We thus perturbed these two main nuclear structures using Lamin A/C knockdown (Lamin A/C is the main lamina component in HeLa cells) and trichostatin A (TSA) to reduce heterochromatin, which globally de-compacts the chromatin. We then applied our dynamic model to the sequential step confinement data to extract the main mechanical parameters of the perturbed nuclei (Fig. 7A,B and Supplementary Fig. 7.A.A-D provide an integrative comparison of all the parameters for the 6 to 5 μm, 5 to 4 μm and 4 to 3 μm confinement steps; Fig. 7 C-F; Supplementary Fig. 7A, E-H; Supplementary Fig. 7B,C,D show each parameter across all confinement heights for the different perturbations). As expected, we observed a significant reduction in the stretching elastic modulus for the Lamin A/C depletion — ∼50% on average (of note, Western blot analysis indicates a partial depletion; Supplementary Fig. 3B). Importantly, despite the decreased envelope modulus, the extent of nuclear volume loss was similar to control cells, reflecting a decreased bulk modulus (Supplementary Fig. 7A, F,G). This suggests a complex response to Lamin A/C depletion where both the envelope stretching modulus and the bulk modulus K decreases (Fig. 7B), effectively compensating each other’s effect on volume loss.

**Figure 7:**
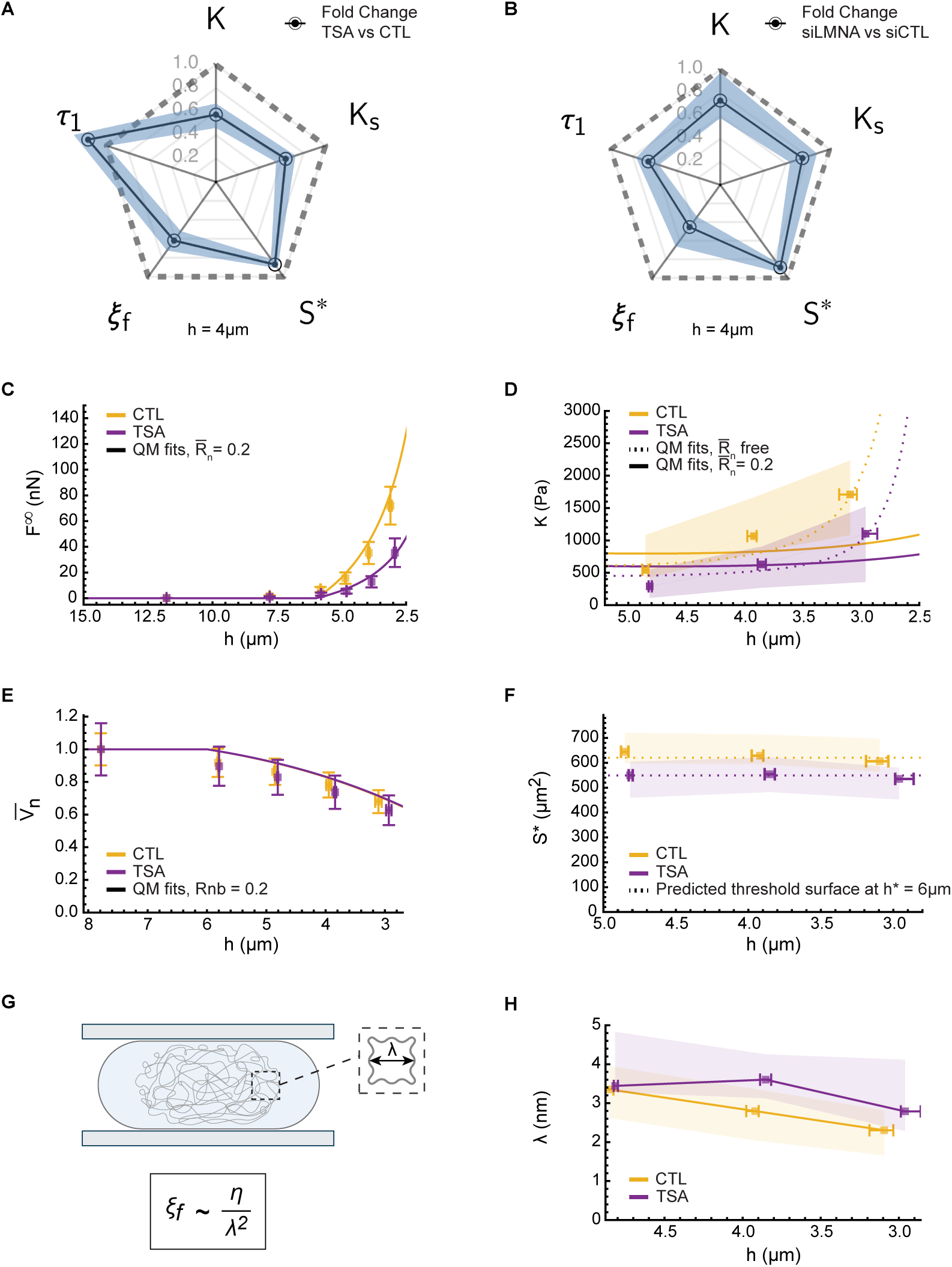
Biological perturbations reveal tight coupling between bulk and nuclear envelope mechanics. Unless otherwise indicated, all cells were treated with Latrunculin A (2μ*M* incubated 20 minutes before experiments). **A,B.** Fold-changes in inferred model parameters (defined in Fig. 5B) for a 5-4 µm confinement step under distinct biological perturbations. **A**. Trichostatin A (TSA; 100*nM* concentration for 16h) treatment (*N=3* independent experiments*, n=22* cells) relative to control (CTL) (*N=3, n=22*) in HeLa cells. **B**. Lamin A/C depletion via siRNA (siLMNA) (*N=10, n=45*) relative to the non-targeting control (siCTL) (*N=7, n=32*). Circles denote the population median, and error bars represent the 50% confidence interval (CI) obtained by bootstrapping the median 10^D^times (Supplementary Information Section III.L). **C-F.** Quasistatic model (QM) validation. **C,D.** Quasistatic model fits to the post-relaxation force *F*^9^ and bulk modulus *K*— as inferred from the dynamic model (Fig. 6 E)— for control and TSA-treated HeLa nuclei (Supplementary Information Section I.H), where 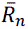 denotes the dry volume fraction of the undeformed nucleus inaccessible to diéusing osmolytes. **E.** Predicted normalized nuclear volume across successive confinement steps. Volumes are normalized to the baseline volume measured at the confinement height ℎ = 8μ*m*, a regime characterized by negligible volume loss where the simpler nuclear geometry minimizes measurement and estimation noise. **F.** Comparison of the threshold surface inferred from the dynamic model at the successive confinement steps with the quasistatic model prediction at a threshold height of ℎ^∗^ = 6μ*m*. For raw data (e.g., force and normalized volume), squares and whiskers denote the mean ± 95% CI. For inferred parameters (bulk modulus and threshold surface), squares denote the median and error bars indicate the 95% CI via bootstrap resampling (Supplementary Information Section III.L). **G, H,** Estimated microscopic nuclear bulk pore size. Pore size estimates are calculated under the assumption that bulk hydraulic friction is dominated by viscous dissipation arising from nucleoplasmic flow through the porous chromatin network. Calculations utilize a baseline nucleoplasmic viscosity of η ≈ 0.1 *Pa*. *s* alongside the empirical bulk friction coeéicients inferred for control and TSA-treated HeLa nuclei (Fig. 6H & Supplementary Fig.7B F). Squares represent the median and error bars denote the 95% confidence interval obtained via bootstrapping resampling (see Supplementary Information, Section III.L).

This raises the question of the microscopic origin of the nuclear bulk modulus *K*. We first asked whether the mechanism proposed in our quasistatic model—namely that *K* is primarily an osmotic modulus set by trapped nuclear osmolytes—can account for both the measured magnitude of *K* and its nonlinear increase. We fitted the quasistatic model as in Fig. 2C, here using our independent AFM-based mechanical characterization of HeLa cell nuclei to further constrain the fit (Fig. 7C-F; Supplementary Fig. 7.A, E-H). When the pre-confinement nuclear dry volume fraction (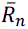) is treated as a free parameter, the quasistatic model captures the observed nonlinear behavior of the bulk modulus at high confinement (∼ 3 μm), though this agreement is strictly limited to regimes where 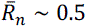. This value is larger than literature estimates for the whole cell (0.1–0.3), although no direct measurements specific to the nucleus are available to our knowledge. We therefore fixed 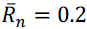 and fitted only the two remaining parameters: γ*_c_*, the plasma-membrane tension, and 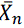, which reflects the ratio of the osmotic pressure due to trapped nuclear osmolytes to that of the external medium.

Under this constrained strategy, the model recovers the magnitude of *K* close to the confinement threshold (∼ 5 μm) for realistic concentrations of trapped nuclear osmolytes (∼ 0.1 mM, corresponding to an osmotic pressure of ∼ 200 Pa in control cells), but fails to reproduce the nonlinearity observed at the highest confinement (∼ 3 μm). We interpret this discrepancy as evidence for additional nonlinear effects not included in the quasistatic microscopic model. These may include higher-order osmotic contributions, such as steric repulsion, that become relevant at very high confinement. In addition, the decrease in 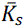 under strong confinement suggests that chromatin may also contribute a nonlinear component to the nuclear osmotic pressure.

Our analysis also provides a measure for a bulk friction value, inferred from our pipeline. In analogy with hydrogels, it could be interpreted as originating from a microscopic dissipation between the chromatin network and the nucleoplasm. Interestingly, its value is decreased (∼60%) by treatment with TSA, which de-compacts chromatin (Fig. 7A). We thus wanted to understand better what we could learn from this number. Based on a dimensional analysis, and considering a simplified picture of the nuclear bulk viewed as a chromatin gel permeated by a viscous nucleoplasm, we expect the bulk friction to scale as the nucleoplasmic viscosity divided by the square of the typical pore size of the chromatin meshwork (Fig. 7G,H). Using a nucleoplasmic viscosity of 10^83^ Pa.s (Mazumder et al., 2008), we estimate a typical pore size on the order of few nanometers. While this estimate is somewhat small, the typical size of a histone being around (∼10 nm), it remains reasonable (Cutter et al., 2015; Vovard et al., 2026). Since TSA treatment strongly decreased the bulk friction, heterochromatin could represent a major source of friction, explaining the low number we found for the pore size. Notably, this perturbation also altered other mechanical parameters, including the surface stretching modulus, suggesting that the chromatin state could also impact the mechanics of the nuclear surface. Reciprocally, the depletion of Lamin A/C also reduced the bulk friction coefficient, suggesting an effect of Lamin A/C depletion on chromatin. Overall, our results show that nuclear surface and bulk mechanics are tightly coupled: perturbations typically assigned to one compartment consistently affect the other, indicating that these contributions cannot be treated independently.

## Discussion

### A unified framework for nuclear mechanics under deformation

Our integrative approach—combining controlled confinement assays, high-resolution imaging, atomic force microscopy, and theoretical modeling—has revealed a bimodal mechanical response of the nucleus to large deformations. Above a threshold confinement —typically 5–6 µm, depending on the cell line—nuclei deform at nearly constant volume, behaving as incompressible objects. Below this threshold, nuclei lose up to 30% of their volume, while maintaining an almost constant surface area. This transition is governed by the onset of nuclear envelope (NE) tension and the unfolding of excess surface area stored in membrane folds. By quantitatively modeling these regimes, we demonstrate that the nucleus operates as a poroelastic material: the envelope dominates the mechanical resistance, while dissipation is controlled by the chromatin network, on the minute timescale. These findings reconcile seemingly contradictory observations in the literature and provide a physical basis for understanding how the nucleus responds to external forces.

### Interpreting force compression curves: A mechanistic understanding of nuclear deformation

One of the key advances of this work is the ability to fully interpret the force required to deform a cell nucleus on the seconds-to-minutes timescale. In particular we show that the nucleus behaves as a strain-rate stiffening object. When deformation occurs faster than volume loss, it proceeds at constant volume, and the force reaches a peak corresponding to the extension of the nuclear envelope. As the nucleus begins to lose volume, the force decreases, and the envelope tension relaxes, reaching a plateau determined by the equilibrium between the pressure exerted by concentrated trapped osmolytes (the bulk osmotic modulus) and the pressure due to NE tension. The force plateau increases with stronger confinement, as the envelope extends and the nucleoplasm concentrates due to volume loss.

### Implications for cell migration

This mechanistic framework—specifically the interplay between deformation speed and the timescale of water efflux— is critical for understanding how cells navigate confined environments, such as during migration through narrow gaps. When deformation is faster than the timescale of water efflux, the nucleus behaves as incompressible, causing strong NE stretching and high resistance to passage through narrow constrictions. Conversely, if water is expelled on similar timescales to deformation, nuclei can adapt by reducing volume at a lower force cost (Stöberl et al., 2024). Nuclear volume loss would thus not be a limitation for slow moving cells, but it could be a crucial aspect of fast immune cell migration, which occurs on the minute timescale. Fast-migrating cells, such as leukocytes, may exploit this by maintaining highly deformable nuclei with low Lamin A/C content and modulating chromatin compaction to increase nuclear permeability (Zhao et al., 2024, Blazquez-Romero et al., 2026). Dynamic regulation of nuclear volume may thus represent a key strategy, complementing nuclear softening, fold opening, or blebbing, to enable rapid migration through dense tissues (for a recent review on the impact of nuclear properties on cell migration see Renkawitz et al. 2026).

### The role of nuclear envelope folds: A buffer for mechanical adaptation

The existence of a “safe deformation” range (between ∼12 µm and 6 µm) is a striking feature of nuclear mechanics. In this regime, the nucleus deforms without significant changes in tension or volume, thanks to the mobilization of excess surface area stored in folds that have been proposed to originate from nuclear assembly at mitotic exit (McKee et al., 2025; Chu et al., 2017). The progressive unfolding of these folds buffers tension until they are exhausted, at which point the nucleus sharply transitions to a mechanosensitive regime with increasing NE tension and volume loss. This duality explains why some studies report nuclear incompressibility (*e.g.,* during cell spreading or substrate stiffening (McKee et al., 2025;), while others observe dramatic volume changes (e.g., during migration through micron-sized pores or micropipette aspiration, Rowat et al., 2006). The threshold behavior is conserved across cell types, despite variations in nuclear size, suggesting a general mechanism for mechanical adaptation.

### Nuclear volume loss and its physiological consequences

The onset of volume loss has profound implications for nuclear physiology. Since volume decreases on the order of minutes—much faster than nuclear protein turnover or import/export—the nuclear dry mass remains constant, while macromolecular crowding increases. We observed a sharp rise in the number of nuclear condensates beyond the volume loss threshold (Fig. 4B), consistent with recent findings on cells during confined migration (Zhao et al., 2024). Given the sensitivity of condensates to crowding and their role in transcriptional regulation, volume loss may directly couple mechanics to gene expression. Concurrently, NE tension is known to promote the import of mechanosensitive transcription factors such as YAP/TAZ (Andreu et al., 2022), suggesting that both crowding and tension-sensitive transport could contribute to nuclear mechano-sensing in the volume loss regime. At higher deformations, envelope tension may exceed the mechanical strength of the lamina and intranuclear pressure surpass the binding strength of the inner nuclear membrane to the lamina, leading to nuclear blebbing and nuclear envelope ruptures (Raab et al. 2016; Denais et al. 2016). Notably, confocal imaging reveals that chromatin is often absent or only partially present in blebs (Supplementary Fig. 1E), supporting the idea that nuclear blebbing is primarily driven by nucleoplasmic pressure, caused by trapped soluble nuclear osmolites, such as proteins (Deviri et al., 2022; Lemière et al., 2022, Rollin et al., 2023) rather than by the chromatin polymer itself.

### Dynamic relaxation: Diffusion-like behavior and the poroelastic nucleus

Our dynamic model reveals that nuclear volume relaxation follows diffusion-like dynamics, with a characteristic time scaling as the square of the projected nuclear radius (*b*^i^, see Fig. 5A) divided by an effective diffusion coefficient. This explains why volume relaxation occurs over tens of seconds, consistent with a nuclear radius of ∼6 µm and a nanometer-scale chromatin mesh size. Although the scaling relationship between the nuclear volume loss timescale, nucleus radius, and diffusion coefficient resembles that of a simple hydrogel, the presence of an additional elastic envelope alters the constant of proportionality (Fig. 5B,C). Importantly, this distinction results in the timescale of nuclear volume loss remaining constant and independent of confinement height, unlike in a hydrogel (Fig. 6D). The inferred bulk friction values correspond to pore sizes of a few nanometers, close to estimates for dense heterochromatin, but still smaller than expected from the literature. This might be because the value we considered for nucleoplasm viscosity, that we did not measure directly, is underestimated, or because more complex processes have to be considered due to the architecture of the chromatin at the nanometer scale, and the large variety of sizes of solutes present in the nucleoplasm. The slow relaxation of nuclear volume, compared to cytoplasmic volume regulation, highlights why the nuclear-to-cytoplasmic (NC) ratio can transiently deviate even when average scaling is preserved. Thus, both the amplitude and the rate of deformation determine whether the nucleus resists incompressibly or relaxes by losing volume.

### Comparing nuclear and cytoplasmic volume regulation

While the cytoplasm and nucleus are governed by the same physical laws of osmosis and force balance (Stewart et al., 2011), they dispaly fundamentally different volume regulation responses. The cytoplasm has an osmotic bulk modulus in the range of hundreds of kPa, due to ∼100 mM of trapped osmolytes. Deformation does not directly reduce cytoplasmic volume but instead acts indirectly via changes in cortical tension and ion fluxes across the plasma membrane (Venkova et al., 2022). In contrast, the nucleus has a much lower osmotic modulus (∼500 Pa), making it directly sensitive to NE tension. When the envelope becomes taut, Laplace pressure is sufficient to drive efflux of water plus small osmolytes, resulting in the concentration of nuclear proteins. This difference explains why cell volume is buffered against direct mechanical compression, while nuclear volume is readily affected.

### Microscopic origins of nuclear mechanics: chromatin, lamins, and osmolytes

Our analysis provides insight into the microscopic origins of nuclear mechanics. The osmotic modulus we measure corresponds to a trapped osmolyte concentration of ∼0.1mM—roughly 10 times lower than the total nuclear protein concentration. Importantly, determination of the bulk modulus does not depend on the model, but only on measurements of the volume and’e’ the force, which makes this measure robust. A low modulus could come from an overestimation of the volume loss, but given the very flat geometry and unfolded nucleus, our measure of the volume change is precise and our observation of an increase in the number of condensates is consistent with a significant volume loss.

This suggests that our measure of *K* can be trusted and that only a small fraction of nuclear macromolecules contribute to the nuclear osmotic pressure. Several studies point to proteins as major nuclear osmolytes (Deviri et al., 2022; Lemière et al., 2022). Since their total concentration is in the mM range, it suggests that only ∼10% contribute to the osmotic pressure —those that are unbound, mobile within the nucleus, but too large to freely pass through NPCs. It also suggests that other species could contribute to a similar extent to the osmotic pressure in the nucleus (counter-ions, entropic contribution of chromatin), and that precise quantification would be required to separate these contributions (for example, there are very few measures of the nuclear membrane potential; Loewenstein et al., 1963). Moreover, this value of *K*, although low relative to the osmotic modulus of the whole cell, corresponds to the typical forces exerted by the cell cytoskeleton, explaining how cells could exert enough forces to deform their nucleus and induce nuclear membrane blebbing during migration (Raab et al., 2016, Denais et al. 2016).

The low osmotic modulus also helps explain why changes in nuclear protein content or nucleocytoplasmic transport strongly affect nuclear volume. Importantly, measuring the bulk modulus in a variety of cell types and perturbations reveals that this parameter can adopt a large range of values, which might reflect that cells can easily play on their nuclear content to modulate how physical constraints affect the nuclear volume. Interestingly, Lamin A/C depletion reduces not only NE stiffness but also the osmotic bulk modulus, suggesting that lamins influence both envelope mechanics and osmotic pressure, potentially through changes in import/export rates, or non-polymerized lamin monomers that are known to be abundant in the nucleus. This dual contribution may allow cells to restore nuclear volume homeostasis after deformation by simultaneously tuning NE stiffness and internal osmolyte concentration via modulation of import/export rates.

Our results highlight the nonlinear response of the nucleus to deformation. While the surface remains in a linear elastic regime (as volume loss occurs at nearly constant NE surface area), the bulk exhibits nonlinear behavior, with an effective modulus that increases with compression, potentially due to the concentration of trapped osmolytes.

### Nuclear volume as a central regulator of mechanotransduction

This study provides an integrated view of nuclear volume regulation under mechanical confinement, resolving long-standing contradictions in the literature by identifying a tension threshold that separates constant-volume and volume-loss regimes. We establish that nuclear mechanics is governed by the envelope, while dissipation is controlled by chromatin, consistent with recent work showing that BRG1-dependent chromatin remodeling can actively fluidize metabolically intact nuclei (Byfield et al., 2025). By quantifying the scales involved—20% volume loss, ∼1 kPa osmotic modulus, ∼50-second relaxation timescales, and nanometer-scale pore sizes—we demonstrate that nuclear volume is a critical determinant of cell migration and mechanosensing. Our results also complement recent in vivo work in developing Drosophila flight muscle, where mechanical pressure on nuclei was shown to promote Tono condensate formation and trigger a transcriptional transition during myogenesis (Zhang et al., 2024). Together, these observations support a view in which nuclear deformation, compression and volume regulation contribute to mechanotransductive control of cell state. These findings open new avenues for exploring how cells modulate nuclear mechanics and water permeability to adapt to mechanical challenges in physiological and pathological contexts, such as immune surveillance, cancer metastasis and tissue development.

### Toward a predictive model of nuclear adaptation

Future work could extend this framework to other cell types, particularly those with distinct nuclear architectures (*e.g.,* stem cells, neurons, or immune cells). Indeed, our approach, while applied here mostly on one cell type, is general and could constitute a reference method to compare nuclear properties, taking advantage of the full characterization of both surface and bulk properties of the nucleus that it offers (see Nava et al. for an application to multiple cell types and perturbations). Additionally, probing the dynamic regulation of nuclear permeability in specific biological contexts such as the maturation of immune cells (*e.g.,* dendritic cells, or neutrophils), may reveal how cells actively tune their nuclear mechanics in response to environmental cues to achieve their function more efficiently. Finally, integrating these mechanical insights with transcriptional and signaling data could elucidate how fine tuning of nuclear properties, by modulating nuclear volume loss and envelope tension impacts gene expression (McCreery et al., 2025).

### Limitations and open questions

While our model provides a robust framework for understanding nuclear mechanics, several questions remain. First, the microscopic origin of the bulk friction—whether it is dominated by specific chromatin structures or by molecules that modulate nucleoplasmic viscosity, or even other sources—requires further investigation (Mazumder et al., 2008; Zhang et al., 2017; Zhang et al., 2020; Vovard et al., 2026). Second, the role of actin and other cytoskeletal elements in nuclear mechanics is complex. Depolymerizing actin did not affect the extent or kinetics of volume loss, suggesting that actin does not directly regulate nuclear osmolarity or volume changes. However, actin may influence NE tension or surface mechanics, as the nucleus is mechanically coupled to the cytoskeleton via the LINC complex (Lombardi et al., 2011). Third, the nonlinearity observed in the bulk modulus at high deformations could arise from multiple sources, including chromatin compaction, changes in osmolyte mobility, or additional steric repulsions between trapped osmolytes due to increase concentration or reorganization of the lamina. Our results suggest that the dominant nonlinearity originates from the bulk rather than the surface, but this may vary across cell types or conditions. Finally, our work focuses on the seconds to minutes timescale, but other mechanisms could become important on longer times. For example, given the importance of the bulk modulus and its dependance on the amount of trapped nuclear proteins (meaning actively imported proteins), the well documented impact of nuclear deformation on nuclear import could lead to a change of the bulk modulus on the hour timescale, generating further non-linear behaviors (also discussed in Pennacchio et al., 2024). This could lead to a long-term increase in intranuclear pressure, explaining the observation of persistent nuclear blebbing after hours of confinement (Nader et al., 2021). Similarly, confinement has been shown to lead to changes in the composition of the nuclear lamina, both at the post-translational and transcriptional levels (Discher et al., 2005; Alraies et al., 2024;), further underlying the need, but also the complexity, of addressing nuclear mechanics on the hour timescale.

## Supporting information

Supplemental model and methods

## Acknowledgements

We thank members of the Sens and Piel teams for helpful discussions. We thank Quan Yuan for help with the condensate experiments and Anne-Sophie Macé for help with image analysis. This work was supported by an ERC Synergy Grant (101071470 SHAPINCELLFATE) and an ANR grant (ANR-21-CE15-0009 MECFUN) to M.P., and by Swiss National Science Foundation grants 51NF40-205608 and 31003A-182587 to D.J.M

