## Supplemental Figures for "Coupled nucleoplasmic flows and envelope expansion govern the mechanical resistance of the nucleus"

### Supplementary Figures:

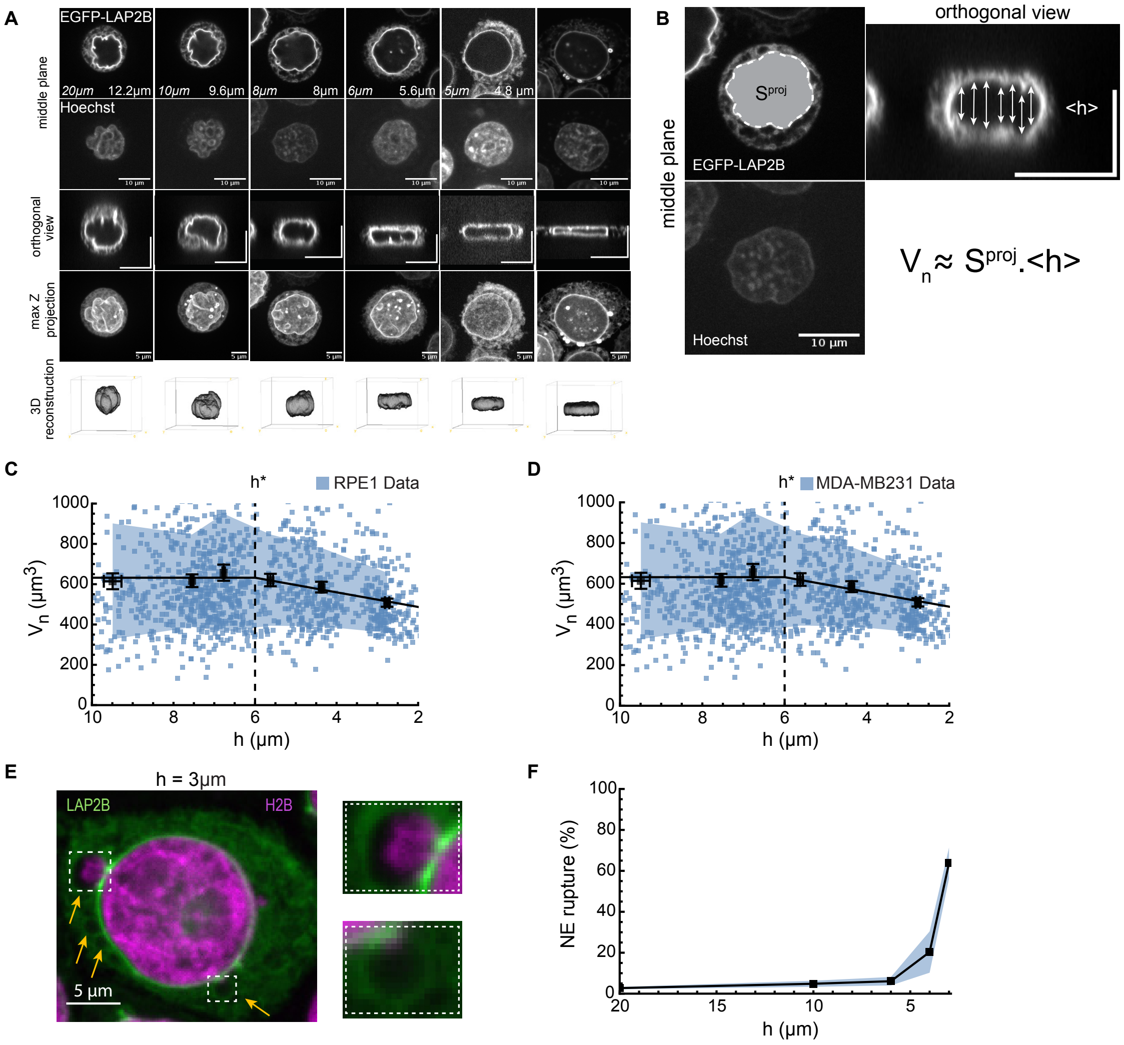

2026_Rollin et al. Sup. Fig1

**Supplementary Figure 1:**

**A.** Representative EGFP-LAP2β-stained HeLa nuclei under varied confinement heights (top rows) and corresponding 3D reconstructions obtained via Hoechst staining (bottom row). Italicized labels indicate the nominal confinement heights (*h*), while non-italicized labels represent the measured maximum nuclear heights from orthogonal views (top row).

**B.** Example of nuclear volume calculation for static six-well confinement at $h = 8\mu m$ (Supplementary Information Section III.I). Utilizing the image profiles from panel **A**, the nuclear volume is estimated by multiplying the projected midplane area ($S^{proj}$) by the average nuclear height ($< h >$) across spatial positions.

**C,D.** Nuclear volume ($V_n$) of RPE1 (**C**) and MDA-MB-231 cells (**D**) as a function of confinement height (*h*). Blue squares account for individual data points. Data are binned into intervals of equal sample size (*N=6,n=1233 for RPE1 cells, N=5,n=550 for* MDA-MB-231 cells), with black squares denoting the binned mean ± 95% confidence interval (CI). A piecewise linear fit (black line) is provided as a visual guide, with the standard deviation indicated in light blue. Although the threshold confinement height for nuclear volume loss ($h^*$) cannot be precisely determined from these data, it falls within the *5-6 μm* range.

**E.** Representative fluorescence micrograph of a HeLa cell nucleus co-expressing H2B (chromatin; pink) and LAP2β (nuclear membrane; green). Yellow arrows denote nuclear membrane blebs; dashed white boxes indicate regions shown at higher magnification in the insets (right). These insets reveal heterogeneous chromatin occupancy within the blebs, ranging from partial filling (top) to complete exclusion (bottom).

**F.** Quantification of fraction of ruptured nuclei across varying confinement heights in Hela cell nuclei with (*N=3, n=315*). Black squares and shaded region denote the mean ± 95% confidence interval (CI).

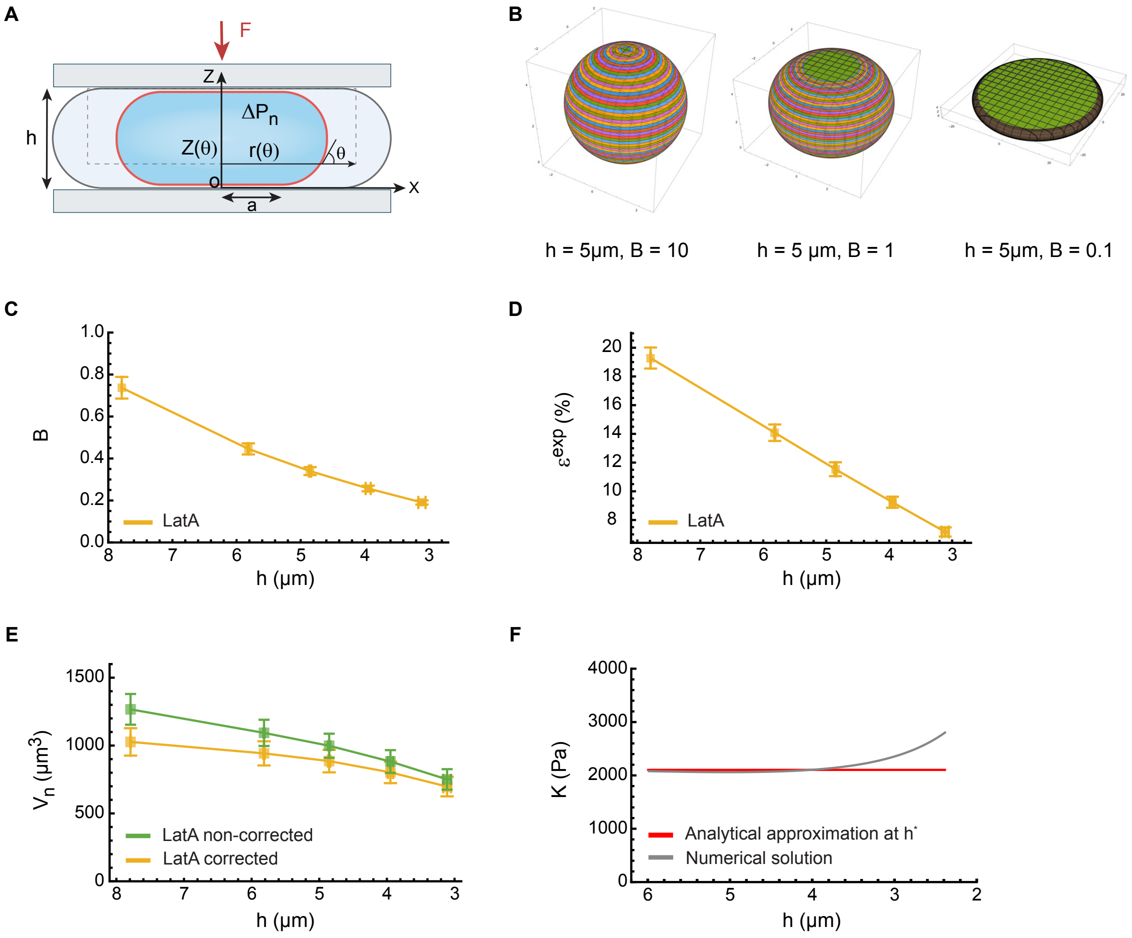

2026_Rollin et al. Sup. Fig2

**Supplementary Figure 2:**

**A.** Schematic parametrization of the quasistatic model. For a nucleus confined at height $h$, the envelope profile is described by the tangent angle $\theta \in [0, \pi]$ relative to the horizontal axis. Spatial nuclear envelope coordinates are mapped by the functions $r(\theta)$ and $z(\theta)$, where $a = r(\theta = 0)$ represents the nucleus-substrate contact radius. The nuclear shape is resolved by formulating a local force balance at the nuclear envelope considering the compressive force $F$, the hydrostatic pressure difference at the nuclear envelope $\Delta P_n$, and the envelope tension (Supplementary Information Section I.C).

**B.** Theoretical nuclear profiles predicted by the model. Range of geometric shapes computed at fixed confinement height $h$ and for various values of the dimensionless parameter $B$. This metric compares the nucleus-substrate contact radius $a$ to the lateral curvature of nuclear envelope (Supplementary Information Section I.C).

**C-E.** Nuclear volume correction in the AFM experiments. **C.** Inferred model parameter $B$ across confinement heights $h$ in Latrunculin-A treated (LatA) HeLa cells, used to determine the corrected nuclear volume from the model (*N=3* independent experiments*, n=22* cells). **D.** Relative error of the raw nuclear volume approximation (height * midplane area) with respect to the model-corrected volume (Supplementary Information Sections III.I and III.J). The error converges to 0 as the confinement is increased. **E.** Comparison between the approximated and corrected nuclear volumes for LatA HeLa cells. In all plots, square points and whiskers represent the mean and 95% confidence interval (CI), respectively.

**F.** Nuclear bulk modulus ($K$) predicted by the quasistatic model using parameter values obtained from the literature (Supplementary Information Section I.E). The red solid line indicates the analytical solution derived in the linear regime near the threshold height ($h^*$); the gray line indicates the numerical solution.

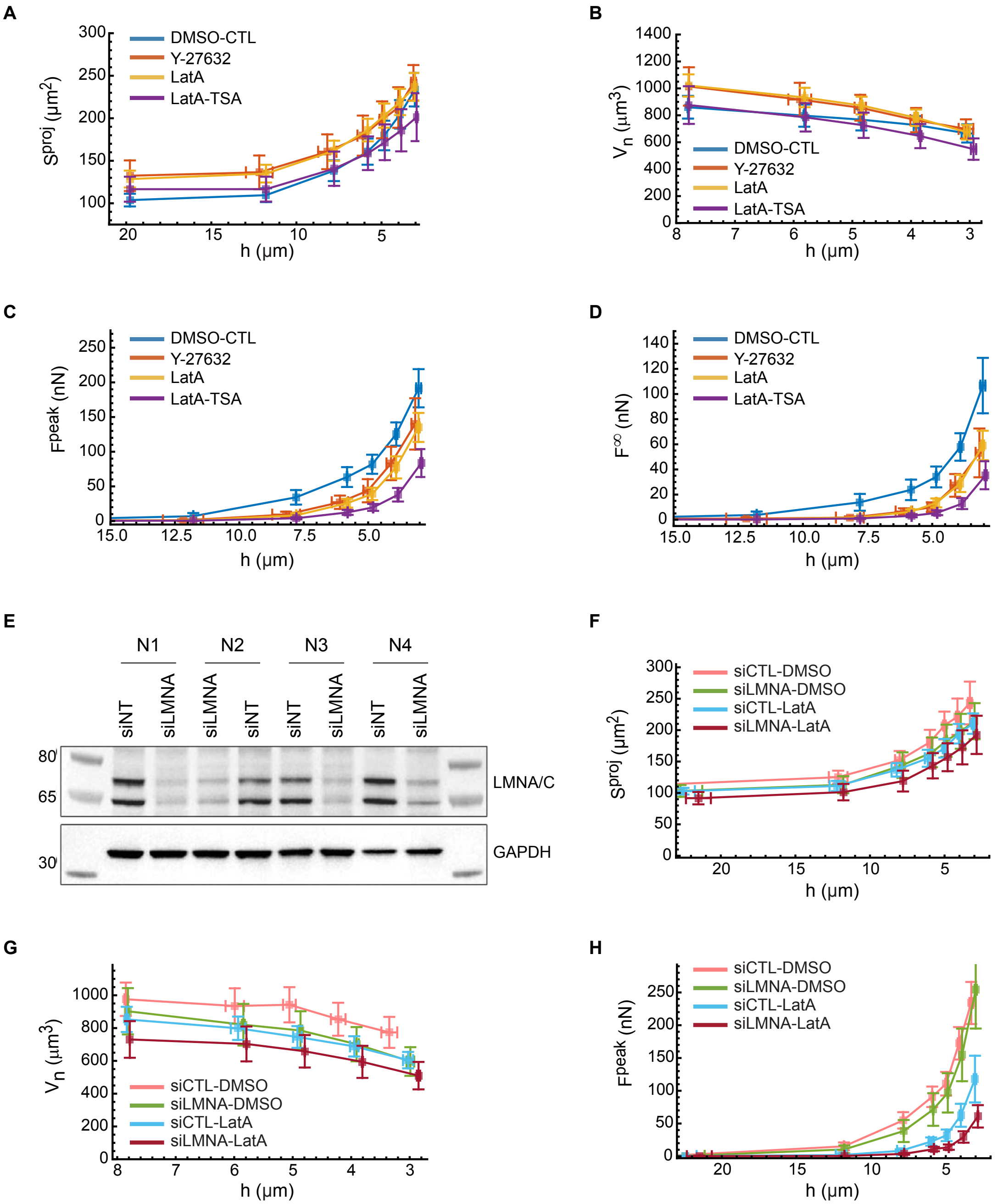

2026_Rollin et al. Sup. Fig3A

**Supplementary Figure 3.A:**

**A-D.** Midplane surface area (**A**), nuclear volume (**B**), peak force (**C**), and post-relaxation force (**D**) measured at varying confinement heights in HeLa cells subjected to four perturbations: DMSO control (N=6 independent experiments, n=25 cells), Y-27632 (N=3,n=21), Latrunculin A (LatA; N=6,n=37), and a combined LatA and Trichostatin A treatment (LatA-TSA; *N=3,n=22*). Squares and whiskers denote the mean ± 95% CI.

**E.** Representative Western blots from four independent biological replicates confirming the efficiency of LMNA/C knockdown (siLMNA) compared with non-targeting siRNA controls (siNT). Additional examples and corresponding quantifications are provided in Supplementary Fig. 3B.

**F-H.** Midplane surface area (**F**), nuclear volume (**G**), peak force (**H**) across varying confinement heights in HeLa cells subjected to four distinct perturbations: control siRNA treated with DMSO (siCTL-DMSO; *N*=3, *n*=26), LaminA/C knockdown treated with DMSO (siLMNA-DMSO; *N*=4, *n*=15), control siRNA with Latrunculin A (siCTL-LatA; *N*=10, *n*=45), and Lamin A/C knockdown with Latrunculin A (siLMNA-LatA; *N*=7, *n*=32). Squares and whiskers denote the mean ± 95% CI.

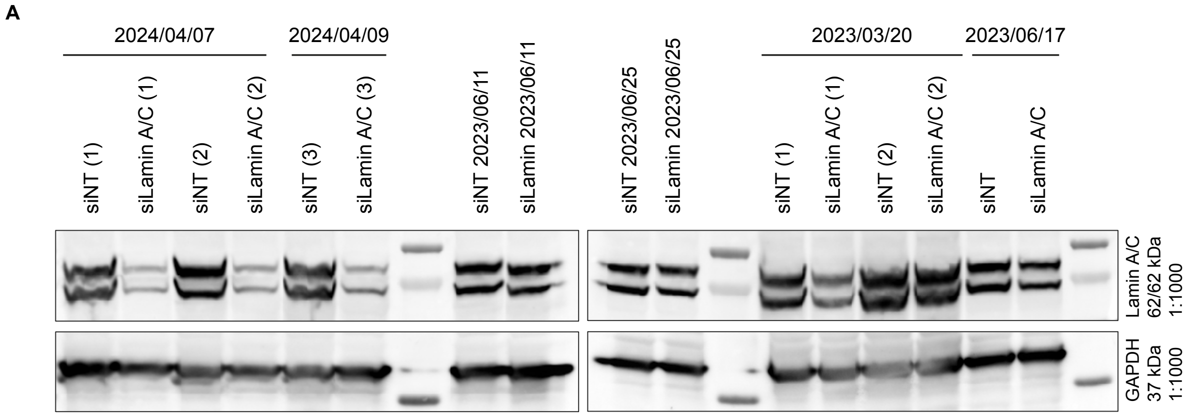

| | area GAPDH | area LaminA/C | area Lamin AC/area GAPDH | ratio |
|---|---|---|---|---|
| siCTR (1) 2024/04/07 | 24221.439 | 25220.317 | 1.041239416 | 1 |
| siLamin A/C (1) 2024/04/07 | 13524.296 | 4296.962 | 0.317721677 | 0.305138 |
| siCTR (2) 2024/04/07 | 17728.782 | 28615.397 | 1.614064463 | 1.550138 |
| siLamin A/C (2) 2024/04/07 | 14957.276 | 5727.669 | 0.382935302 | 0.367769 |
| siCTR (3) 2024/04/09 | 18620.196 | 25919.66 | 1.392018645 | 1.336886 |
| siLamin A/C (3) 2024/04/09 | 21371.539 | 6740.296 | 0.315386552 | 0.302895 |
| siCTR 2023/06/11 | 26298.589 | 21221.095 | 0.806929033 | 0.77497 |
| siLamin A/C 2023/06/11 | 33562.024 | 19415.238 | 0.578488294 | 0.555577 |

| | area GAPDH | area LaminA/C | area Lamin AC/area GAPDH | ratio |
|---|---|---|---|---|
| siCTR 2023/06/25 | 20913.054 | 15026.953 | 0.718544169 | 1 |
| siLamin A/C 2023/06/25 | 19290.255 | 12126.518 | 0.628634406 | 0.874872 |
| siCTR (1) 2023/03/20 | 16992.43 | 21921.004 | 1.290045273 | 1.79536 |
| siLamin A/C (1) 2023/03/20 | 7393.267 | 10660.761 | 1.441955363 | 2.006773 |
| siCTR (2) 2024/03/20 | 7702.589 | 25763.69 | 3.344809128 | 4.654981 |
| siLamin A/C (2) 2024/03/20 | 12885.054 | 24441.439 | 1.896882931 | 2.639897 |
| siCTR 2023/06/17 | 11817.095 | 16731.205 | 1.41584755 | 1.970439 |
| siLamin A/C 2024/06/17 | 16260.359 | 13290.569 | 0.817360121 | 1.137522 |

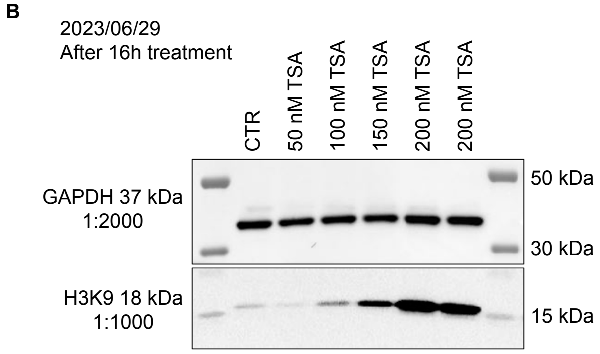

| | Area Plot GAPDH | AreaPlot H3K9 | area H3K9/areaGAPDH | ratio |
|---|---|---|---|---|
| CTR | 19099.669 | 2029.761 | 0.106272051 | 1 |
| 50 nM TSA | 16096.811 | 803.154 | 0.049895225 | 0.469505 |
| 100 nM TSA | 18116.983 | 3941.024 | 0.217532025 | 2.046935 |
| 150 nM TSA | 15827.054 | 6809.643 | 0.43025335 | 4.048603 |
| 200 nM TSA | 20433.589 | 12157.593 | 0.594980794 | 5.598657 |
| 500 nM TSA | 19302.497 | 14473.149 | 0.749807085 | 7.055544 |

2026_Rollin et al. Sup. Fig3B

**Supplementary Figure 3.B:**

**A,B.** Representative Western blots with corresponding quantifications showing the efficiency of LMNA/C knockdown (**A**) and TSA treatments (**B**). TSA was used at 100 nM for 16 h. For LMNA/C knockdown, cells were treated with siLMNA and compared with non-targeting siRNA controls.

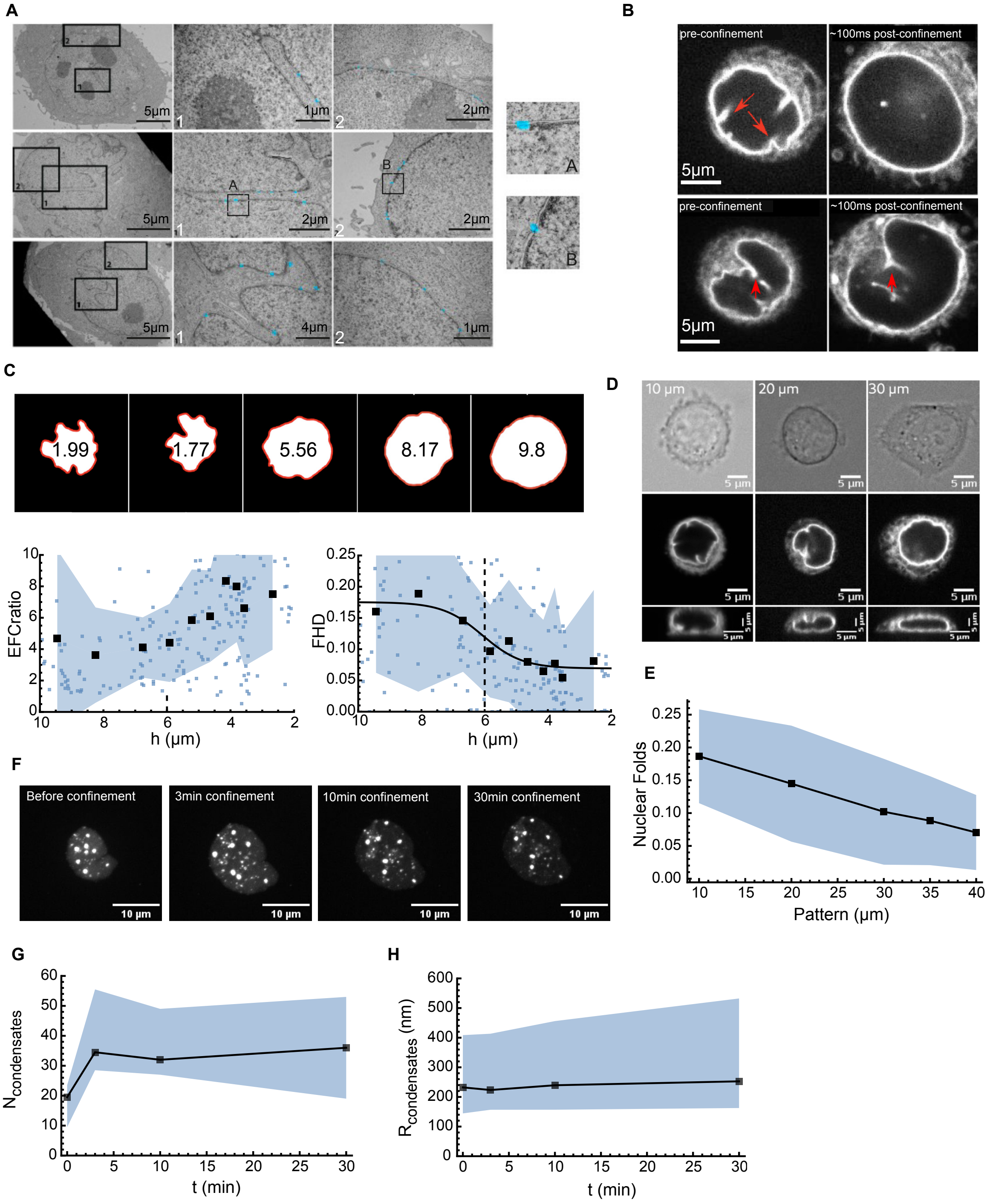

2026_Rollin et al. Sup. Fig4

**Supplementary Figure 4:**

**A.** Representative electron microscopy images of HeLa cell nuclei. Numbered black boxes indicate regions magnified in the corresponding insets, either within a nuclear envelope fold (**1**) or outside a fold (**2**). Lettered black boxes (A, B) indicate regions further magnified to show nuclear pore complexes (NPCs), which are highlighted in blue. No apparent structural differences are observed between folded and non-folded regions of the nuclear envelope.

**B.** Representative images of two HeLa cell nuclei expressing EGFP–LAP2β, shown before and 100 ms after confinement to 8 µm using the dynamic confiner device. Red arrows indicate nuclear folds. Confinement to 8 µm does not significantly reduce nuclear volume (Fig. 1c), but is associated with nuclear envelope smoothening at the single-nucleus level.

**C.** Population-level nuclear envelope smoothening upon confinement. Top, elliptic Fourier coefficient ratio (EFC ratio) for representative HeLa cell nuclei. Bottom, population quantifications of EFC ratio and fraction of high-distance points (FHD) in HeLa cell nuclei (*N = 3* independent experiments, *n = 218* nuclei; see Supplementary Information, Section III.J). Blue squares indicate individual nuclei, the shaded region represent the standard deviation, black squares indicate binned data (20 nuclei per bin), and the solid line shows a sigmoidal least-squares fit used as a visual guide. The dashed line indicates the confinement-height threshold, $h^*$, below which nuclear volume loss occurs, as determined in Fig. 1c.

**D, E.** HeLa nuclear spreading area correlates with nuclear envelope fold opening. **D.** Representative images of HeLa cell nuclei expressing EGFP–LAP2β on adhesive micropatterns of increasing diameter, shown as maximum z-projections (top row), midplane views (middle row), and side views (bottom row). As pattern diameter decreases, cell spreading is restricted and nuclei display rounder envelopes with more pronounced folds. **E.** Quantification of the fraction of high-curvature nuclear envelope regions (nuclear folds) as a function of adhesive micropattern diameter in HeLa cells (*N=3* independent experiments*; n= 72,71,72,77,70* nuclei for *10, 20, 30, 35 and 40 µm* pattern diameters*;* Supplementary Information, Section III.J). Black squares indicate the mean, and the shaded region indicates the standard deviation across nuclei.

**F–H,** Time-dependent evolution in nuclear condensate number and radius following confinement to 4 µm. **F.** Representative images of HeLa nuclei expressing DAXX–emGFP, with labelled condensates (Garcia-Jove Navarro *et al.*, 2019). **G.** Nuclear condensate number as a function of time after confinement (*N=2, n =16*). **H.** Condensate radius as a function of time after confinement (*N=2, n =16*). Black squares indicate the mean, and the shaded region indicate the standard deviation.

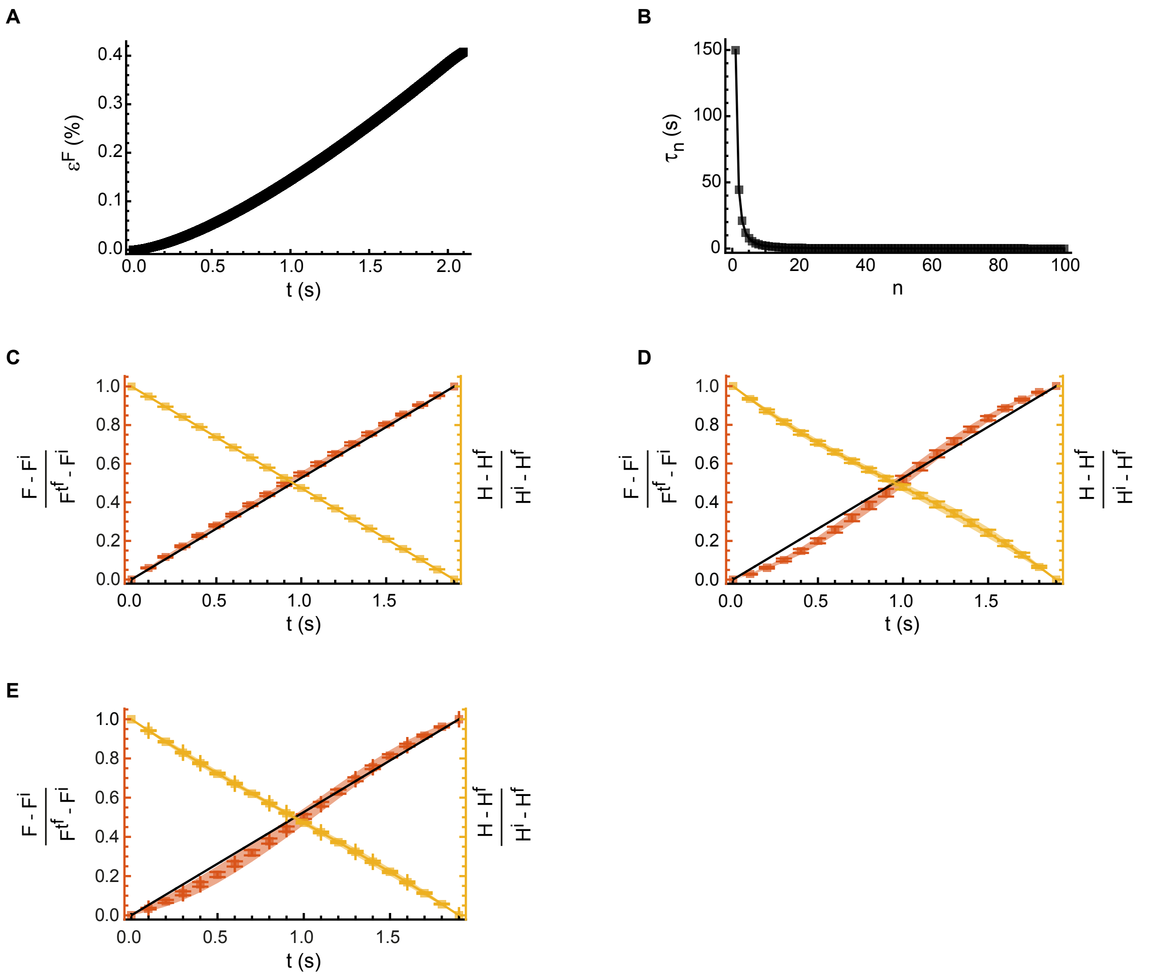

2026_Rollin et al. Sup. Fig5

**Supplementary Figure 5:**

**A.** Relative error in the predicted compression force between the full analytical solution and the limiting regime of no nuclear volume loss (see Supplementary Information, Section II.G). Parameters were determined self-consistently from the inference pipeline shown in Fig. 6.

**B.** Predicted volume and force relaxation timescale $\tau_n$ as a function of deformation mode n (see Supplementary information, Section. II.E).

**C-E.** Force and confinement height during the 4-to-3 μm compression step for two representative siCTL–LatA replicates (**C**, n = 7; **D**, n = 12) and averaged across all replicates (**E**; N = 10 independent experiments, n = 45 cells). In all plots, square points and whiskers indicate the mean and 95% confidence interval, respectively, while shaded regions indicate the standard deviation.

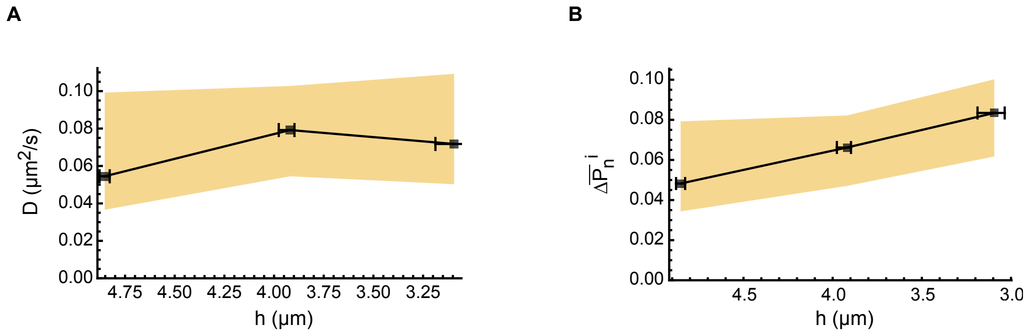

2026_Rollin et al. Sup. Fig6

**Supplementary Figure 6:**

**A,B.** Additional parameters inferred from the dynamic model-fitting pipeline for Latrunculin A treated HeLa cells (*N=3* independent experiments*, n=22* cells; Supplementary Information, Section II.G). Squares indicate the median, and error bars represent the 95% confidence interval obtained by bootstrapping the median $10^4$ times (see Supplementary Information, Section III.L).

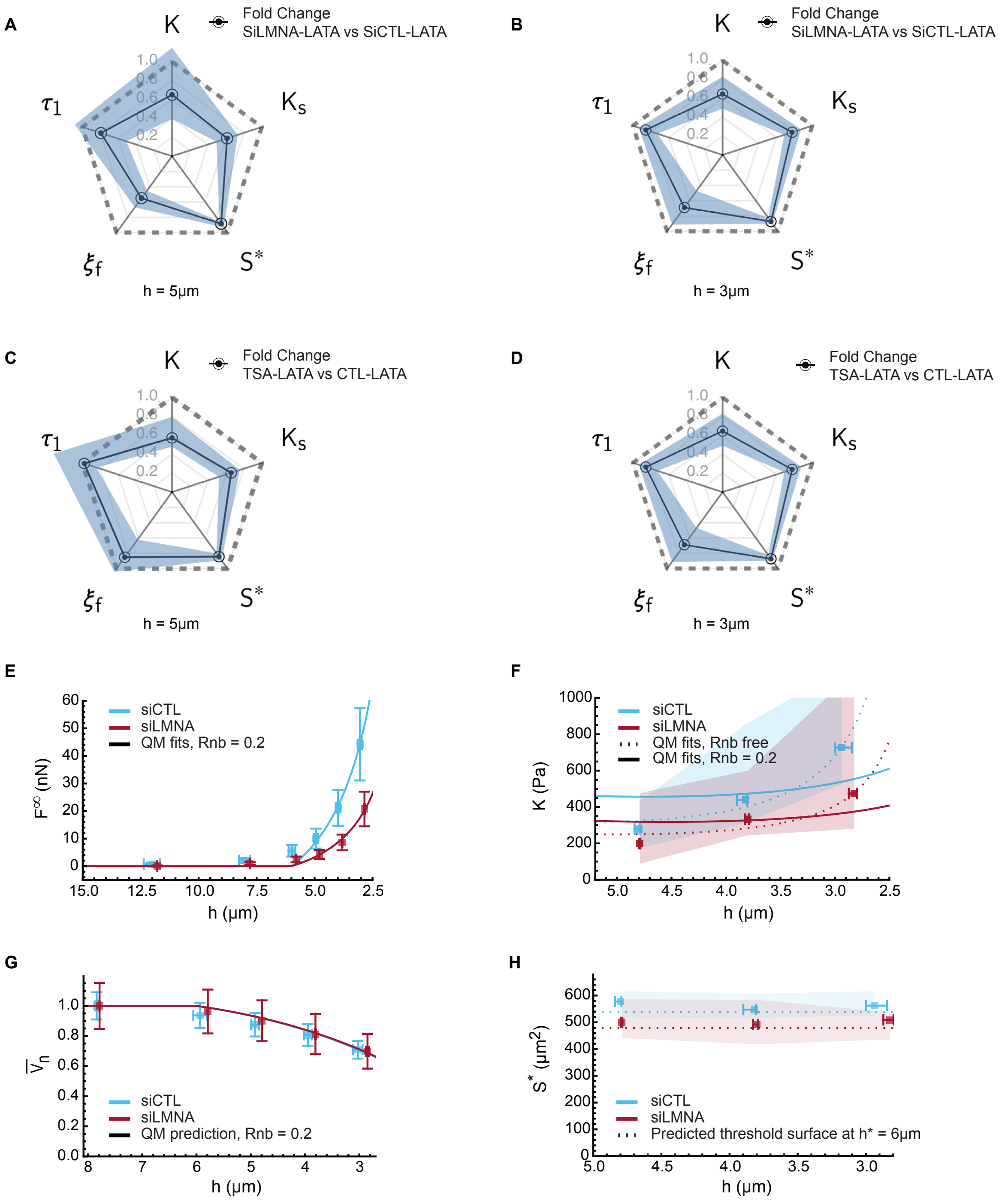

2026_Rollin et al. Sup. Fig7A

133
134

**Supplementary Figure 7 A:**

Unless otherwise indicated, all cells were treated with Latrunculin A (incubated 20 minutes before experiments).

**A–D.** Fold-changes in inferred model parameters for the 6–5 μm (A, C) and 5–4 μm (B, D) confinement steps under different perturbations. **A,B.** Lamin A/C depletion via siRNA (siLMNA; *N=7,n=32*) relative to the non-targeting control (siCTL; *N=10, n=45*). **C,D.** Trichostatin A (TSA; *N=3, n = 22*) treatment relative to control (CTL; N=6, n = 37) in HeLa cell nuclei. Circles denote the population median, and error bars represent the 50% confidence interval (CI) obtained by bootstrapping the median times (Supplementary Information Section III.L).

**E–H.** Quasistatic model (QM) validation for the AFM data. **E, F.** The model was fitted to the post-relaxation force and the bulk modulus, as inferred from the dynamic model (Supplementary Figure 7C,D), for siCTL and siLMNA HeLa nuclei (Supplementary Information, Section I.H). **G.** Predicted normalized nuclear volume across successive confinement steps. Volumes are normalized to the baseline volume measured at the initial confinement height. **H.** Comparison of the threshold surface inferred from the dynamic model with that predicted by the quasistatic model, assuming a threshold height of 6 μm. For raw data (force and volume), squares and whiskers denote the mean ± 95% CI. For inferred parameters (bulk and threshold surface), squares denote the median and error bars indicate the 95% CI via bootstrap resampling.

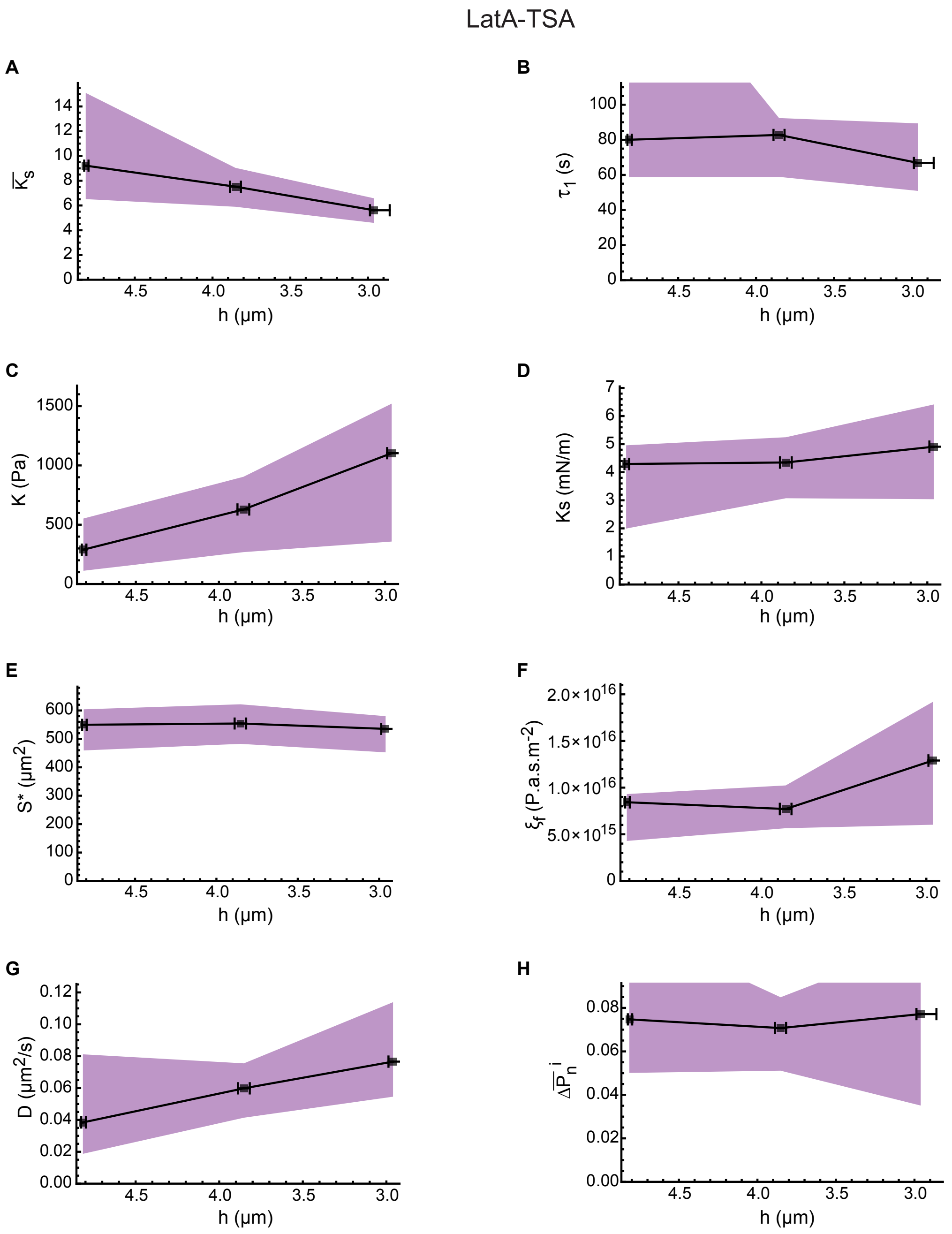

2026_Rollin et al. Sup. Fig7B

siCTL-LatA

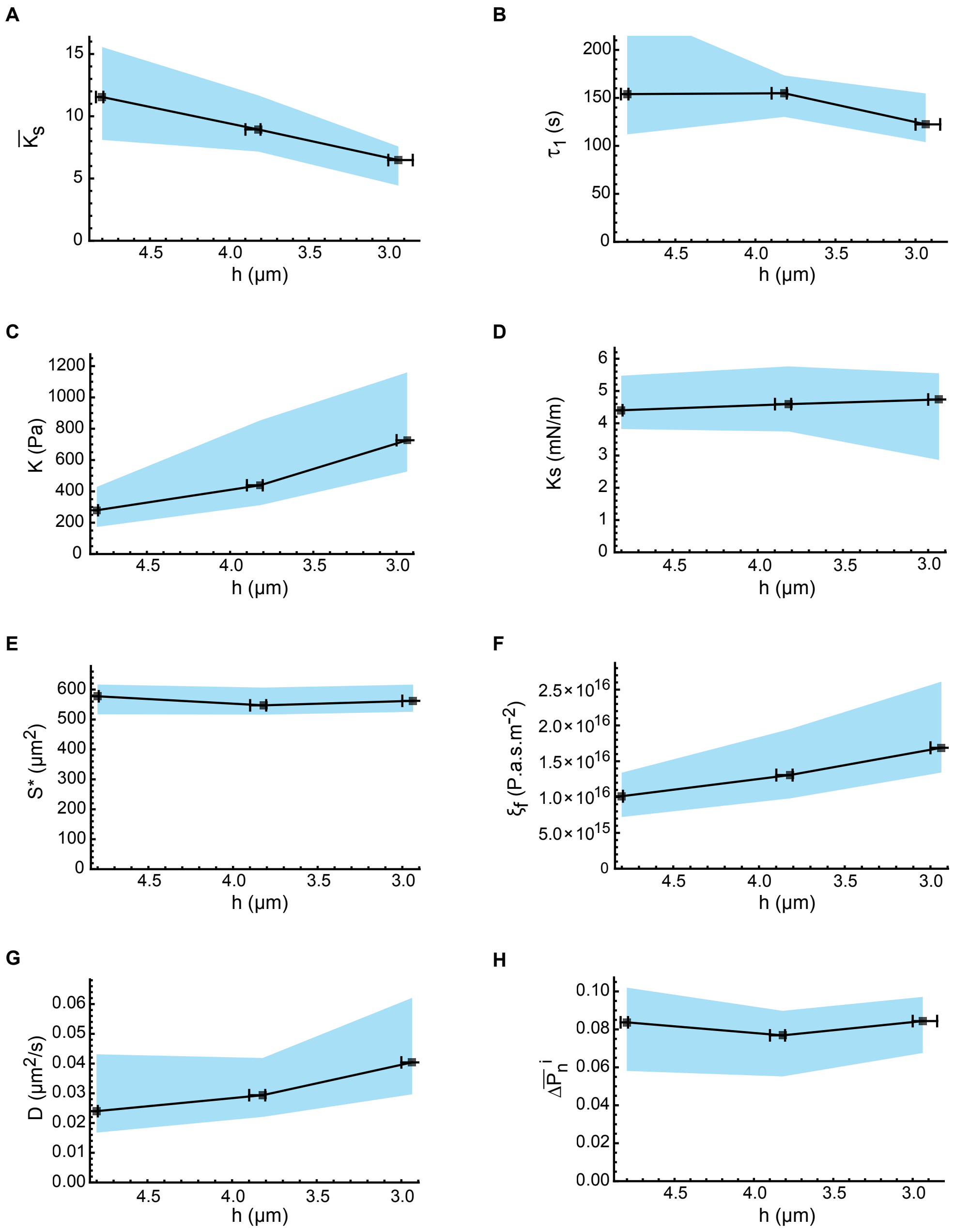

2026_Rollin et al. Sup. Fig7C

siLMNA-LatA

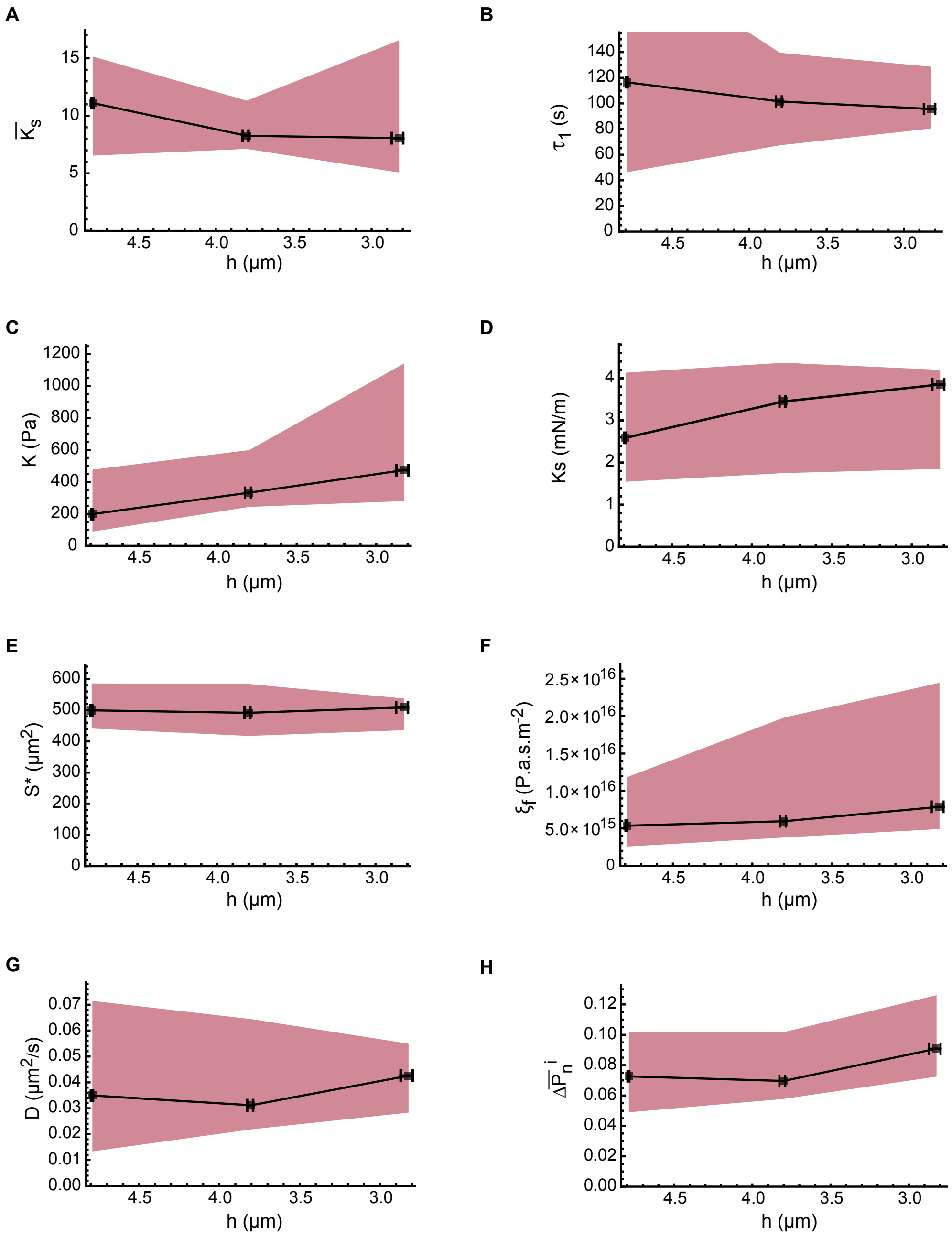

2026_Rollin et al. Sup. Fig7D

**Supplementary Figure 7 B, C, D:**

Inferred model parameters, defined in Fig. 5B, across successive confinement steps for HeLa cell nuclei treated with LatA–TSA (Supplementary Fig. 7B; $N = 3$ independent experiments, $n = 22$ nuclei), siCTL–LatA (Supplementary Fig. 7C; $N = 10$, $n = 45$), or siLMNA–LatA (Supplementary Fig. 7D; $N = 7$, $n = 32$). Treatments are defined as follows: LatA–TSA, Latrunculin A combined with trichostatin A; siCTL–LatA, control siRNA combined with Latrunculin A; siLMNA–LatA, LMNA/C knockdown combined with Latrunculin A. Black squares indicate the population median; whiskers and shaded regions indicate 95% confidence intervals for the x- and y-errors, respectively. Error margins were estimated by bootstrapping the median $10^4$ times (Supplementary Information, Section III.L). Dimensional parameters were obtained by combining experimental measurements of the relaxed midplane nuclear surface area with fitted dimensionless parameter values (Supplementary Information, Section II.G).
