## Supplemental model and methods for "Coupled nucleoplasmic flows and envelope expansion govern the mechanical resistance of the nucleus"

#### I. QUASISTATIC MODEL OF NUCLEAR VOLUME LOSS UPON UNIAXIAL CONFINEMENT.

We adapt in this section the nested pump-leak model that we proposed in ([1]) to describe nuclear volume based on three generic physical constraints: balance of water chemical potential at the nuclear envelope, balance of small osmolytes fluxes, and electroneutrality.

##### A. Minimal osmotic model of the nucleus

We and others showed that significant volume changes due to external perturbations require stresses comparable to the osmotic pressure exerted by trapped osmolytes — either sterically confined or retained to maintain electroneutrality [1]. Typical cellular osmolarity is on the order of several hundreds of millimolars, which corresponds to an osmotic pressure ($k_BT \cdot c$) of about $10^6$Pa at room temperature (where c is the total solute concentration and $k_BT$ is the thermal energy). This osmotic pressure is two to three orders of magnitude lower than the Laplace pressure generated by cortical tension, explaining why changes in plasma membrane tension have a negligible direct effect on cell volume. In contrast, here we hypothesize that Laplace pressure at the nuclear envelope (NE) in the taut regime cannot be neglected. This is due to the high permeability of nuclear pores, which allow the free diffusion of many small osmolytes through the NE. Our previous quantitative estimates support this view, showing that permeable osmolytes—such as ions and metabolites—are present at concentrations on the order of 100 mMol, whereas impermeable macromolecules, such as proteins, are found in the mMol range. Accordingly, our model distinguishes between two classes of osmolytes :

- Trapped osmolytes are confined within the nucleus and cannot freely diffuse through nuclear pores, but are free to move within the nuclear compartment. These include large proteins and DNA counterions necessary for maintaining electroneutrality. We denote their numbers as Xn in the nucleus and Xc in the cytoplasm.
- Untrapped osmolytes can freely diffuse through nuclear pores but cannot escape the cell. These include smaller molecules such as ions and amino acids. We denote their numbers as Mn and Mc in the nucleus and cytoplasm, respectively.

The balance of water chemical potential, .i.e. balance of osmotic and hydrostatic pressure difference at the nuclear envelope and plasma membrane, lead to an expression for the nucleus and cytoplasmic volumes of the form:

$$V_n = R_n + \frac{1}{\beta \Pi_0} \cdot \frac{X_n + M_n(\frac{\Delta P_n}{\Pi_0})}{1 + \frac{\Delta P_n}{\Pi_0}} \tag{1}$$

$$V_c = R_c + \frac{X_c + M_c}{\beta \Pi_0} \tag{2}$$

where, we neglected the role of the laplace pressure at the plasma membrane in $V_c$. $R_n$ is the osmotically inactive volume in the nucleus (also called dry volume or excluded volume). $\beta = \frac{1}{kT}$ is the inverse of the thermal energy and $\Delta P_n$ the hydrostatic pressure difference at the nuclear envelope. We highlight that $\frac{\Delta P_n}{\Pi_0}$ is a small parameter as $\Delta P_n \sim 10^2 Pa$ and $\Pi_0 \sim 10^5 - 10^6 Pa$ ([1]). Yet, it cannot be neglected as it impacts the number of small osmolytes in the nucleus $M_n(\frac{\Delta P_n}{\Pi_0})$. Moreover, for simplicity, and due to limited knowledge of the electric potential across the nuclear envelope, we assume that freely diffusing species equilibrate their concentrations across the nuclear envelope on the timescale of interest in this section (tens of minutes):

$$\frac{M_n}{V_n - R_n} = \frac{M_c}{V_c - R_c} \tag{3}$$

Finally, we denote by $M_t$ the total number of small osmolytes in the cell, and assume that these osmolytes neither leak out of the cell nor are significantly produced during the confinement experiments, given the short timescale of a few minutes under consideration.

$$M_t = M_c + M_n \tag{4}$$

##### B. Nuclear envelope tension

The Laplace pressure at the nuclear envelope is related to the tension and the mean curvature of the free NE surface (not in contact with the PDMS plates) through the Laplace law: $\Delta P_n \sim 2 \cdot \gamma \cdot \mathcal{C}$. Thus, to predict the influence of confinement on nuclear volume we need to assume a constitutive relation between the nuclear surface deformation and the tension build-up at the NE.

$$\gamma = \begin{cases} 0 \quad \text{for} \quad S \leq S^* \\ K_s \cdot \left(\frac{S}{S^*} - 1\right) \quad \text{for} \quad S \geq S^* \end{cases} \tag{5}$$

where, $S$ is the total surface of the nucleus, $S^*$ is the surface at which the NE becomes taut (no more folds opening upon deformation), and $K_s$ is the elastic stretching modulus of the surface. This non-linear elastic constitutive relation of the NE was initially motivated by hypo-osmotic shocks experiments conducted on articular chondrocytes nuclei and more recent findings showing that nuclear wrinkles unfold at constant volume upon rounding and spreading. We then quantitatively validated this behavior using atomic force microscopy (AFM) experiments (see Fig. 3).

##### C. Geometry of the nucleus upon uniaxial confinement.

###### *1. Force Balance.*

We base our derivation from the paper [2]. We assume that the geometry of the nucleus is axisymmetric upon compression, i.e., there is an invariance by rotation around the axis of compression. We can thus fully describe a point located on the surface of the nucleus by two coordinates: (r($\theta$),z($\theta$)). We call $\theta$ the angle between the tangent to the nucleus surface in the plane (Oxz) and the horizontal axis (see Supplementary Fig. 2A). For sake of simplicity, we will assume that $\theta$ varies between [0,$\pi$]. We next write the force balance on the dashed rectangle displayed in Fig. 2A to obtain a constraint on the shape of the nucleus surface. For simplicity, we neglect the influence of the cytoplasm in the force balance.

$$F + 2\pi \cdot r(\theta) \cdot \gamma \cdot sin(\theta) - \Delta P_n \cdot \pi r^2(\theta) = 0 \tag{6}$$

With, F the compressive force applied by the PDMS plate and $\Delta P_n$ the difference of hydrostatic pressures between the nucleus and the cytoplasm. The latter two quantities are not independent. Taking $\theta = 0$ in Eq.6 leads to the relation:

$$F = \Delta P_n \cdot \pi a^2 \tag{7}$$

Substituting Eq.7 in the force balance equation Eq.6 leads to a second order polynomial for r:

$$r^2 - \frac{2\gamma \cdot sin(\theta)}{\Delta P_n} \cdot r - a^2 = 0 \tag{8}$$

Which implies that:

$$r(\theta) = a \cdot \left(B \cdot sin(\theta) + \sqrt{1 + B^2 \cdot sin^2(\theta)}\right) \quad With, \quad B = \frac{1}{2 \cdot a \cdot \mathcal{C}} \tag{9}$$

Where, $\mathcal{C} = \frac{\Delta P_n}{2\gamma}$ is the average curvature at the edge of the nucleus. We introduce an important dimensionless parameter $B$ that naturally arises in the problem. It compares the average radius of curvature $\frac{1}{\mathcal{C}}$ at the edge of the nucleus with the radius of contact $a$ (see Supplementary Fig. 2A). $z(\theta)$ can then be deduced from $r(\theta)$ from the following geometrical constraint:

$$\frac{\mathrm{d}z}{\mathrm{d}r} = tan(\theta) \implies \frac{\mathrm{d}z}{\mathrm{d}\theta} = tan(\theta) \cdot \frac{\mathrm{d}r}{\mathrm{d}\theta} \tag{10}$$

By enforcing that $z(\theta = 0) = 0$, we further integrate Eq.10 between 0 and $\theta$ and obtain:

$$z(\theta) = a \cdot \left[B \cdot [1 - cos(\theta)] + E_1(\theta, -B^2) - E_2(\theta, -B^2)\right] \quad \text{With,} \quad \begin{cases} E_1 = \int_0^\theta \sqrt{1 + B^2 \cdot sin^2(\theta')}\,\mathrm{d}\theta' \\ E_2 = \int_0^\theta \frac{1}{\sqrt{1+B^2 \cdot sin^2(\theta')}}\,\mathrm{d}\theta' \end{cases} \tag{11}$$

Eq.11 is written according to two Euler incomplete elliptic integrals of the first and second kind. For clarity in the following, we simply denote by $E_1$ and $E_2$ the values of these integrals for $\theta = \frac{\pi}{2}$. We further take $\theta = \frac{\pi}{2}$ in Eq.11 to obtain the constraint relating $h$, $B$ and $a$:

$$\frac{h}{2} = a \cdot [B + E_1 - E_2] \tag{12}$$

We further use Eq.11 and Eq.9 to express the surface of the nucleus as a function of $a$ and $B$. We decompose the surface in two terms: the contact area with the two PDMS plates $2\pi a^2$, and the free surface (in contact with the cytoplasm) $S^e$. We express the latter area as:

$$\frac{S^e}{2} = \int_0^{\frac{\pi}{2}} 2\pi r\,\mathrm{d}s = 2\pi \cdot (aB)^2 \left[2 + \frac{2}{B} \cdot E_1 - \frac{1}{B} \cdot E_2\right] \tag{13}$$

Such that the total surface of the nucleus $S$ and the midplane surface $S^{proj}$ read:

$$S = S^e + 2\pi a^2 = 4\pi(aB)^2 \left[2 + \frac{2}{B} \cdot E_1 - \frac{1}{B} \cdot E_2\right] + 2\pi a^2 \tag{14}$$

$$S^{proj} = \pi \cdot r^2(\theta = \frac{\pi}{2}) = \pi \cdot a^2 \cdot \left(B + \sqrt{1 + B^2}\right)^2 \tag{15}$$

Finally, following the same reasoning, the nucleus volume can be expressed as:

$$V_n^{(2)} = \int_0^{\frac{\pi}{2}} \pi r^2\,\mathrm{d}z = \frac{2\pi}{3} \cdot (aB)^3 \cdot \left[\left(\frac{3}{B^2} + 8\right) + \frac{1}{B} \cdot \left(8 + \frac{1}{B^2}\right) \cdot E_1 - \frac{1}{B} \cdot \left(4 + \frac{1}{B^2}\right) \cdot E_2\right] \tag{16}$$

##### *2. Summary of key geometrical equations.*

$$\frac{h}{2} = a \cdot [B + E_1 - E_2] \tag{17}$$

$$V_n^{(2)} = \frac{2\pi}{3} \cdot (aB)^3 \cdot \left[\left(\frac{3}{B^2} + 8\right) + \frac{1}{B} \cdot \left(8 + \frac{1}{B^2}\right) \cdot E_1 - \frac{1}{B} \cdot \left(4 + \frac{1}{B^2}\right) \cdot E_2\right] \tag{18}$$

$$S = S^e + 2\pi a^2 = 4\pi(aB)^2 \left[2 + \frac{2}{B} \cdot E_1 - \frac{1}{B} \cdot E_2\right] + 2\pi a^2 \tag{19}$$

$$S^{proj} = \pi \cdot a^2 \cdot \left(B + \sqrt{1 + B^2}\right)^2 \tag{20}$$

##### *3. Asymptotic regimes of nuclear shape*

*a. For $B << 1$,* the nucleus takes the shape of a pancake and the nuclear volume $V_n$, surface $S$ and mean curvature of the side surface $\mathcal{C}$ equations reduce to :

$$V_n \approx \pi a^2 h \quad , \quad S \approx 2\pi a^2 + 2\pi a h \quad , \quad \mathcal{C} \approx \frac{1}{2} \cdot \left(\frac{2}{h}\right) \tag{21}$$

*b. For $B >> 1$,* the nucleus takes the shape of a sphere and the nuclear volume $V_n$, surface $S$ and mean curvature of the edges $\mathcal{C}$ equations reduce to :

$$V_n \approx \frac{4}{3}\pi \left(\frac{h}{2}\right)^3 \quad , \quad S \approx 4\pi \left(\frac{h}{2}\right)^2 \quad , \quad \mathcal{C} \approx \frac{2}{h} \tag{22}$$

#### D. Analytical solution close to the volume loss threshold.

##### *1. Normalization.*

It is convenient to normalize the dimensional parameters of the model to simplify the subsequent analytical and numerical resolutions. We choose the following conventions for the normalization of the model parameters :

- Pressures are normalized by the external osmotic pressure $\Pi_0$.
- Osmolyte numbers (.i.e. proteins, metabolites etc) are normalized by the number of small osmolytes in the nucleus before confinement $M_n^0$.
- Volumes by the nuclear volume before confinement $V_n^0$, Areas by $V_n^{0\,\frac{2}{3}}$, and Lengths by $V_n^{0\,\frac{1}{3}}$.
- Tensions by $\Pi_0 \cdot V_n^{0\,\frac{1}{3}}$

##### *2. Summary of model's equations in normalized form.*

The nuclear volume expression arising from the osmotic constraint reads:

$$\overline{V}_n = \overline{R}_n + (1 - \overline{R}_n) \cdot \frac{1 - \frac{1 - \overline{M}_n}{1 + \overline{X}_n}}{1 + \overline{\Delta P_n}} \tag{23}$$

With,

$$\overline{\Delta P_n} = 2 \cdot \overline{\gamma} \cdot \overline{\mathcal{C}} \quad , \quad \overline{\mathcal{C}} = \frac{1}{2 \cdot \overline{a} \cdot B} \quad , \quad \overline{\gamma} = \begin{cases} 0 \quad \text{for} \quad \overline{S} \leq \overline{S}^* \\ \overline{K_s} \cdot \left(\frac{\overline{S}}{\overline{S}^*} - 1\right) \quad \text{for} \quad \overline{S} \geq \overline{S}^* \end{cases} \tag{24}$$

And,

$$\overline{S} = 4\pi(\overline{a}\cdot B)^2\left[2+\frac{2}{B}\cdot E_1-\frac{1}{B}\cdot E_2\right]+2\pi\overline{a}^2 \tag{25}$$

The expression of nuclear volume coming from the force balance reads:

$$\overline{V_n}^{(2)} = \frac{2\pi}{3}\cdot(\overline{a}B)^3\cdot\left[\left(\frac{3}{B^2}+8\right)+\frac{1}{B}\cdot\left(8+\frac{1}{B^2}\right)\cdot E_1-\frac{1}{B}\cdot\left(4+\frac{1}{B^2}\right)\cdot E_2\right] \tag{26}$$

With,

$$\overline{a} = \frac{\overline{h}}{2\cdot[B+E_1-E_2]} \tag{27}$$

The cytoplasmic volume reads:

$$\overline{V}_c = \overline{R}_c + (1-\overline{R}_n)\cdot\frac{\overline{X}_c+\overline{M}_t-\overline{M}_n}{1+\overline{X}_n} \tag{28}$$

To solve this non-linear system of equations, we first solve $\overline{M_n}$ analytically using Eq.3, see below, and then numerically equate Eq.26 and Eq.23 to self-coherently determine $B$ using the Mathematica software implementation of the Newton-Raphson method.

##### 3. *Discussion on the parameters of the model.*

*a. Expression of $\overline{X}_c$ and $\overline{M}_t$.* Before confinement, nuclei must comply with Eq.1 with $\Delta P_n = 0$. This imposes constraints on the model's parameter that we detail below. We have :

$$\frac{M_n^0}{V_n^0-R_n^0} = \frac{M_c^0}{V_c^0-R_c^0} \quad \text{And,} \quad \frac{X_n}{V_n^0-R_n^0} = \frac{X_c}{V_c^0-R_c^0} \tag{29}$$

where we assumed that, on the timescale of a few minutes, proteins do not significantly redistribute between the cytoplasm and the nucleus ($X_n \approx X_n^0$). By further calling $NC^0 = \frac{V_n^0-R_n^0}{V_c^0-R_c^0}$, we have :

$$\overline{X}_c = \frac{\overline{X}_n}{NC^0} \tag{30}$$

$$\overline{M}_t = 1+\frac{1}{NC^0} \tag{31}$$

*b. Expression of $\overline{X}_n$* Before confinement, the nuclear envelope is not taut such that the Laplace pressure is negligible and there is osmotic balance at the nuclear envelope. Because the osmotic pressure in the cytoplasm is with a very good accuracy equal to the external osmotic pressure ([1]) we can write:

$$\Pi_0 = k_BT\cdot\left(x_n+m_n^0\right) \tag{32}$$

By calling $\Pi_t = k_BT\cdot x_n$ the osmotic pressure of the trapped nuclear osmolytes prior to the volume loss threshold we then express the parameter $\overline{X}_n$ as:

$$\overline{X}_n = \frac{X_n}{M_n^0} = \frac{x_n}{m_n^0} = \frac{\Pi_t}{\Pi_0-\Pi_t} \tag{33}$$

*c. Estimation of $NC^0$, $\overline{R}_n$ and $\overline{R}_c$.* Experimentally we measure an average HeLa cell nuclear volume of $V_n^0 \approx 870\mu\text{m}^3$ (see Fig. 1C). We measured in previous studies a total average HeLa cell volume of $V^0 = 2160\mu\text{m}^3$ ([3]) which imply an average cytoplasmic volume of $V_c^0 \approx 1290\mu\text{m}^3$ before confinement. We showed doing osmotic shock experiments on HeLa cells that $R_{tot} \sim R_n + R_c$ was about 10 to 30% of the whole cell volume. Calling $\alpha_n = \frac{R_n}{V_n^0}$, $\alpha_c = \frac{R_c}{V_c^0}$ and $\alpha_{tot} = \frac{R_{tot}}{V_{tot}}$ we must have :

$$(\frac{\alpha_n}{\alpha_{tot}} - 1) \cdot V_n^0 + (\frac{\alpha_c}{\alpha_{tot}} - 1) \cdot V_c^0 = 0 \tag{34}$$

We do not have direct access to $\alpha_n$ nor $\alpha_c$ in our experiments. For simplicity, we take the simplest solution of the previous equation, .i.e. $\alpha_n = \alpha_c = \alpha_{tot} = 0.2$ but we checked that modifying the value of $\alpha_n$ and $\alpha_c$ does not change our conclusions. We summarize all the parameter values used to fit the quasistatic model to the data in Tab.I.

##### *4. Resolution.*

*a. Permeable osmolyte number in the nucleus* Injecting Eq.1 into 3 we obtain a second order polynomial to solve in $\overline{M}_n$, with one solution lower than one:

$$\overline{M}_n = \frac{1}{2} \cdot \left( \overline{M}_t \cdot (1 + \frac{\overline{X}_n}{\overline{\Delta P}_n}) + \overline{X}_c - \sqrt{\left( \overline{M}_t \cdot (1 + \frac{\overline{X}_n}{\overline{\Delta P}_n}) + \overline{X}_c \right)^2 - 4 \cdot \frac{\overline{X}_n}{\overline{\Delta P}_n} \cdot \overline{M}_t} \right) \tag{35}$$

*b. Analytical resolution close to the threshold and pancake shape.* As $\overline{M}_n$ depends on the parameter $\epsilon = \frac{\overline{\Delta P}_n}{\overline{X}_n}$ and because this parameter is small close to the tension threshold we Taylor expand all the variables of our problem with respect to $\epsilon$:

$$\overline{M}_n = 1 - a_1 \cdot \epsilon \quad \overline{V}_n = 1 - a_2 \cdot \epsilon \quad \overline{h} = \overline{h}^* - h_1 \cdot \epsilon \quad \overline{S} = \overline{S}^* + a_3 \cdot \epsilon \quad \overline{\Delta P}_n = a_4 \cdot \epsilon = \overline{X}_n \cdot \epsilon \tag{36}$$

where, $a_1, a_2, a_3, a_4, h_1$ are constants to determine according to the parameters of the problem and the sign in front of them is chosen so that these constants are positive. By assuming that the nuclei have a pancake shape, we find :

$$a_1 = 1 + \frac{\overline{X}_c - 1}{\overline{M}_t} = \frac{1 + \overline{X}_n}{1 + NC} \tag{37}$$

$$a_2 = \frac{(1 - \overline{R}_n) \cdot (1 + \overline{X}_n \cdot (1 + NC))}{1 + NC} \tag{38}$$

$$a_3 = h_1 \cdot f(\overline{h}^*) - a_2 \cdot g(\overline{h}^*) \quad \text{with,} \quad f(\overline{h}^*) = \frac{2}{\overline{h}^{*2}} - \sqrt{\frac{\pi}{\overline{h}^*}} \quad \text{and,} \quad g(\overline{h}^*) = \frac{2}{\overline{h}^*} + \sqrt{\pi \overline{h}^*} \tag{39}$$

$$a_4 = \frac{a_3 \cdot \overline{K}_s}{q(\overline{h}^*)} \quad \text{with,} \quad q(\overline{h}^*) = 1 + \overline{h}^{*3/2} \cdot \sqrt{\pi} \tag{40}$$

$$\epsilon = \frac{\overline{h}^*}{h_1} \cdot (1 - \frac{\overline{h}}{\overline{h}^*}) \tag{41}$$

Finally, we determine $h_1$ by imposing that by definition $a_4 = \overline{X}_n$ :

$$h_1 = \frac{f(\overline{h}^*)}{g(\overline{h}^*)} \cdot a_2 + \frac{q(\overline{h}^*)}{f(\overline{h}^*)} \cdot \frac{\overline{X}_n}{\overline{K}_s} \tag{42}$$

*c. Solutions.* The solution for the nuclear volume surface, small osmolytes and difference of pressure thus read:

$$\overline{V}_n = 1 - v_1 \cdot (1 - \frac{\overline{h}}{\overline{h}^*}) \quad \text{with,} \quad v_1 = \frac{a_2 \cdot \overline{h}^*}{h_1} = \frac{v_1^{lim}}{1 + \frac{q(\overline{h}^*)}{g(\overline{h}^*)} \cdot \frac{1}{a_2} \cdot \frac{\overline{X}_n}{\overline{K}_s}} \quad \text{and,} \quad v_1^{lim} = \overline{h}^* \cdot \frac{f(\overline{h}^*)}{g(\overline{h}^*)} = \frac{2 - \sqrt{\pi} \cdot \overline{h}^{*3/2}}{2 + \sqrt{\pi} \cdot \overline{h}^{*3/2}} \tag{43}$$

$$\overline{S} = \overline{S}^* + \frac{a_3 \cdot \overline{h}^*}{h_1} \cdot (1 - \frac{\overline{h}}{\overline{h}^*}) = \overline{S}^* + \overline{h}^* \cdot f(\overline{h}^*) \cdot (1 - \frac{v_1}{v_1^{lim}}) \cdot (1 - \frac{\overline{h}}{\overline{h}^*}) \tag{44}$$

$$\overline{M}_n = 1 - \frac{a_1 \cdot \overline{h}^*}{h_1} \cdot (1 - \frac{\overline{h}}{\overline{h}^*}) \tag{45}$$

$$\overline{\Delta P}_n = \frac{\overline{X}_n \cdot \overline{h}^*}{h_1} \cdot (1 - \frac{\overline{h}}{\overline{h}^*}) \tag{46}$$

*d. Limit of stiff nuclear envelope - deformation at constant surface* An important limit is when the parameter $v_1 = v_1^{lim}$, this corresponds to a regime where the nucleus loses volume at constant surface upon confinement. According to our estimates the parameter that makes $v_1$ close to $v_1^{lim}$ in our experiments is $\frac{\overline{X}_n}{\overline{K}_s}$ which must be small to fit our data (Fig. 2C). We estimated $\overline{X}_n \approx 1.6 \cdot 10^{-3}$ from literature values (see Table.I). In this experimental limit $\frac{\overline{X}_n}{\overline{K}_s}$ can be approximated by the following expression:

$$\frac{\overline{X}_n}{\overline{K}_s} \sim \frac{\Pi_t \cdot V_n^{0\,\frac{1}{3}}}{K_s} \tag{47}$$

With, $\Pi_n^{0,p}$ the nuclear osmotic pressure of trapped nuclear osmolytes before confinement. Thus in the experimental limit and close to the tension increase threshold, the parameter that is critical for volume loss is $\frac{\Pi_t \cdot V_n^{0\,\frac{1}{3}}}{K_s}$. Based on our data analysis pipeline (Section II G), we find for LatA HeLa Cells ($K_s \sim 10\,\mathrm{mN/m}$, $\Pi_t \sim 200\,\mathrm{Pa}$, $V_n^0 \sim 1000\,\mu\mathrm{m}^3$). We thus verify that $\Pi_t < \frac{K_s}{V_n^{0\,\frac{1}{3}}} \sim 1000Pa$ which confirms that we are in the stiff regime.

##### E. Bulk modulus of the nucleus

*a. Definition* We define the bulk modulus of the nucleus as the inverse of the classical compressibility modulus.

$$\frac{1}{K} = -\frac{1}{V_n} \cdot \frac{\partial V_n}{\partial P_n} \tag{48}$$

This modulus tells how much the volume will change upon application of a small increment of pressure inside the nucleus:

$$\frac{\delta V_n}{V_n} = -\frac{1}{K} \cdot \delta P_n \tag{49}$$

Note that since in our problem the change of volume of the nucleus is due to the movement of water, the bulk modulus is of osmotic origin, .i.e. related to the chemical potential of water.

*b. Numerical computation for LatA treated HeLa Cells.* Our quasistatic model allows us to predict the evolution of $\overline{V}_n$ and $\overline{\Delta P}_n$ according to $\overline{h}$. The normalized nuclear bulk modulus $\overline{K} = \frac{K}{\Pi_0}$ reads :

$$\overline{K}(\overline{h}) = -\overline{V_n} \cdot \frac{1}{\frac{\partial \overline{V}_n}{\partial \overline{P}_n}} \tag{50}$$

Using chain rule we express $\frac{\partial \overline{V}_n}{\partial \overline{P}_n}$ as:

$$\frac{\partial \overline{V}_n}{\partial \overline{P}_n} = \frac{1}{\frac{\partial \overline{P}_n}{\partial \overline{h}}} \cdot \frac{\partial \overline{V}_n}{\partial \overline{h}} \tag{51}$$

We then express the nuclear pressure as the sum of the cytoplasmic pressure $P_c$ and the nucleus-cytoplasm difference of pressure $\Delta P_n$ :

$$\overline{P}_n = \overline{P}_c + \overline{\Delta P}_n \tag{52}$$

The Laplace pressure at the plasma membrane reads:

$$\overline{P}_c = \overline{P}_0 + \frac{2\overline{\gamma_c^0}}{\overline{h}} \tag{53}$$

Where, $\overline{\gamma_c^0}$ denotes the normalized plasma membrane tension. We use the superscript $^0$ to indicate our assumption that the plasma membrane tension remains constant during confinement after LatA treatment. This assumption is motivated by the presence of membrane reservoirs in the plasma membrane and by the fact that LatA abolishes cellular contractility. Moreover, we assumed that the mean curvature of the lateral side of the plasma membrane, $\mathcal{C}_c$, satisfies $\mathcal{C}_c = \frac{2}{h}$. This approximation is justified in the strongly confined regime, where the cell's contact radius is much larger than the confinement height. In this limit, the cell side edges are circles of radius $\frac{h}{2}$. Finally, $\overline{P}_0$ denotes the pressure of the external medium. Under these assumptions we have:

$$\overline{P}_n = \overline{P}_0 + \frac{2\overline{\gamma_c^0}}{\overline{h}} + \overline{\Delta P}_n \tag{54}$$

Which can be differentiated numerically to predict $\frac{\partial \overline{P}_n}{\partial \overline{h}}$ and thus $\overline{K}(\overline{h})$.

*c. Analytical approximation for LatA treated HeLa Cells at the threshold.* We now leverage the analytical solution derived in the previous section in the vicinity of the height threshold $h^*$ to elucidate how the bulk modulus $\overline{K}$ depends on the model parameters. Since $\overline{V}_n$ and the nuclear pressure difference $\overline{\Delta P_n}$ were expanded to linear order in $\overline{h}$, taking a derivative with respect to $\overline{h}$ to compute $\overline{K}$ reduces the accuracy of the approximation to the zeroth order in $\frac{\overline{h}-\overline{h}^*}{\overline{h}^*}$. In the remainder of this paragraph, we will therefore compute $\overline{K}(\overline{h}^*)$. For clarity, we recall here the main result of the previous computation. We showed that :

$$\overline{V}_n = 1 - v_1 \cdot \left(1 - \frac{\overline{h}}{\overline{h}^*}\right) \quad \text{with,} \quad v_1 = \frac{a_2 \cdot \overline{h}^*}{h_1} \tag{55}$$

$$\overline{\Delta P}_n = \frac{\overline{X}_n \cdot \overline{h}^*}{h_1} \cdot (1 - \frac{\overline{h}}{\overline{h}^*}) \tag{56}$$

We first expand the cytoplasmic pressure. By using, $\overline{h} = \overline{h}^* \cdot \left(1 + \frac{\overline{h}-\overline{h}^*}{\overline{h}^*}\right)$ and Laplace law, we have at linear order in $\frac{\overline{h}-\overline{h}^*}{\overline{h}^*}$ :

$$\overline{P}_c = \overline{P}_0 + \overline{\Delta P}_c^* \cdot \left(2 - \frac{\overline{h}}{\overline{h}^*}\right) \tag{57}$$

With, $\overline{\Delta P}_c^* = 2 \cdot \frac{\overline{\gamma}_c^0}{\overline{h}^*}$. Thus, the nuclear pressure becomes:

$$\overline{P}_n = \overline{P}_0 + \overline{\Delta P}_c^* \cdot \left(2 - \frac{\overline{h}}{\overline{h}^*}\right) + \frac{\overline{X}_n \cdot \overline{h}^*}{h_1} \cdot (1 - \frac{\overline{h}}{\overline{h}^*}) \tag{58}$$

And,

$$\frac{\partial \overline{P}_n}{\partial \overline{h}} = -\left(\frac{\overline{X}_n}{h_1} + \frac{\overline{\Delta P}_c^*}{\overline{h}^*}\right) \tag{59}$$

Finally, We have :

$$\frac{\partial \overline{V}_n}{\partial \overline{h}} = \frac{v_1}{\overline{h}^*} \tag{60}$$

Which leads to:

$$\frac{\partial \overline{V}_n}{\partial \overline{P}_n} = -\frac{1}{\left(\frac{\overline{X}_n}{h_1} + \frac{\overline{\Delta P}_c^*}{\overline{h}^*}\right)} \cdot \frac{v_1}{\overline{h}^*} \tag{61}$$

Thus the bulk modulus at the threshold reads:

$$\overline{K}(\overline{h}^*) = \frac{\overline{h}^*}{v_1} \cdot \left(\frac{\overline{X}_n}{h_1} + \frac{\overline{\Delta P}_c^*}{\overline{h}^*}\right) = \frac{\overline{X}_n}{a_2} \cdot \left(1 + \frac{h_1 \cdot \overline{\Delta P}_c^*}{\overline{X}_n \cdot \overline{h}^*}\right) \tag{62}$$

In the limit where the nuclear osmotic pressure of trapped osmolytes $\Pi_t$ is small compared to the external osmotic pressure $\Pi_0$, which corresponds to the experimental limit, the parameter $\frac{\overline{X}_n}{a_2}$ can be reexpressed as:

$$\frac{\overline{X}_n}{a_2} = \frac{1 + NC^0}{1 - \overline{R}_n} \cdot \frac{\Pi_t}{\Pi_0} \tag{63}$$

Which means that the bulk modulus of the nucleus scales with the osmotic pressure of trapped osmolytes :

$$K \sim \frac{1 + NC^0}{1 - \overline{R}_n} \cdot \Pi_t \tag{64}$$

##### F. Determination of the threshold surface in the approximated pancake-shape limit.

Below the threshold compression, the volume is constant. Thus :

$$\overline{V}_n = 1 \longrightarrow \overline{a} = \sqrt{\frac{1}{\pi \cdot \overline{h}}} \tag{65}$$

Thus, at the height threshold we have:

$$\overline{S}^* = 2 \cdot \left(\frac{1}{\overline{h}^*} + \sqrt{\pi} \cdot \sqrt{\overline{h}^*}\right) \tag{66}$$

We confirmed that the value of $\overline{S}^*$ inferred from our AFM data analysis pipeline (see Section. II G) was consistent with this estimate when assuming a height threshold of $h^* = 6\,\mu$m. Across all perturbations and confinement steps, the two estimates agreed within 10%, providing support for the robustness of our method (Fig. 7F; Supplementary Fig. 7A, H;).

##### G. Fitting procedure of the quasistatic model to the six-well confiner volume loss data

In Fig. 2C, we fitted the model predictions to the six-well confiner volume loss data shown in Fig. 1C. To reproduce the observed slope of volume loss, the model must operate in a regime in which volume decreases at nearly constant surface area, consistent with a stiff nuclear envelope and a low bulk modulus. In this limit, the data constrain only the ratio of the dimensionless parameters $\overline{X}_n/\overline{K}_s$, rather than their individual values. A fit to this data alone hence does not allow a meaningful parameter determination. We therefore asked whether standard literature values could nevertheless account for the observed volume loss magnitude through an increase in the nuclear envelope tension. The parameter values used, together with their literature support, are summarized in Table I.

TABLE I. Description and values of the fixed parameters used to compare experimental data to theoretical predictions in Fig. 2C.

| Symbol | Typical Value | Meaning |
|---|---|---|
| $V_n^0$ | $870\,\mu\mathrm{m}^3$ | Average nuclear volume before confinement for CTL HeLa cells (Fig. 1C). |
| $h^*$ | $6\,\mu\mathrm{m}$ | Height threshold at which volume loss occurs. |
| $K_s$ | $10\,\mathrm{mN/m}$ | NE stretching modulus. In the literature its value ranges from 1 to $20\,\mathrm{mN/m}$ typically ([4], [5]). |
| $\Pi_0$ | $6 \cdot 10^5$ Pa | Total osmotic pressure of the outer medium, corresponding to an external osmolarity of 300 mMol. |
| $V^0$ | $2160\,\mu\mathrm{m}^3$ | Average HeLa cell volume before confinement [3]. |
| $V_c^0$ | $\sim 1300\,\mu\mathrm{m}^3$ | Average HeLa cell cytoplasmic volume before confinement. |
| $NC^0$ | 0.7 | Nuclear volume to cytoplasmic volume ratio. |
| $\gamma_c^0$ | $10^{-4}\,\mathrm{mN/m}$ | Typical plasma membrane tension. It typically ranges between $10^{-5}$ to $10^{-3}\,\mathrm{mN/m}$ [6]. |
| $\overline{R}_n$ | 0.2 | Nuclear dry volume fraction before confinement. It typically ranges between 0.1 to 0.3. |
| $\overline{X}_n$ | $\approx 1.6 \cdot 10^{-3}$ | Ratio of trapped osmolyte versus external osmotic pressures. We use a concentration of trapped osmolytes $x_n = 0.5$ mMol based on [7] which corresponds to a colloid osmotic pressure of $\sim 1000$ Pa. |
| $kT$ | $4.1\,\mathrm{pN} \cdot \mathrm{nm}$ | Thermal energy. |

##### H. Fitting procedure of the quasistatic model to the AFM data.

In Fig. 7C–E and Supplementary Fig. 7.A, E-H, we fitted the quasistatic model to AFM data from TSA-LatA, LatA, SiCTL-LatA HeLa, and SiLMNA-LatA HeLa cells. For this, we used our AFM-based inference pipeline to determine $K_s$ (see Fig. 6F). We then explored two fitting strategies.

First, we fitted the three unknown model parameters, $(\overline{R}_n, \gamma_c^0, \overline{X}_n)$, to the $F^\infty$ and $K$ data, with all remaining parameters fixed to the values listed in Table I. This strategy reproduced the nonlinearity of K (Fig. 7D and Supplementary Fig. 7.A, F), but consistently returned $\overline{R}_n \sim 0.5$ for all perturbations. This value appears high compared with literature estimates for the whole cell (0.1–0.3). Although no direct nuclear measurement is available, we interpret such a large value as evidence of overfitting needed to capture the nonlinearity of K.

Because of this, we fixed $\overline{R}_n = 0.2$ and fitted only the two remaining parameters. Fits were performed by least-squares minimization using the Nelder–Mead implementation of NMinimize in Mathematica. The best-fit parameters are summarized in Table II. With this constrained strategy, the quasistatic model recovers the magnitude of K at confinements close to the threshold ($5\mu m$), but no longer reproduces the nonlinearity observed at high confinement, typically around ($3\mu m$). We interpret this as evidence for additional nonlinear contributions that are not included in the model. At very high confinement, higher-order osmotic effects, such as steric repulsion, may become important.

In addition, the decrease in $\overline{K}_s$ at high confinement suggests that chromatin may contribute an additional nonlinear component to the nuclear osmotic pressure.

TABLE II. Best fit parameter values used to compare AFM data to the quasistatic model for $\overline{R}_n = 0.2$.

| $\overline{X}_n$ | $\gamma_c^0$ (N/m) | **Condition** |
|---|---|---|
| $4.9 \cdot 10^{-4}$ | $5.0 \cdot 10^{-4}$ | LatA |
| $1.4 \cdot 10^{-4}$ | $5.0 \cdot 10^{-4}$ | TSA-LatA |
| $1.8 \cdot 10^{-4}$ | $2.5 \cdot 10^{-4}$ | SiCTL-LatA |
| $8.1 \cdot 10^{-5}$ | $2.3 \cdot 10^{-4}$ | SiLMNA-LatA |

#### II. A DYNAMICAL MODEL OF NUCLEAR VOLUME CHANGE UPON UNIAXIAL COMPRESSION.

##### 1. *The Nucleus as a 2 two-component linear poroelastic material.*

In this section, we summarize the closed set of equations governing the dynamics of a two-component linear poroelastic material. These equations originate from the foundational work of Biot and Terzaghi, originally developed to describe saturated soils ([8],[9]), and were later extended by Tanaka and Fillmore [10], who introduced the stress-diffusion coupling model to describe the behavior of polymer gels under osmotic shocks. This framework was subsequently refined by Doi [11].

$$\left\{\begin{array}{l} \xi^f \cdot (1-\phi_0) \cdot \left(\dot{\vec{u}} - \vec{q}^f\right) = \vec{\nabla} P^f \\ \vec{\nabla} \cdot \overline{\overline{\sigma}}^{el} = \vec{\nabla} P^f \\ \vec{\nabla} \cdot \vec{q} = 0 \end{array}\right. \qquad \text{With,} \quad \left\{\begin{array}{l} \overline{\overline{\sigma}}^{el} = \left(K \cdot \vec{\nabla} \cdot \vec{u}\right) \cdot \overline{\overline{\mathbb{1}}} + 2G \cdot \left(\overline{\overline{u}} - \left(\frac{1}{3} \vec{\nabla} \cdot \vec{u}\right) \cdot \overline{\overline{\mathbb{1}}}\right) \\ \overline{\overline{u}} = \frac{1}{2} \cdot \left[\left(\overline{\overline{\nabla}} \vec{u}\right) + \cdot \left(\overline{\overline{\nabla}} \vec{u}\right)^T\right] \\ \vec{q} = \phi_0 \cdot \dot{\vec{u}} + (1-\phi_0) \cdot \vec{q}^f \end{array}\right. \tag{67}$$

where, for clarity, we call $\phi_0 = n^s w^s$ the volume fraction of the dry content in the mesoscopic elements of volume $V_0$ (Fig. 5C). At linear order, $\phi_0$ is assumed constant during the deformation. K and G are the osmotic bulk and shear moduli of the dry content. $\vec{q}$ accounts for the displacement of the flux velocity which is scaling in the linear theory with the center of mass velocity of the mesoscopic volume elements. $\vec{u}$ and $\dot{\vec{u}}$ are the displacement and velocity of the dry component. $\xi^f$, $q^f$ and $P^f$ account for the fluid (superscript $^f$) friction, velocity and pressure.

##### 2. *Boundary conditions*

To fully solve the problem, we will need to impose boundary conditions. There exist two kinds of boundary conditions:

*a. Mechanical condition* The total stress tensor for this two-component system has two contributions: an osmo-elastic contribution arising from the dry-content within the nucleus (chromatin, proteins etc) and a pressure from the solvent : $\overline{\overline{\sigma}} = \overline{\overline{\sigma}}^{el} - P^f \cdot \mathbb{1}$. Then the force balance at the boundary gives the following condition:

$$\left(\overline{\overline{\sigma}}^{el} - P^f \cdot \mathbb{1}\right) \cdot \vec{n} = \vec{f}_{ext} \tag{68}$$

Where, $f_{ext}$ accounts for the force acting on the boundary surface of the gel and $\vec{n}$ the outward unit normal vector to the gel surface.

*b. Permeation condition* If the fluid permeates freely through the gel surface, the fluid pressure must be continuous with the external fluid pressure $P^f_{ext}$. If the fluid cannot permeate through the gel surface, the normal velocity of the network and of the fluid must be identical and from Darcy's law Eq.67 we obtain that the fluid pressure gradient normal to the boundary must cancel. We can consider intermediate regimes by adding a surface friction $\xi_s$ to describe the dynamic of flow through the boundary surface of the gel. All the regimes can be expressed mathematically as follows:

$$\left\{\begin{array}{ll} P^f = P^f_{ext} & \text{Permeable boundary} \\ \vec{\nabla} P^f \cdot \vec{n} = 0 & \text{Impermeable boundary} \\ P^f - P^f_{ext} = \xi_s \cdot \left(\vec{q}^f - \dot{\vec{u}}\right) \cdot \vec{n} & \text{Intermediate permeable boundary} \end{array}\right. \tag{69}$$

##### 3. *Adaptation of the stress-diffusion coupling model to the uniaxial-confinement experiments.*

*a. Validity of the linear model.* The equations in Eq.67 are linear and thus strictly valid only for small deformations, i.e., weak confinements. However, our experiments probe nuclear relaxation dynamics under confinement heights between 6 and 3 $\mu$m, which correspond to relatively large deformations ($50\% - 80\%$ strains relative to unconfined nuclear heights). As a result, applying the linear model over the full confinement range would not be valid, as nonlinear effects are expected to play a significant role. While a nonlinear model could in principle be used, we deliberately avoided this for two reasons. First, choosing a specific nonlinear form would require arbitrary assumptions about nuclear mechanics, given the limited information available. Second, such an approach would add unnecessary complexity to the analysis, given the relative simplicity of our experimental setup. To circumvent this limitation, we developed a stepwise AFM confinement protocol featuring successive small vertical displacements ($6 \rightarrow 5\mu$m$, 5 \rightarrow 4\mu$m$, 4 \rightarrow 3\mu$m), allowing the nucleus to relax to steady state between each step. Each step was analyzed independently, with model parameters allowed to vary. This strategy effectively probes the linear response around different confinement heights, enabling us to capture the nonlinear dependence of key mechanical parameters ($K(h^i), G(h^i), \xi^f(h^i), \phi_0(h^i)$, etc.) without relying on specific assumptions about their functional form (Fig. 6).

*b. Approximate geometry.* In the regime of volume loss (.i.e. of high confinement), we showed in Supplementary Methods, Sec.III I 2 that the nuclei could be well approximated by a pancake-like geometry. For readibility, we decompose the surface of the nucleus as the sum of the midplane surface: $S^{proj} = \pi b^2$ and the edge surface $S^e = 2\pi bh$. In this geometry, the approximate volume, surface and curvature of the nucleus read:

$$V_n \approx S^{proj} \cdot h \quad , \quad S \approx 2 \cdot S^{proj} + S^e \quad , \quad \mathcal{C} \approx \frac{1}{2} \cdot \left(\frac{2}{h} + \frac{1}{b}\right) \tag{70}$$

Note that the curvature is effectively assuming that the edges of the nucleus are circular. Moreover, we do not discard b in front of h because for a typical surface of $150\mu m^2$ we have that b=$7\mu m$ and h varies between 6 to $3\mu m$ in our experiments.

##### *4. Formulation of the problem.*

*a. Osmo-elastic stress.* We use the following constitutive equation to describe the osmo-elasticity of the nucleus:

$$\overline{\overline{\sigma}}^{el} = \overline{\overline{\sigma}}_i^{el} + \left(K - \frac{2}{3} \cdot G\right) \cdot \vec{\nabla} \cdot \vec{u} \cdot \overline{\overline{\mathbb{1}}} + G \cdot \left(\overline{\overline{\nabla}}\vec{u} + \left(\overline{\overline{\nabla}}\vec{u}\right)^T\right) \tag{71}$$

With,

$$\overline{\overline{\sigma}}_i^{el} = \begin{pmatrix} -\Delta P_n^i \\ 0 \\ -f^i \end{pmatrix} \cdot \overline{\overline{\mathbb{1}}} \tag{72}$$

Note that $\overline{\overline{\sigma}}_i^{el}$ was chosen to respect the mechanical force balance at the edge of the nucleus both in the (r,z) directions. $\Delta P_n^i = 2 \cdot \gamma \cdot \mathcal{C}$ is the Laplace pressure. Moreover, when there is no shear, $f^i = \Delta P_n^i$

For the sake of generality, we assumed in the previous equation that the envelope is tensed before the start of the confinement which induces a $\Delta P_n^i \neq 0$. This is tailored to describe the different small steps of confinements we apply on single HeLa nuclei ($6\mu m \longrightarrow 5\mu m$, $5\mu m \longrightarrow 4\mu m$, $4\mu m \longrightarrow 3\mu m$).

*b. Equation ruling nuclear volume change.* Using Eq.67 we derive the governing equation for the nuclear volume strain at the mesoscopic scale within the nuclear bulk. Using the definition of the flux velocity $\vec{q}$, we first rewrite the fluid velocity as:

$$\vec{q}^f = \frac{1}{1-\phi_0} \cdot \left(\vec{q} - \phi_0 \cdot \dot{\vec{u}}\right) \tag{73}$$

Thus Darcy law can be re-express as:

$$\vec{\nabla} P^f = \xi^f \cdot (\dot{\vec{u}} - \vec{q}) \tag{74}$$

The force balance now reads:

$$\vec{\nabla} \cdot \overline{\overline{\sigma}}^{el} = \xi^f \cdot (\dot{\vec{u}} - \vec{q}) \tag{75}$$

Using vector calculus, and the constitutive equation, we express $\vec{\nabla} \cdot \overline{\overline{\sigma}}^{el}$ as:

$$\vec{\nabla} \cdot \overline{\overline{\sigma}}^{el} = (K + \frac{4}{3}G) \cdot \Delta\vec{u} + (K + \frac{1}{3}G) \cdot \vec{\nabla} \wedge \vec{\nabla} \wedge \vec{u} \tag{76}$$

We thus have:

$$\vec{q} + D \cdot \Delta\vec{u} + \frac{(K + \frac{1}{3}G)}{\xi^f} \cdot \vec{\nabla} \wedge \vec{\nabla} \wedge \vec{u} = \partial_t \vec{u}\,, \quad D = \frac{K + \frac{4}{3}G}{\xi^f} \tag{77}$$

By further taking the divergence of the previous equation we obtain:

$$D \cdot \Delta\left(\vec{\nabla} \cdot \vec{u}\right) = \partial_t \left(\vec{\nabla} \cdot \vec{u}\right)\,, \quad D = \frac{K + \frac{4}{3}G}{\xi^f} \tag{78}$$

Where, $\vec{\nabla} \cdot \vec{u} = \frac{\delta V_n}{V_n^i}$ is the mesoscopic nuclear volume strain. While this diffusion equation is not convenient for a full resolution, it allows for a very simple interpretation of the water movements within the nucleus Fig.3C. Note that the modulus $K + \frac{4}{3}G$ is also known as the P-wave modulus, longitudinal modulus or constrained modulus in linear elasticity.

*c. Nuclear bulk displacement.* To make the problem analytically amenable, we will further assume that in the strong confinement regime under study here $h^2 << b^2$, the water movements in the z direction equilibrate quasi-statically. This is equivalent to assume that the time to equilibrate water movement in the z direction, $\tau_h \sim \frac{h^2}{D}$ is much faster than both the compression time $t^{\text{peak}}$ and the time to equilibrate the water movements in the radial direction $\tau_b \sim \frac{b^2}{D}$. Under this assumption we look for a non-rotational solution of the form $u_z(z,t)$, $u_r(r,t)$, $v_r(r,t)$, $v_z(z,t)$. The equation for the displacement thus reduces to:

$$v_r + D \cdot \Delta u_r = \partial_t u_r \quad \text{And,} \quad \Delta u_z = 0 \tag{79}$$

The displacement in the z-direction and the volume flux velocity are hence readily obtained as:

$$\Delta u_z = 0 \longrightarrow u_z = -\frac{qt}{h^i} \cdot z = -\frac{h^i - h(t)}{h^i} \cdot z \tag{80}$$

Moreover, using the incompressibility of the system {fluid + dry content} we obtain:

$$\vec{\nabla} \cdot \vec{q} = 0 \longrightarrow \{v_r = \frac{1}{2} \cdot \frac{q}{h^i} \cdot r, v_z = -\frac{q}{h^i} \cdot z\} \tag{81}$$

##### A. Computation of the linearized Laplace pressure $\Delta P_n$.

*a. Computation of the linearized nuclear surface.* We linearize the Laplace pressure according to the two relative displacements $\frac{u_r}{b^i}$ and $\frac{u_z}{h^i}$. By writting $h = h^i \cdot (1 + \frac{u_z}{h^i})$ and $b = b^i \cdot (1 + \frac{u_r}{b^i})$, the linearized total surface of the nucleus thus reads:

$$S = S^i + (4 \cdot S^{proj,i} + S^{e,i}) \cdot \frac{u_r}{b^i} + S^{e,i} \cdot \frac{u_z}{h^i} \tag{82}$$

*b. The linearized tension and curvature read.*

$$\gamma = \gamma^i + K_s \cdot \left(\frac{S^{e,i} + 4 \cdot S^{proj,i}}{S^*}\right) \cdot \frac{u_r}{b^i} + K_s \cdot \frac{S^{e,i}}{S^*} \cdot \frac{u_z}{h^i} \tag{83}$$

$$\mathcal{C} = \mathcal{C}^i - \frac{1}{2b^i} \cdot \frac{u_r}{b^i} - \frac{1}{h^i} \cdot \frac{u_z}{h^i} \tag{84}$$

*c. The linearized Laplace pressure reads.* We choose to normalize the pressures in this dynamic problem by the longitudinal modulus $K + \frac{4}{3}G$. The linearized expression of this Laplace pressure thus reads:

$$\overline{\Delta P_n} = \overline{\Delta P_n^i} + \tilde{K}_s^r \cdot \frac{u_r}{b^i} + \tilde{K}_s^z \cdot \frac{u_z}{h^i} \tag{85}$$

With,

$$\tilde{K}_s^r(K_s, S^*, b^i, h^i) = \frac{K_s}{K + \frac{4}{3}G} \cdot \left[\frac{1}{b^i}\left(1 + \frac{2S^{proj,i}}{S^*}\right) + \frac{2}{h^i}\left(\frac{S^{e,i} + 4S^{proj,i}}{S^*}\right)\right] \tag{86}$$

And,

$$\tilde{K}_s^z(K_s, S^*, b^i, h^i) = \frac{K_s}{K + \frac{4}{3}G} \cdot \left[\frac{1}{b^i} \cdot \frac{S^{e,i}}{S^*} - \frac{2}{h^i} \cdot \left(\frac{2S^{proj,i}}{S^*} - 1\right)\right] \tag{87}$$

Note that $K_s^z = -\Delta P_n^i + \mathcal{U}$, with $\mathcal{U} = \frac{2K_s}{b^i} \cdot \left(\frac{3S^{proj} + S^{e,i}}{S^*} - \frac{1}{2}\right)$. We thus check that when $b^i \longrightarrow \infty$ we find: $K_s^z = -\Delta P_n^i$. We also find that $K_s^r = \frac{K_s}{h^i} \cdot 8 \cdot \frac{S^{proj,i}}{S^*}$ ($S^{e,i}$ negligible in this limit too). Since in this limit $S \approx 2.S^{proj}$ we find the simpler expression: $K_s^r = 4 \cdot \frac{S^i}{S^*} \cdot \frac{K_s}{h^i}$

#### B. Boundary conditions

##### 1. *Permeation boundary condition.*

We compute the fluid pressure at the internal radial edge of the nucleus using the permeation condition:

$$P^f(b^i) - P^{f,out} = \frac{\xi_s \cdot b^i}{(1-\phi)} \cdot \left(\frac{1}{2} \cdot \frac{q}{h^i} - \frac{\dot{u}_r(b^i, t)}{b^i}\right) \tag{88}$$

We choose $P^{f,out}$ as the reference pressure and thus set it to 0 from now on. Note that we implicitly assume that the fluid pressure is identical everywhere out of the nucleus. This is justified in the limit under study where the dissipation that dominates is the friction between the fluid and the chromatin network and thus we neglected the term in $\eta^f \cdot \overline{\overline{\nabla}}v$.

##### 2. *Radial force balance.*

We now derive the boundary condition based on the constraints we discussed in the previous section. The radial force balance at the edge of the nucleus normalized by the longitudinal modulus reads:

$$\left.\overline{\overline{\overline{\sigma}}_{rr}} - \overline{P^f}\right|_{t, r=b^i} = -\overline{\Delta P_n(t)} \tag{89}$$

With,

$$\overline{\overline{\overline{\sigma}}_{rr}(b^i, \overline{t})} = -\overline{\Delta P_n^i} + \partial_r u_r + (1-\alpha) \cdot \frac{u_r}{b^i} - (1-\alpha) \cdot \frac{h^i - h(t)}{h^i} \quad , \quad \alpha = \frac{2G}{K + \frac{4}{3}G} \quad , \quad \tau = \frac{{b^i}^2}{D} \tag{90}$$

And,

$$\overline{P^f}(b^i) = \overline{\xi}_s \cdot \left(\frac{1}{2} \cdot \frac{q\tau}{h^i} - \frac{\tau \dot{u}_r(b^i, t)}{b^i}\right) \quad , \quad \overline{\xi}_s = \frac{\xi_s}{\xi_f \cdot (1-\phi) \cdot b^i} \tag{91}$$

This leads to the boundary condition:

$$\partial_r u_r + (1+\overline{K}_s)\cdot\frac{u_r}{b^i} + \bar{\xi}_s\cdot\tau\cdot\frac{\dot{u}_r}{b^i}\bigg|_{r=b^i,t} = (1+\overline{K}^z_s)\cdot\frac{qt}{h^i} + \frac{1}{2}\cdot\bar{\xi}_s\cdot\frac{q\tau}{h^i} \tag{92}$$

With, $\overline{K}_s = \tilde{K}^r_s - \alpha$ and $\overline{K}^z_s = \tilde{K}^z_s - \alpha$

*3. Summary of the problem*

The problem to solve in $u_r$ is the following:
*a. During confinement (* $\mathbf{t} \leq \mathbf{t}^{peak}$ *):*

$$\frac{1}{2}\cdot\frac{q}{h^i}\cdot r + D\cdot\Delta u_r = \partial_t u_r \tag{93}$$

With the boundary condition (BC) and initial condition (IC) reading:

$$\begin{aligned} &\text{IC}: \quad u_r(t=0,r)=0 \\ &\text{BC}: \quad \partial_r u_r + (1+\overline{K}_s)\cdot\tfrac{u_r}{b^i} + \bar{\xi}_s\cdot\tau\cdot\tfrac{\dot{u}_r}{b^i}\bigg|_{r=b^i,t} = (1+\overline{K}^z_s)\cdot\tfrac{h^i-h(t)}{h^i} + \tfrac{1}{2}\cdot\bar{\xi}_s\cdot\tfrac{q\tau}{h^i} \end{aligned} \tag{94}$$

*b. After confinement (* $\mathbf{t} \geq \mathbf{t}^{peak}$ *):* We take the previous problem with $q=0$. And we impose the continuity of the displacement at $t = t^{\text{peak}}$:

$$u_r(t=(t^{\text{peak}})^+) = u_r(t=(t^{\text{peak}})^-) \tag{95}$$

Note that this problem is non-hermitian as the eigen-vectors of the operator laplacian are not orthogonal. This is due to the presence of $\dot{u}_r$ in the BC. Nevertheless, the problem is still analytical as the eigen-vectors still form a basis.

#### C. Displacement resolution.

*1. Derivation of the solution during the uniaxial confinement at a constant velocity*

The mathematical trick is to look for a solution of the form:

$$u_r(r,t) = A(t)\cdot r + B\cdot r^3 + w(r,t) \tag{96}$$

It is sufficient for the problem under study to look for a solution where $A(t) = a_1\cdot t + a_2$ and $B$ is independent of time. Moreover, we impose that $A(t)\cdot r + B\cdot r^3$ respect the boundary condition of the problem. The problem to solve for w is thus:

$$D\Delta w - \partial_t w = (\dot{A} - \frac{1}{2}\cdot\frac{q}{h^i} - 8D\cdot B)\cdot r \tag{97}$$

$$\partial_r w + (1+\overline{K}_s)\cdot\frac{w}{b^i} + \bar{\xi}_s\cdot\tau\cdot\frac{\dot{w}}{b^i}\bigg|_{r=b^i} = 0 \tag{98}$$

$$w(r,0) = -A(0)\cdot r - B\cdot r^3 \tag{99}$$

We choose B in order to cancel out the right hand side of the diffusion equation:

$$B = \frac{1}{8D} \cdot \left( a_1 - \frac{1}{2} \cdot \frac{q}{h^i} \right) \tag{100}$$

$a_1$ and $a_2$ are determined with the boundary condition. They read:

$$a_1 = \frac{1}{2} \cdot \delta \cdot \frac{q}{h^i} \quad \text{and} \quad a_2 = \frac{1}{2} \cdot (1 - \delta) \cdot \frac{q}{h^i} \cdot \frac{1 + \frac{\overline{K}_s}{4}}{1 + \frac{\overline{K}_s}{2}} \cdot \frac{\tau}{4} \cdot \left( 1 + \frac{2 \cdot \bar{\xi}_s}{1 + \frac{\overline{K}_s}{4}} \right) \tag{101}$$

With,

$$\delta = \frac{1 + \overline{K_s}^z}{1 + \frac{\overline{K}_s}{2}} \tag{102}$$

We decompose w on the basis of eigen-vectors of the laplacian which respect the boundary condition:

$$w(y,t) = \sum_{n=1}^{\infty} w_n(t) \cdot J_1(x_n \cdot y) \tag{103}$$

Where, $J_1$ is the cylindrical Bessel function of the first kind. We do not consider the Bessel functions of the second kind because they would lead to divergence of the displacement at the center of the nucleus which is not physically acceptable. The time dependency of each mode is determined using the diffusion equation ruling the dynamics of w. Using the fact that $J_1$ is a eigen-vector of the laplacian vector ($\Delta J_1(x_n y) = -\left(\frac{x_n}{b^i}\right)^2 \cdot J_1(x_n y)$) we obtain:

$$w_n(t) = w_n(0) \cdot e^{-\frac{t}{\tau_n}} \quad \text{with,} \quad \tau_n = \frac{\tau}{x_n^2} \tag{104}$$

The modes $x_n$ are determined using the boundary condition for w:

$$x_n \cdot J_0(x_n) = (-\overline{K}_s + \bar{\xi}_s \cdot x_n^2) \cdot J_1(x_n) \tag{105}$$

We finally determine the constants $w_n(0)$ using the initial condition for w. To do so we decompose the linear function $r$ and the cubic function $r^3$ on the basis of the eigen-vectors:

$$r = \sum_{n=1}^{\infty} r_{n,1} \cdot b^i \cdot J_1(x_n y) \tag{106}$$

$$r^3 = \sum_{n=1}^{\infty} r_{n,3} \cdot b^{i^3} \cdot J_1(x_n y) \tag{107}$$

To determine the coefficients $r_{n,1}$ and $r_{n,3}$ (similar to fourier coefficients but not on the same basis of functions), we use the following integrals:

$$\begin{cases} \text{If } k \neq n: \int_0^1 J_1(x_k y) \cdot J_1(x_n y) \cdot y \, \mathrm{dy} = \frac{\mathrm{x_n J_0(x_n) J_1(x_k) - x_k J_0(x_k) J_1(x_n)}}{\mathrm{x_k^2 - x_n^2}} = -\bar{\xi}_\mathrm{s} \cdot \mathrm{J_1(x_n) J_1(x_k)} \\ \int_0^1 J_1(x_k y)^2 \cdot y \, \mathrm{dy} = \frac{1}{2} \cdot \left( \mathrm{J_0(x_k)^2 + J_1(x_k)^2 - \frac{2 \cdot J_0(x_k) \cdot J_1(x_k)}{x_k}} \right) = \frac{\mathrm{J_1(x_k)^2}}{\mathrm{2x_k^2}} \cdot \left( \mathcal{P}(\mathrm{x_k}) - 2 \cdot \bar{\xi}_\mathrm{s} \cdot \mathrm{x_k^2} \right) \\ \int_0^1 J_1(x_k y) \cdot y^2 \, \mathrm{dy} = \frac{\mathrm{J_2(x_k)}}{\mathrm{x_k}} = \frac{\mathrm{J_1(x_k)}}{\mathrm{x_k^2}} \cdot (2 + \overline{\mathrm{K}}_\mathrm{s} - \bar{\xi}_\mathrm{s} \cdot \mathrm{x_k^2}) \\ \int_0^1 J_1(x_k y) \cdot y^4 \, \mathrm{dy} = \frac{\mathrm{4 \cdot J_3(x_k) - x_k \cdot J_4(x_k)}}{\mathrm{x_k^2}} = \frac{\mathrm{J_1(x_k)}}{\mathrm{x_k^4}} \cdot \left( \mathcal{Q}(\mathrm{x_k}) - \bar{\xi}_\mathrm{s} \cdot \mathrm{x_k^4} \right) \end{cases} \tag{108}$$

With,

$$\mathcal{P}(x_n) = \overline{\xi}_s^2 \cdot x_n^4 + x_n^2 \cdot \left(1 - 2 \cdot \overline{K}_s \cdot \bar{\xi}_s\right) + \overline{K}_s \cdot (\overline{K}_s + 2) \tag{109}$$

And,

$$\mathcal{Q}(x_k) = x_k^2 \cdot \left(\overline{K}_s + 4 + 8 \cdot \bar{\xi}_s\right) - 8 \cdot (2 + \overline{K}_s) \tag{110}$$

$r_{n,1}$ and $r_{n,3}$ can now be expressed as:

$$r_{n,1} = \frac{4 \cdot \left(1 + \frac{\overline{K}_s}{2}\right)}{\mathcal{P}(x_n) \cdot J_1(x_n)} \tag{111}$$

$$r_{n,3} = \frac{2 \cdot \mathcal{Q}(x_n)}{\mathcal{P}(x_n) \cdot x_n^2 \cdot J_1(x_n)} \tag{112}$$

Where we have made use of the equality $1 = \sum_{n=1}^{\infty} r_{n,1} \cdot J_1(x_n)$. The solution $u_r$ thus reads:

$$u_r(r,t) = a_1 \cdot t \cdot r + \sum_{n=1}^{\infty} \left(a_2 \cdot b^i \cdot r_{n,1} + B \cdot b^{i^3} \cdot r_{n,3}\right) \cdot \left(1 - e^{-\frac{t}{\tau_n}}\right) \cdot J_1(x_n y) \tag{113}$$

Which simplifies into:

$$\frac{u_r(y,t)}{b^i} = \frac{1}{2} \cdot \delta \cdot \frac{q \cdot t}{h^i} \cdot y + (1-\delta) \cdot \frac{1}{2} \cdot \frac{q}{h^i} \cdot \sum_{n=1}^{\infty} \tau_n \cdot r_{n,1} \cdot (1 - e^{-\frac{t}{\tau_n}}) \cdot J_1(x_n y) \tag{114}$$

##### *2. Solution in the post-confinement relaxation phase.*

The solution in the relaxation phase is obtained using the same method, imposing continuity of displacement for $t = t^{\text{peak}}$. The boundary condition reads:

$$\partial_r u_r + (1 + \overline{K}_s) \cdot \frac{u_r}{b^i} + \bar{\xi}_s \cdot \tau \cdot \frac{\dot{u}_r}{b^i}\bigg|_{r=b^i,t} = (1 + \overline{K}_s^z) \cdot \frac{h^i - h^\infty}{h^i} \tag{115}$$

We look for an ansatz solution of the form:

$$u_r(r,t) = a_1^\infty \cdot r + w(r,t) \tag{116}$$

$a_1^\infty$ is determined using the boundary condition and reads:

$$a_1^\infty = \frac{1}{2} \cdot \delta \cdot \frac{h^i - h^\infty}{h^i} \tag{117}$$

$w$ is decomposed on the same basis. Solving the diffusion equation for $t \geq t^{\text{peak}}$:

$$\omega_n(t) = \omega_n(t^{\text{peak}}) \cdot e^{-\frac{t - t^{\text{peak}}}{\tau_n}} \tag{118}$$

The constant $\omega_n(t^{\text{peak}})$ is determined using the continuity of the displacement at $t = t^{\text{peak}}$. We finally obtain:

$$\frac{u_r(y,t)}{b^i} = \frac{1}{2} \cdot \delta \cdot \frac{h^i - h^\infty}{h^i} \cdot y + (1-\delta) \cdot \frac{1}{2} \cdot \frac{q}{h^i} \cdot \sum_{n=1}^{\infty} \tau_n \cdot r_{n,1} \cdot (1 - e^{-\frac{t^{\text{peak}}}{\tau_n}}) \cdot e^{-\frac{t - t^{\text{peak}}}{\tau_n}} \cdot J_1(x_n y) \tag{119}$$

##### D. Derivation of the force during and after confinement.

###### 1. Force per unit surface.

We use the mechanical boundary condition expressed in Eq.68 for $z = h^i$ to express the force exerted on the top surface of the gel.

$$f(y,t) = -\overline{\overline{\sigma}}_{zz}(h^i, y, t) + P^f(y,t) \tag{120}$$

*a. Vertical stress.* We next express the vertical stress $\overline{\overline{\sigma}}_{zz}(h^i, y, t)$ using the elastic constitutive equation.

$$\overline{\overline{\sigma}}_{zz}(h^i, y, t) = -f^i + (K - \frac{2}{3}G) \cdot \vec{\nabla} \cdot \vec{u} + 2G \cdot \partial_z u_z \tag{121}$$

Which can be re-expressed in normalized form as:

$$-\overline{\overline{\overline{\sigma}}_{zz}} = \overline{f^i} - (1-\alpha) \cdot \left(\partial_r u_r + \frac{u_r}{r}\right) - \partial_z u_z \tag{122}$$

In both regimes, $u_r$ and $u_z$ are of the form:

$$u_r = \gamma_1 \cdot r + \sum \gamma_{2,n} \cdot b^i \cdot J_1(x_n y) \tag{123}$$

$$u_z = -\frac{h^i - h(t)}{h^i} \cdot z \tag{124}$$

With,

$$\gamma_1 = \frac{1}{2} \cdot \delta \cdot \left(1 - \frac{h(t)}{h^i}\right) \tag{125}$$

And,

$$\gamma_{2,n} = \frac{1}{2} \cdot (1-\delta) \cdot \frac{q}{h^i} \cdot \tau_n \cdot r_{n,1} \cdot v_n(t) \tag{126}$$

Thus, the radial part of the divergence reads:

$$\partial_r u_r + \frac{u_r}{r} = 2\gamma_1 + \gamma_{2,n} x_n J_0(x_n y) \tag{127}$$

Where we made use of the fact that $J_1'(x) + \frac{J_1}{x} = J_0(x)$ and implicitly assumed summation over the modes. The resulting stress reads:

$$-\overline{\overline{\overline{\sigma}}_{zz}} = \overline{f^i} + \frac{h^i - h(t)}{h^i} - (1-\alpha) \cdot (2\gamma_1 + \gamma_{2,n} x_n J_0(x_n y)) \tag{128}$$

*b. Fluid pressure in the nucleus.* We first express the fluid pressure in the nucleus using Darcy's law and the permeation boundary condition. We find:

$$P^f(y,t) = \frac{\xi_s}{(1-\phi)} \cdot (v_r(b^i,t) - \dot{u}_r(b^i,t)) + \xi_f \cdot b^i \cdot \int_y^1 (v_r(y,t) - \dot{u}_r(y,t))\, dy \tag{129}$$

With, $y = \frac{r}{b^i}$. The difference of velocity reads:

$$\frac{v_r - \dot{u}_r}{b^i} = \left( (\frac{1}{2} \cdot \frac{q}{h^i} \cdot \zeta - \dot{\gamma}_1) \cdot r_{n,1} - \dot{\gamma}_{2,n} \right) \cdot J_1(x_n y) \tag{130}$$

Where, we made use of the fact that $1 = \sum_{n=1}^{\infty} r_{n,1} \cdot J_1(x_n)$ and $\zeta$ is a dirac-like function equal to 1 during confinement and 0 after. Using the relation $\int_y^1 J_1(x_n y)\mathrm{d}y = \frac{-J_0(x_n)+J_0(x_n y)}{x_n}$ we can further express the normalized expression of the fluid pressure as:

$$\overline{P^f} = \tau \cdot \left( \bar{\xi}_s \cdot \frac{v_r(b^i,t) - \dot{u}_r(b^i,t)}{b^i} + \int_y^1 \frac{v_r - \dot{u}_r}{b^i} \mathrm{d}y \right) \tag{131}$$

Finally, using the previous two equations and the relation $x_n \cdot J_0(x_n) = (-\overline{K}_s + \bar{\xi}_s \cdot x_n^2) \cdot J_1(x_n)$, we express the fluid the pressure as:

$$\overline{P^f} = \tau_n \cdot J_1(x_n) \cdot \left( \frac{1}{2} \cdot (1-\delta) \cdot \frac{q}{h^i} \cdot \zeta \cdot r_{n,1} - \dot{\gamma}_{2,n} \right) \cdot \left( \overline{K}_s + x_n \cdot \frac{J_0(x_n y)}{J_1(x_n)} \right) \tag{132}$$

Replacing the expression of $\gamma_1$ and $\gamma_{2,n}$ we can finally derive the normalized force per unit surface.

*c. Force per unit surface during confinement (* $\mathbf{t} \leq \mathbf{t}^{peak}$ *):*

$$\overline{f}(y,t) = \overline{f}^i + \frac{q}{h^i} \cdot \Bigg\{ (1-(1-\alpha)\cdot\delta)\cdot t\,(...)$$
$$(...) + (2+\overline{K}_s)\cdot(1-\delta)\cdot\sum_{n=1}^{\infty}\left(\overline{K}_s + \alpha\cdot\frac{x_n \cdot J_0(x_n y)}{J_1(x_n)}\right)\cdot\frac{\tau_n}{\mathcal{P}(x_n)}\cdot(1-e^{-\frac{t}{\tau_n}})\Bigg\} \tag{133}$$

*d. Force per unit surface post-confinement (* $\mathbf{t} \geq \mathbf{t}^{peak}$ *):*

$$\overline{f}(y,t) = \overline{f}^i + \frac{q}{h^i} \cdot \Bigg\{ (1-(1-\alpha)\cdot\delta)\cdot t^{\text{peak}}\,(...)$$
$$(...) + (2+\overline{K}_s)\cdot(1-\delta)\cdot\sum_{n=1}^{\infty}\left(\overline{K}_s + \alpha\cdot\frac{x_n \cdot J_0(x_n y)}{J_1(x_n)}\right)\cdot\frac{\tau_n}{\mathcal{P}(x_n)}\cdot(1-e^{-\frac{t^{\text{peak}}}{\tau_n}})\cdot e^{-\frac{t-t^{\text{peak}}}{\tau_n}}\Bigg\} \tag{134}$$

*2. Force to deform nuclei at constant velocity.*

The force $F$ applied on the nucleus by the cantilever reads:

$$F = \int_0^{\frac{b(t)}{b^i}} f(y,t) \cdot {b^i}^2 \cdot 2\pi y dy \tag{135}$$

With :

$$\frac{b(t)}{b^i} = 1 + \frac{u_r}{b^i} \tag{136}$$

The normalized pressure exerted by the cantilever on the nucleus is of the form (with implicit einstein summation on n):

$$\overline{f} = \overline{f}^i + \gamma_3 + \gamma_{4,n} \cdot \left( \overline{K}_s + \alpha \cdot x_n \cdot \frac{J_0(x_n y)}{J_1(x_n)} \right) \tag{137}$$

With:

$$\gamma_3 = \frac{q}{h^i} \cdot \left(1 - \frac{h(t)}{h^i}\right) \cdot (1 - (1-\alpha)\cdot\delta) \quad , \quad \gamma_{4,n} = 2 \cdot \frac{q}{h^i} \cdot (1-\delta) \cdot \left(1 + \frac{\overline{K}_s}{2}\right) \cdot \frac{\tau_n}{\mathcal{P}(x_n)} \cdot v(t) \tag{138}$$

Note that we can reexpress $(1-\delta)\cdot\left(1+\frac{\overline{K}_s}{2}\right)$ as $\left(\frac{\overline{K}_s}{2} - \overline{K}_s^z\right)$. We also emphasize that the upper born of the integral is not one. This is due to the fact that in f(y,t) there is an order 0 (the initial force per unit surface contained in $\overline{\overline{\sigma}}_{zz}$) such that we would miss an order one term if we limited the integration to the order 0. Truncating all terms to order one displacement we find :

$$\frac{F}{2\pi b^{i^2} \cdot (K + \frac{4}{3}G)} = \frac{1}{2} \cdot \overline{f}^i + \frac{1}{2} \cdot (1 - \frac{h}{h^i}) \cdot \left(1 + \delta \cdot (\overline{f}^i + \alpha - 1)\right) + \gamma_{4,n} \cdot \left(\overline{f}^i + \alpha + \frac{\overline{K}_s}{2}\right) \tag{139}$$

Using that, $F^i = \pi \cdot b^{i^2} \cdot f^i$ and $\gamma_{4,n} \longrightarrow 0$ when $t \longrightarrow \infty$, we first express the post-relaxational force as:

$$\frac{F^\infty - F^i}{F^i} = \frac{1}{\overline{f}^i} \cdot \left(1 - \frac{h^\infty}{h^i}\right) \cdot \left(1 + \delta \cdot (\overline{f}^i + \alpha - 1)\right) \tag{140}$$

Where we used that :

$$\frac{u_r(b^i, t)}{b^i} = \frac{1}{2} \cdot \delta \cdot (1 - \frac{h}{h^i}) + \gamma_{4,n} \tag{141}$$

And,

$$\int_0^1 J_0(x_n y) y \mathrm{d}y = \frac{J_1(x_n)}{x_n} \tag{142}$$

Finally, the force reads:

$$\frac{F - F^i}{F^i} = \frac{F^\infty - F^i}{F^i} \cdot \frac{t}{t^{\text{peak}}} + \frac{1}{\overline{f}^i} \cdot 4 \cdot \frac{q}{h^i} \cdot \left(\frac{\overline{K}_s}{2} - \overline{K}_s^z\right) \cdot \left(\overline{f}^i + \alpha + \frac{\overline{K}_s}{2}\right) \cdot \sum_{n=1}^{\infty} \frac{\tau_n}{\mathcal{P}(x_n)} \cdot v_n(t) \tag{143}$$

##### E. Summary

The solutions of the problem read :

$$\frac{u_r(y,t)}{b^i} = \frac{1}{2} \cdot \delta \cdot \left(1 - \frac{h(t)}{h^i}\right) \cdot y + 2 \cdot \frac{q}{h^i} \cdot \left(\frac{\overline{K}_s}{2} - \overline{K}_s^z\right) \cdot \sum_{n=1}^{\infty} \frac{\tau_n}{\mathcal{P}(x_n)} \cdot v_n(t) \cdot \frac{J_1(x_n y)}{J_1(x_n)} \tag{144}$$

$$\frac{u_z(z,t)}{h^i} = -\frac{q \cdot t^{\text{peak}}}{h^i} \cdot \zeta(t) \cdot z \tag{145}$$

$$\overline{f}(y,t) = \overline{f}^i + (1 - (1-\alpha)\cdot\delta) \cdot \left(1 - \frac{h(t)}{h^i}\right) + 2 \cdot \frac{q}{h^i} \cdot \left(\frac{\overline{K}_s}{2} - \overline{K}_s^z\right) \cdot \sum_{n=1}^{\infty} \left(\overline{K}_s + \alpha \cdot \frac{x_n \cdot J_0(x_n y)}{J_1(x_n)}\right) \cdot \frac{\tau_n}{\mathcal{P}(x_n)} \cdot v_n(t) \Bigg\} \tag{146}$$

$$\mathcal{F} = \mathcal{F}^{\infty} \cdot \zeta(t) + \frac{1}{\overline{f}^i} \cdot 4 \cdot \frac{q}{h^i} \cdot \left(\frac{\overline{K}_s}{2} - \overline{K}_s^z\right) \cdot \left(\overline{f}^i + \alpha + \frac{\overline{K}_s}{2}\right) \cdot \sum_{n=1}^{\infty} \frac{\tau_n}{\mathcal{P}(x_n)} \cdot v_n(t) \tag{147}$$

With,

$$\mathcal{F}(t) = \frac{F(t) - F^i}{F^i} \quad , \quad F^i = f^i \cdot \pi \cdot {b^i}^2 \quad , \quad \mathcal{F}^{\infty} = \frac{1}{\overline{f}^i} \cdot \left(1 - \frac{h^{\infty}}{h^i}\right) \cdot \left(1 + \delta \cdot (\overline{f}^i + \alpha - 1)\right) \quad , \quad \overline{f}^i = \frac{f^i}{K + \frac{4}{3}G} \tag{148}$$

$$v_n(t) = \begin{cases} \text{for } t \geq t^{\text{peak}}, & (1 - e^{-\frac{t^{\text{peak}}}{\tau_n}}) \cdot e^{-\frac{t - t^{\text{peak}}}{\tau_n}} \\ \text{for } t \leq t^{\text{peak}}, & (1 - e^{-\frac{t}{\tau_n}}) \end{cases} \tag{149}$$

$$h(t) = h^i - q \cdot t^{\text{peak}} \cdot \zeta(t) \quad , \quad \zeta(t) = \begin{cases} \text{for } t \leq t^{\text{peak}}, & \frac{t}{t^{\text{peak}}} \\ \text{for } t \geq t^{\text{peak}}, & 1 \end{cases} \tag{150}$$

$$\overline{K}_s = \frac{K_s}{K + \frac{4}{3}G} \cdot \left[\frac{1}{b^i}\left(1 + \frac{2S^{proj,i}}{S^*}\right) + \frac{2}{h^i}\left(\frac{S^{e,i} + 4S^{proj,i}}{S^*}\right)\right] - \alpha \tag{151}$$

And,

$$\overline{K}_s^z = \frac{K_s}{K + \frac{4}{3}G} \cdot \left[\frac{1}{b^i} \cdot \frac{S^{e,i}}{S^*} - \frac{2}{h^i} \cdot \left(\frac{2S^{proj,i}}{S^*} - 1\right)\right] - \alpha \tag{152}$$

$$\alpha = \frac{2G}{K + \frac{4}{3}G} \quad , \quad \delta = \frac{1 + \overline{K}_s^z}{1 + \frac{\overline{K}_s}{2}} \quad , \quad \tau_n = \frac{{b^i}^2}{D \cdot x_n^2} \quad , \quad D = \frac{K + \frac{4}{3}G}{\xi_f} \tag{153}$$

$$x_n \cdot J_0(x_n) = (-\bar{K}_s + \bar{\xi}_s \cdot x_n^2) \cdot J_1(x_n) \tag{154}$$

$$\mathcal{P}(x_n) = \bar{\xi}_s^2 \cdot x_n^4 + x_n^2 \cdot \left(1 - 2 \cdot \bar{K}_s \cdot \bar{\xi}_s\right) + \bar{K}_s \cdot (\bar{K}_s + 2) \tag{155}$$

$$\bar{\xi}_s = \frac{\xi_s}{(1 - \phi) \cdot \xi_f \cdot b^i} \tag{156}$$

#### F. Verification of the solution.

##### 1. Limits of constant volume.

*a. Volume.* In all of the limits below:

$$\begin{cases} K_s \longrightarrow 0 \quad , \quad \alpha = 0 \\ K \longrightarrow \infty \\ \xi_s \longrightarrow \infty \\ \xi_f \longrightarrow \infty \end{cases} \tag{157}$$

We verify that the general solution gives:

$$u_r = \frac{1}{2} \cdot \frac{qt}{h^i} \cdot r \tag{158}$$

Which corresponds to a deformation at constant volume in the linear regime.

*2. Regime where the surface dominates the dynamics.*

We consider the limit $\xi_f \longrightarrow 0$.

*a. Simplification of the general solution.* Starting from the mode equation we have:

$$x_n^2 \cdot J_1(x_n) = \frac{x_n \cdot J_0(x_n) + \bar{K}_s \cdot J_1(x_n)}{\bar{\xi}_s} \tag{159}$$

Thus, for $n > 1$, we have $J_1(x_n) = 0$ and $x_n$ have non-zero values. However, for $n = 1$, we have $x_1 \longrightarrow 0$. This implies that:

$$J_1(x_1) = \frac{x_1}{2}, \quad J_0(x_1) = 1 - \frac{x_1^2}{4} \tag{160}$$

This leads to:

$$x_1 = \sqrt{\frac{2 + \bar{K}_s}{\bar{\xi}_s}} \tag{161}$$

$$\mathcal{P}(x_1) = 4 \cdot \left(1 + \frac{\bar{K}_s}{2}\right), \quad r_{1,1} = \frac{1}{J_1(x_1)} = \frac{2}{x_1} \tag{162}$$

$$\begin{cases} \tau_1 = \frac{\xi_s \cdot b^i}{K \cdot (1-\phi)(2+\bar{K}_s)} \\ \text{For } n > 1, \tau_n \longrightarrow 0 \end{cases} \tag{163}$$

All the modes relax infinitely fast except the first one. The relaxation thus becomes single exponential due to the envelope friction:

$$\frac{u_r(y,t)}{b^i} = \frac{1}{2} \cdot \delta \cdot \frac{q \cdot t}{h^i} \cdot y + (1-\delta) \cdot \frac{1}{2} \cdot \frac{q}{h^i} \cdot \tau_1 \cdot r_{1,1} \cdot (1 - e^{-\frac{t}{\tau_1}}) \cdot J_1(x_1 y) \tag{164}$$

Which can be written as given that $J_1(x_1 y) \approx \frac{x_1 y}{2}$:

$$\frac{u_r(y,t)}{b^i} = \frac{1}{2} \cdot \delta \cdot \frac{q \cdot t}{h^i} \cdot y + (1-\delta) \cdot \frac{1}{2} \cdot \frac{q}{h^i} \cdot \tau_1 \cdot (1 - e^{-\frac{t}{\tau_1}}) \cdot y \tag{165}$$

*b. Direct resolution of the Dynamical equation.* The diffusion coefficient is infinite in this limit. This implies that:

$$\Delta u_r = \frac{1}{D} \cdot (\partial_t u_r - v_r) = 0 \tag{166}$$

Thus,

$$u_r = A(t) \cdot r \tag{167}$$

Using the boundary condition we obtain the differential equation that rules the evolution of $A(t)$:

$$\dot{A}(t) + \frac{A}{\tau_1} = \frac{1}{2} \cdot \frac{q}{h^i} \cdot (1 + \delta \cdot \frac{t}{\tau_1}) \tag{168}$$

Knowing that A(0) = 0, this implies that:

$$A(t) = \frac{1}{2} \cdot \delta \cdot \frac{q \cdot t}{h^i} + (1-\delta) \cdot \frac{1}{2} \cdot \frac{q}{h^i} \cdot \tau_1 \cdot (1 - e^{-\frac{t}{\tau_1}}) \tag{169}$$

Which is coherent with the general solution.

*c. Agreement with previous works in this limit.* In the classical Pump-leak model, the equation ruling the volume change dynamic is written as:

$$\frac{dV}{dt} = l_p \cdot (\Delta\Pi - \Delta P) \tag{170}$$

Where, $l_p$ is the permeability of the membrane. While our formalism is different to include the poroelasticity of the bulk, we emphasize that this equation is equivalent to our boundary condition Eq.88 in the limit under study where only one timescale related to the surface dominates. Indeed, the fluid pressure in our theory is the difference between the hydrostatic pressure and the osmotic pressure.

##### G. Comparison with experiments.

Based on the experimental design and data, we justify in this section the fitting pipeline that we designed to extract the physical model's parameters from the force relaxation curves.

###### 1. *Discussion on the relation between $\alpha$, $f^i$ and $\Delta P_n^i$.*

For the sake of generality, we purposely introduced two constants in our calculation - $f^i$ and $\Delta P_n^i$ - to respectively describe the radial and vertical stress before the step compression. Indeed, when there is a shear modulus, the radial and vertical stresses become different. This suggests that $f^i$ and $\Delta P_n^i$ are not independent but are related through $\alpha$. To show the dependency, we consider two resting states before and after compression, denominated by the uppercase $^i$ and $^\infty$. We have:

$$\overline{\sigma_{zz}}^\infty = -\overline{f}^i + (1-\alpha)\cdot(\partial_r u_r + \frac{u_r}{r}) + \partial_z u_z \tag{171}$$

$$\overline{\sigma_{rr}}^\infty = -\overline{\Delta P_n}^i + \partial_r u_r + (1-\alpha)\cdot\frac{u_r}{r} + (1-\alpha)\cdot\partial_z u_z \tag{172}$$

At steady-state, the laplacian of the displacement vanishes such that the radial and vertical displacements are linear functions of their coordinates : $u_r = \frac{1}{2}\delta(1-\tilde{h})r$ and $u_z = -(1-\tilde{h})z$, with $\tilde{h} = \frac{h^\infty}{h^i}$. The stress is thus homegenenous in the nucleus. Moreover, using the boundary conditions we have that $\overline{\sigma_{rr}}^\infty = -\overline{\Delta P_n}^\infty$ and $\overline{\sigma_{zz}}^\infty = -\overline{f}^\infty$. We thus have:

$$\overline{f}^\infty = \overline{f}^i - (1-\alpha)\cdot\delta\cdot(1-\tilde{h}) + (1-\tilde{h}) \tag{173}$$

$$\overline{\Delta P_n}^\infty = \overline{\Delta P_n}^i - (1-\frac{\alpha}{2})\cdot\delta\cdot(1-\tilde{h}) + (1-\alpha)\cdot(1-\tilde{h}) \tag{174}$$

We can thus express $\delta\cdot(1-\tilde{h})$ as :

$$\delta\cdot(1-\tilde{h}) = \frac{(1-\tilde{h}) - (\overline{f}^\infty - \overline{f}^i)}{(1-\alpha)} \tag{175}$$

Such that we obtain the relation :

$$\overline{\Delta P_n}^\infty - \overline{\Delta P_n}^i = \left(\frac{1-\frac{\alpha}{2}}{1-\alpha}\right)\cdot\left(\overline{f}^\infty - \overline{f}^i\right) - \frac{\alpha\cdot(\frac{3}{2}-\alpha)}{1-\alpha}\cdot(1-\tilde{h}) \tag{176}$$

Let's consider the step that passes through the force threshold such that $\Delta P_n^i = f^i = 0$ but $f^\infty \neq 0$. This step typically corresponds to the $6->5\mu m$ confinement in our experiments. This implies that we have:

$$\overline{\Delta P_n}^{5\mu m} = \left(\frac{1-\frac{\alpha}{2}}{1-\alpha}\right) \cdot \left(\overline{f}^{5\mu m}\right) - \frac{\alpha \cdot (\frac{3}{2}-\alpha)}{1-\alpha} \cdot (1 - \frac{h^{5\mu m}}{h^{6\mu m}}) \tag{177}$$

And we could do the same for the next confinement step $5- > 4\mu m$:

$$\overline{\Delta P_n}^{4\mu m} - \overline{\Delta P_n}^{5\mu m} = \left(\frac{1-\frac{\alpha}{2}}{1-\alpha}\right) \cdot \left(\overline{f}^{4\mu m} - \overline{f}^{5\mu m}\right) - \frac{\alpha \cdot (\frac{3}{2}-\alpha)}{1-\alpha} \cdot (1 - \frac{h^{4\mu m}}{h^{5\mu m}}) \tag{178}$$

We chose to simplify the relationship between $\Delta P_n^i$ and $f^i$ in our fitting pipeline to $\Delta P_n^i \approx f^i$. This approximation is valid because our data falls within a regime where the parameter $\alpha$ is close to zero.

###### *2. Discussion on the parameter $\alpha$*

Our data supports the approximation that $\alpha \approx 0$ for two reasons:

- First, we do not observe any significant post-relaxational force increase for the confinement steps $12 \longrightarrow 8\mu m$ and $8 \longrightarrow 6\mu m$ while the nuclei are still significantly compressed. Indeed, the typical diameter of HeLa cell nuclei is $12\mu m$ considering an average volume of $870\mu m^3$. We extend our force formulas for the $8 \longrightarrow 6\mu m$ confinement step where the envelope is not stretched, which makes the effective stretching modulus of the envelope going to 0 in our formulas $K_s \longrightarrow 0$ and $F^i = 0$. We have:

$$F^{\infty} = 2G \cdot S^{proj,i} \cdot (1 - \tilde{h}^{\infty}) \cdot \frac{(\frac{3}{2}-\alpha)}{1-\frac{\alpha}{2}} \tag{179}$$

  Given that we observe $F^{\infty} \approx 0$ before the envelope's stretching, we infer that the contribution of the shear modulus ($G$) is below the detection limit of our force measurements. This allows us to make the approximation $G \approx 0$, and hence $\alpha \approx 0$.

- The approximation is also justified a posteriori: fitting the model yields $\overline{K}_s > 1$, whereas a dominant $\alpha$ would have resulted in $\overline{K}_s < 0$.

###### *3. Discussion on the volume loss during the compression.*

In this subsection we show that in the limit of the experiment where $t^{\text{peak}} << \tau_1$, the volume loss during the compression, .i.e. $0 \leq t \leq t^{\text{peak}}$, is negligible and we can thus simplify the equations.

###### *4. Force profiles during a compression at constant volume*

In our equations, the exact condition of compression at no volume loss is:

$$\tau_1 > \tau_2 > ... > \tau_n >> t^{\text{peak}} \tag{180}$$

Under this assumption we can approximate :

$$e^{-\frac{t}{\tau_n}} \sim 1 - \frac{t}{\tau_n} \tag{181}$$

Thus,

$$\frac{u_r(b^i,t)}{b^i} = \frac{1}{2} \cdot \delta \cdot \frac{qt}{h^i} + 2 \cdot \frac{qt}{h^i} \cdot \left(\frac{\overline{K}_s}{2} - \overline{K}_s^z\right) \cdot \sum_{n=1}^{\infty} \frac{1}{\mathcal{P}(x_n)} \tag{182}$$

Given that, $\sum_{n=1}^{\infty} \frac{1}{\mathcal{P}(x_n)} = \frac{1}{4\cdot(1+\frac{\overline{K}_s}{2})}$, we finally obtain that :

$$\frac{u_r(b^i,t)}{b^i} = \frac{1}{2}\cdot\frac{qt}{h^i} \tag{183}$$

We also have,

$$V = \pi b^2 \cdot h \tag{184}$$

With, $b = b^i + u_r(b^i)$ and $h = h^i + u_z(h^i)$. At linear order V becomes :

$$\frac{V(t)-V^i}{V^i} = \frac{u_z(h^i,t)}{h^i} + 2\cdot\frac{u_r(b^i,t)}{b^i} \tag{185}$$

Thus, the condition Eq.183 implies that the volume is conserved during the compression. Moreover, in this limit we also have that the force increases lineary with time:

$$\frac{F-F^i}{F^{t^{\text{peak}}}-F^i} = \frac{t}{t^{\text{peak}}} \tag{186}$$

Moreover, in this limiting regime the post-confinement relaxation of the force reads :

$$F(t) = F^{\infty} + (F^{t^{\text{peak}}} - F^{\infty})\cdot 4\cdot(1+\frac{\overline{K}_s}{2})\cdot\sum_{n=1}^{\infty}\frac{1}{\mathcal{P}(x_n)}\cdot e^{-\frac{t-t^{\text{peak}}}{\tau_n}} \tag{187}$$

*5. Experimental force and volume are well approximated by the limiting regime of no volume loss during compression.*

We nevertheless note that as $n\to\infty$, $\tau_n\to 0$. Thus, the criterion in Eq.180 fails beyond a certain mode for any fixed compression time (Supplementary Fig. 5B). For the experimental parameters, the criterion ceases to hold for modes $n \gtrsim 10$. This implies that, irrespective of the compression time, a residual volume loss is unavoidable during the compression. The question is therefore whether this residual volume loss is significant under our experimental conditions. To theoretically estimate the volume loss during the 2 s compression in our experiments, we computed the relative errors on the volume and force during the compression between the analytical solution without approximation (see Sec. II E) and the limiting regime of no volume loss. We define such errors as:

$$\mathcal{E}^V_{\%} = \left|\frac{V(t)-V^i}{V(t)}\right|\cdot 100 \quad , \quad \mathcal{E}^F_{\%} = \left|\frac{F(t)-F^{V=\text{cst}}(t)}{F(t)}\right|\cdot 100 \tag{188}$$

Note that for convenience regarding Eq.147 we re-wrote $\mathcal{E}^F_{\%}$ as :

$$\mathcal{E}^F_{\%} = \left|1 - \frac{1+\frac{\delta F^{V=\text{cst}}}{F^i}}{1+\frac{\delta F}{F^i}}\right|\cdot 100 \tag{189}$$

The parameters used to compute these errors were determined self-consistently from the inference pipeline under the assumption of no volume loss (Fig. 6). We summarize them in Table.III. We find that the maximal relative error in volume during compression is approximately 1% (Fig. 5E, inset, blue curve), while the relative error in force is less than 1% (Supplementary Fig. 5A). These results confirm that our equations can be accurately approximated by the limiting regime of no-volume loss during compression. Moreover, we experimentally verified that the force increase during compression was linear as predicted by the limiting regime (Eq. 186) (Supplementary Fig. 5C-E). Note that, in some replicates (Supplementary Fig. 5D), the force increase is not perfectly linear in time. We attribute this deviation not to volume loss, but rather to small departures from constant confinement speed due to experimental control errors (Fig. 5D, yellow curve).

TABLE III. Description and values of the parameters used to estimate the error on the force and volume assuming no volume loss during the 5 to 4 $\mu$m confinement step (Fig. 5D-F and Supplementary Fig. 5A).

| **Symbol** | **Typical Value** | **Meaning** |
|---|---|---|
| $h_i$ | $5\,\mu$m | Initial confinement height. |
| $h$ | $4\,\mu$m | Height after confinement. |
| $K_s$ | $6\,$mN/m | NE stretching modulus. |
| $F_i$ | $6\,$nN | Force before compression. |
| $S^*$ | $600\,\mu$m$^2$ | Nuclear surface at volume loss threshold. |
| $S^{proj,i}$ | $170\,\mu$m$^2$ | Midplane nuclear surface area before compression. |
| $K$ | $500\,$Pa | Nuclear bulk modulus. |
| $t^{\text{peak}}$ | $2\,$s | Duration of the compression step. |
| $v$ | $0.5\,\mu$m/s | Compression speed during the confinement step. |
| $G$ | $0\,$Pa | Nuclear bulk shear modulus (see Section II G 2 for a discussion). |
| $\xi_s$ | $0\,\text{Pa}\cdot\text{s}\cdot\text{m}^{-1}$ | Surface friction (see Section II G 8 for a discussion). |
| $\tau_1$ | $60\,$s | Typical timescale of nuclear volume loss as observed in experiments. |

*6. Relationship between force and volume relaxations.*

In the experimental regime where the volume loss is negligible during the compression, we show in this section that both the normalized force and volume relaxations are identical. In the latter limit we have :

$$v_n(t) = \begin{cases} \text{for } t \geq t^{\text{peak}}, & \sim \frac{t^{\text{peak}}}{\tau_n} \cdot e^{-\frac{t-t^{\text{peak}}}{\tau_n}} \\ \text{for } t \leq t^{\text{peak}}, & (1 - e^{-\frac{t}{\tau_n}}) \end{cases} \tag{190}$$

Such that :

$$u_r(t^{\text{peak}}, y=1) - u_r^{\infty}(y=1) = 2 \cdot \frac{q t^{\text{peak}}}{h^i} \cdot \left( \frac{\overline{K}_s}{2} - \overline{K}_s^z \right) \cdot \sum_{n=1}^{\infty} \frac{1}{\mathcal{P}(x_n)} \tag{191}$$

By making use of the decomposition of the linear function r on the basis of the eigenvectors $J_1(xy)$ (Eq.111), we show that:

$$\sum_{n=1}^{\infty} \frac{1}{\mathcal{P}(x_n)} = \frac{1}{4 \cdot \left(1 + \frac{\overline{K}_s^r}{2}\right)} \tag{192}$$

Thus the normalized radial displacement relaxation reads:

$$\frac{u_r(t, y=1) - u_r^{\infty}(y=1)}{u_r(t^{\text{peak}}, y=1) - u_r^{\infty}(y=1)} = 4 \cdot \left(1 + \frac{\overline{K}_s^r}{2}\right) \cdot \sum_{n=1}^{\infty} \frac{1}{\mathcal{P}(x_n)} \cdot e^{-\frac{t-t^{\text{peak}}}{\tau_n}} \tag{193}$$

Moreover, in the relaxation phase, $h = h^{\infty}$ such that the volume reads:

$$V_n(t) = S^{proj}(t) \cdot h^{\infty} \quad \text{with,} \quad S^{proj}(t) \approx (1 + 2 \cdot \frac{u_r(t, y=1)}{b^i}) \cdot S^{proj,i} \tag{194}$$

This directly implies that:

$$\frac{V_n(t) - V_n^{\infty}}{V_n(t^{\text{peak}}) - V_n^{\infty}} = \frac{u_r(t, y=1) - u_r^{\infty}(y=1)}{u_r(t^{\text{peak}}, y=1) - u_r^{\infty}(y=1)} \tag{195}$$

Similarly we can show that,

$$\mathcal{F}^{t^{\text{peak}}} = \mathcal{F}^{\infty} + \frac{1}{\overline{f}^{i}} \cdot \frac{qt^{\text{peak}}}{h^{i}} \cdot \frac{\frac{\overline{K}_s}{2} - \overline{K}_s^{z}}{1 + \frac{\overline{K}_s}{2}} \cdot \left(\overline{f}^{i} + \alpha + \frac{\overline{K}_s}{2}\right) \tag{196}$$

Such that :

$$\frac{\mathcal{F}(t) - \mathcal{F}^{\infty}}{\mathcal{F}^{t^{\text{peak}}} - \mathcal{F}^{\infty}} = \frac{V_n(t) - V_n^{\infty}}{V_n(t^{\text{peak}}) - V_n^{\infty}} \tag{197}$$

We verified this relationship experimentally in Fig. 6A.

##### *7. Determination of $\overline{\Delta P_n^i}$ and $\overline{K}_s$.*

We first use the measured normalized force at peak, $\mathcal{F}^{t^{\text{peak}}}$, and after relaxation, $\mathcal{F}^{\infty}$ to determine $\overline{\Delta P_n^i}$ and $\overline{K}_s$ analytically in the regime where $\alpha \approx 0$. We first re-express $\overline{K}_s^{z} = -\overline{\Delta P}_n^{i} + \overline{\mathcal{U}}$. We showed that :

$$\overline{K}_s^{z} = -\overline{\Delta P}_n^{i} + \overline{\mathcal{U}} \quad , \quad \overline{\mathcal{U}} = \frac{2Ks}{b^{i} \cdot (K + \frac{4}{3}G)} \cdot \left(\frac{3S^{proj} + S^{e,i}}{S^{*}} - \frac{1}{2}\right) \tag{198}$$

For simplicity, we next introduce the variable $x = \frac{b^i}{h^i}$. We then re-express $\mathcal{U}$ as a function of the two variables we are looking for:

$$\mathcal{U} = \frac{(1+2x) \cdot \overline{K}_s + 2 \cdot (1+x) \cdot \overline{\Delta P_n}^{i}}{(1+2x)^2} \tag{199}$$

Thus $\overline{K}_s^{z}$ is of the form:

$$\overline{K}_s^{z} = \alpha_1 \cdot \overline{\Delta P_n}^{i} + \alpha_2 \cdot \overline{K}_s \quad , \quad \alpha_1 = -1 + \frac{2(1+x)}{(1+2x)^2} \quad , \quad \alpha_2 = \frac{1}{1+2x} \tag{200}$$

We next express $\mathcal{F}^{\infty}$ and $\mathcal{F}^{t^{\text{peak}}}$ as :

$$\mathcal{F}^{\infty} = \frac{(1-\tilde{h})\left(1 + \frac{2(-1+\overline{\Delta P_n}^{i})(1+\overline{K}_s\alpha_2+\alpha_1\overline{\Delta P_n}^{i})}{2+\overline{K}_s}\right)}{\overline{\Delta P_n}^{i}} \tag{201}$$

And,

$$\mathcal{F}^{t^{\text{peak}}} = \frac{(-1+\tilde{h})\left(\overline{K}_s(-1+2\alpha_2) + 2(-1+\alpha_1)\overline{\Delta P_n}^{i}\right)}{2\overline{\Delta P_n}^{i}} \tag{202}$$

Thus,

$$\overline{K}_s = \frac{2(-1+\mathcal{F}^{t^{\text{peak}}} + \tilde{h} + \alpha_1 - \tilde{h}\alpha_1)\overline{\Delta P_n}^{i}}{(-1+\tilde{h})(-1+2\alpha_2)} \tag{203}$$

Injecting the latter equation in the equation for $\mathcal{F}^{\infty}$ we obtain an equation for $\overline{\Delta P_n}^{i}$ which we solve to finally find:

$$\overline{\Delta P_n}^{i} = -\frac{(\mathcal{F}^{t^{\text{peak}}} - \mathcal{F}^{\infty})(1-\tilde{h})(-1+2\alpha_2)}{\mathcal{F}^{\infty}(\mathcal{F}^{t^{\text{peak}}} + (\alpha_1 - 1)\cdot(1-\tilde{h})) - (1-\tilde{h})(\alpha_1(1-\tilde{h}) + 2(\mathcal{F}^{t^{\text{peak}}} - (1-\tilde{h}))\alpha_2)} \tag{204}$$

And,

$$\overline{K}_s = -\frac{2(\mathcal{F}^{\infty} - \mathcal{F}^{t^{\text{peak}}})(\mathcal{F}^{t^{\text{peak}}} + (\alpha_1 - 1)\cdot(1-\tilde{h}))}{\mathcal{F}^{\infty}(\mathcal{F}^{t^{\text{peak}}} + (\alpha_1 - 1)\cdot(1-\tilde{h})) - (1-\tilde{h})(\alpha_1\cdot(1-\tilde{h}) + 2(\mathcal{F}^{t^{\text{peak}}} - (1-\tilde{h}))\alpha_2)} \tag{205}$$

Where we remind that:

$$\alpha_1 = -1 + \frac{2(1+x)}{(1+2x)^2} \quad , \quad \alpha_2 = \frac{1}{1+2x} \quad , \quad x = \frac{b^i}{h^i} \quad , \quad \tilde{h} = \frac{h^{\infty}}{h^i} \tag{206}$$

##### 8. *Determination of $\tau_1$ and $\overline{\xi}_s$.*

We designed our experiment so that the amount of volume loss during the compression is small. In this regime, we showed in the previous section that the normalized relaxation of the force reads:

$$\frac{\mathcal{F}(t) - \mathcal{F}^{\infty}}{\mathcal{F}^{t^{\text{peak}}} - \mathcal{F}^{\infty}} = 4\cdot\left(1 + \frac{\overline{K}_s}{2}\right)\cdot\sum_{n=1}^{\infty}\frac{1}{\mathcal{P}(x_n)}\cdot e^{-\frac{t-t^{\text{peak}}}{\tau_1}\cdot\left(\frac{x_n}{x_1}\right)^2} \tag{207}$$

We determined the remaining free parameters, $\tau_1$ and $\overline{\xi}_s$, by performing a least-mean-square fit to the normalized force relaxation data. This normalization was achieved using the previously determined value of $\overline{K}_s$ (Eq. 205).

##### 9. *Discussion on the parameter $\overline{\xi}_s$.*

In our least-mean-square fit, we consistently found that the parameter $\overline{\xi}_s$ was extremely small, with values ranging from $10^{-6}$ to $10^{-3}$. This result indicates that the system is in a limiting regime where the model's output is highly insensitive to this parameter. Consequently, the fit showed no visual or significant numerical improvement between a zero value and the determined value, making it impossible to reliably determine $\overline{\xi}_s$ with our current data.

The fitting method is yielding a small value for $\overline{\xi}_s$ because the experimental data exhibits a strong multimodal response at short timescales. This observed behavior is inconsistent with the model's predictions in the high surface friction regime ($\overline{\xi}_s > 1$), where a single mode would dominate the response (see Section II F).

We next verify whether the model's prediction of a low $\overline{\xi}_s$ is consistent with a theoretical estimate derived from a Poiseuille flow through the NPC. We first estimate an upper bound for the associated surface friction coefficient, $\xi_s$. This calculation is based on a set of typical cellular parameters: a lamina thickness of $H \sim 10-50$ nm, an NPC water permeation radius of $r \sim 5$ nm, a total of $N_{NPC} \sim 3000$ NPCs per HeLa cell nucleus, and an NPC viscosity of $\eta \sim 10^{-1}$ Pa.s. Assuming Poiseuille flow, the permeability per unit pressure gradient for a single pore is:

$$l_p^{1,\text{pore}} \sim \frac{r^4}{\eta H} \tag{208}$$

The corresponding friction of a single pore is thus given by:

$$\xi_s^{1,\text{pore}} \sim \frac{\eta H}{r^2} \tag{209}$$

The total surface friction, accounting for all NPCs, is then:

$$\xi_s \sim \frac{\xi_s^{1,\text{pore}}}{N_{NPC}} \sim 10^5 \text{ Pa.s.m}^{-1} \tag{210}$$

This value is negligibly small when compared to the effective bulk friction we measured, which is on the order of $\xi^f \cdot b^i \sim 10^9$ Pa.s.m$^{-1}$ taking $\xi^f \sim 10^{15}$ Pa.s.m$^{-2}$, and $b^i \sim 10\mu$m. This substantial difference confirms that the water flow through nuclear pores is not limiting nuclear deformations.

##### *10. Determination of the dimensional physical parameters*

By fitting our model to the dynamical data in the limit $t^{\text{peak}} << \tau_1$ and $\alpha \sim 0$, we obtained: $(\tau_1\,,\,\overline{\Delta P}^i_n\,,\,\overline{K}_s)$. We first use the definition of $\overline{\Delta P}^i_n$ and $\Delta P^i_n$ to determine K:

$$\begin{cases} F^i = \Delta P^i_n \cdot S^{proj,i} \\ \overline{\Delta P}^i_n = \frac{\Delta P^i_n}{K} \end{cases} \longrightarrow K = \frac{F^i}{S^{proj,i}\cdot \overline{\Delta P}^i_n} \tag{211}$$

We then use the definition of $\tau_1$ and $D$ to determine $D$ and $\xi_f$:

$$D = \frac{b^{i^2}}{\tau_1 \cdot x_1^2} \quad , \quad \xi_f = \frac{K}{D} \tag{212}$$

$$S^* = S_n^{\text{proj},\,i} \cdot \frac{\left(1+\frac{1}{x}\right)\left(2+\frac{1}{x}\right)\overline{K}_s - \left(4+\frac{3}{x}\right)\overline{\Delta P}^i_n}{\left(1+\frac{1}{2x}\right)\overline{K}_s + \frac{1}{2x}\overline{\Delta P}^i_n} \tag{213}$$

$$K_s = K \cdot h^i \cdot \frac{\left(1+\frac{1}{x}\right)\left(2+\frac{1}{x}\right)\overline{K}_s - \left(4+\frac{3}{x}\right)\overline{\Delta P}^i_n}{\left(2+\frac{1}{x}\right)^3} \tag{214}$$

#### H. Uniaxial compression of polyacrylamide beads.

As a side note we highlight that our theory can also describe the deformation of gel beads under uniaxial confinement by taking the limit $\overline{\xi}_s \longrightarrow 0$ and $K_s \longrightarrow 0$.

### III. METHODS.

#### A. Cell culture.

HeLa Kyoto cells with various markers were cultured in Dulbecco's Modified Eagle Medium enriched with Glutamax (DMEM/Glutamax, Life Technologies) supplemented with 10% v/v fetal bovine serum and 1% v/v penicillin-streptomycin (Gibco), called complete DMEM. RPE1 cells were cultured in DMEM/F12 (Life Technologies) supplemented with 10% v/v fetal bovine serum and 1% v/v penicillin-streptomycin. Both these cell lines were maintained at 37°C with 5% CO2. MDA-MB-231 cells were cultured in Leibowitz-15 medium (L-15, Sigma Aldrich) supplemented with 2 mM glutamine (Gibco), 15% v/v horse bovine serum and 1% v/v penicillin-streptomycin. They were maintained at 37°C with 1% CO2. Cells are detached with Tryple Express (Gibco) prior to experiments.

#### B. Cloning.

The lentivector plasmid pTRIP-SFFV-EGFP-LAP2b coding for the tagged nuclear envelope marker EGFP-LAP2b was obtained by cloning the LAP2b sequence from pEGFP-LAP2b (Euroscarf) into pTRIP-SFFV-EGFP (from Nicolas Manel's lab). Both plasmids were digested with the restriction enzymes BsrgI (New England Biolabs) and KpnI (New England Biolabs) in CutSmart buffer following manufacturer recommendations. Fragments were migrated on a 1% agarose gel and purified, then ligated using the T4 DNA ligase.

#### C. Transduction for establishment of stable cell lines.

Stable cell lines were established by transduction of lentiviral vectors. Lentiviral particles are produced in HEK293FT. HEK293FT are plated in DMEM at 0.8 M / well in a 6-well plate and transfected using the TransIT reagent (Mirrus) with 0.4 $\mu g$ pCMV-VSVG, 1 $\mu g$ psPax2 and 1.6 $\mu g$ of the lentiviral vector of interest. The next day, the medium is changed to 3 mL of the medium of the target cells. The day after, is then added on target cells at the ratio of 2:1 v/v to cell medium, with protamine at a final concentration of 1 $\mu$g/mL. The medium is changed the next day.

#### D. Lamin A/C knock down.

The transfection follows an adapted version of the protocol described in ([12]). Cells are plated at 0.45 M / well in a 6-well plate. They are transduced using the RNAiMAX reagent (Life Technologies) and non-targeting siRNA non-targeting (Horizon Discovery) or targeting LMNA (SMARTpool of 4 siRNA, Horizon Discovery), according to the manufacturer's specifications. The scheme includes two rounds of transfection separated by 48 hours, with a change of medium 24 hours after each transfection. The experiments are performed 48 or 72 hours after the second transfection.

#### E. Western blot.

Protein levels were controlled by Western blot. Cells are harvested by trypsinization and counted to normalize future protein loading by cell number. Typically, 0.5 to 1 million cells were lysed. Cells are washed once in cold PBS and either frozen as dry pellets for later lysis, or directly lysed in RIPA lysis buffer with protease and phosphatase inhibitors (100µL buffer/1 million cells). Lysates are then centrifuged at 11 000 g and the supernatant is transferred to a clean tube. It is then either frozen or used directly. Laemmli 2x is added and the lysates are run on commercial SDS-PAGE 4-12% gels (BioRad) at 90V for 15 min and 120V for 1-1.5 hr. Proteins are transferred on a nitrocellulose membrane using a semi-dry transfer protocol (Invitrogen). Membrane saturation is performed for 1 hr with PBS + Tween 0.1% + 4% BSA. Washes are done with PBS + Tween 0.1%. All antibodies are diluted in PBS + Tween 0.1% + 4% BSA. Primary antibodies are typically used at 1:200 dilutions and are incubated overnight at 4°C. Secondary antibodies are coupled to the Horseradish Peroxydase. They are typically diluted at 1:1000 or 1:5000 (GAPDH) and are incubated for 1 hr at room temperature. Membranes are revealed using the SuperSignal West PicoPLUS reagents (Invitrogen) following manufacturer's recommendations.

#### F. Drugs treatment.

Cells are pre-incubated with the drugs for 20 min before the experiment. Latrunculin A (Sigma) is resuspended in DMSO and used at 2 $\mu M$ unless specified otherwise. Y27632 is resuspended in water and used at 10 $\mu M$ unless specified otherwise. Nocodazole is resuspended in DMSO and used at 10 $\mu M$ unless specified otherwise. Trichostatin A (TSA) was dissolved in DMSO and administered at 100 nM for 16 h prior to measurement.

#### G. Confinement devices.

##### 1. *Static 6-well plate confiner.*

The 6-well confiner is described in ([13]). It uses a glass-bottom 6-well plate (MatTek) and polydimethylsilane (PDMS), prepared from a mix of polymer (RTV, A) and reticulating agent (RTV, B). The device consists of pistons attached to the plate's lid, at the edge of which we position coverslips with a microfabricated PDMS layer with micropillars of controlled height. The pistons are cast in soft PDMS (ratio 37.5:1 w/w) in a metallic mold and cured overnight at 65°C. The 12 mm diameter glass #1 coverslips (VWR) are coated with a microfabricated layer of rigid PDMS (ratio 5:1 w/w). The PDMS is poured on a SU8 wafer obtained by photolithography with the array of micropillars of the desired height. It is cured for 20 min at 95°C. The PDMS is plasma-activated using a plasma cleaner system and incubated with poly-L-Lysine-g-Polyethylene-glycol (pLL-PEG, SuSoS) at 0.1 mg/mL in HEPES at pH 7.4 for 1 hour to prevent cells from adhering. Coverslips are then incubated for at least 1 hour in the medium used for an experiment in the absence of drug treatment and at least 4 hours with drug treatment

##### 2. *Dynamic confiner.*

The dynamic confiner device is also described in ([13]). The device is molded in rigid PDMS (ratio 10:1) to which a coverslip similar to what has been described for the static confiner can be affixed. A glass-bottom 35 mm dish (Fluorodish) is plasma-activated and coated with pLL at 1 mg/mL in HEPES at pH 7.4. Cells are plated on the dish and left for 20 min. Upon connection of the device to a pump, the negative pressure lowers the roof of the device and brings down the micropillars to the glass-bottom 35 mm dish (Fluorodish), imposing the desired confinement height.

##### 3. *Micropatterns.*

Micropatterns were prepared on glass coverslips following the protocol described in ([14]), following the protocol for glass coverslips. Glass coverslips were mounted on bottom-less 35 mm dishes. Cells were plated at 0.5 M/mL and let spread overnight. Imaging was performed the next day on a Leica DMI8 microscope, equipped with CSU-X1 Yokogawa spinning disk module. The acquisition was realized with a 40x oil objective (N.A. 1.30) and collected by the Hamamatsu Orca flash 4.0 camera.

##### 4. *Atomic Force Microscopy (AFM).*

The 35 mm glass-bottom dishes were mounted in a dish heater (JPK Instruments) and kept at 37 °C under an inverted light microscope (Axio Observer.Z1; Zeiss) equipped with a confocal microscope unit (LSM 700; Zeiss) and the atomic force microscope (AFM) head (CellHesion 200; JPK Instruments). Custom-made, wedge cantilevers enable to confine and probe an entire cell, as describe in ([15]) and (Fig. 3A,B & Fig. 5A). These focused ion beam (FIB)-sculpted, flat silicon microcantilevers were processed and calibrated as described in ([16]). The microcantilevers were fixed on a standard JPK glass block and mounted in the AFM head. They were calibrated each time they were being mounted. The confinements were typically done by programming preset target heights and the cantilever was lowered at a speed of 0.5 $\mu$m/s. Images are acquired with a 60x water objective. For the stepwise confinement with target heights 20, 12, 8, 6, 5, 4 and 3 $\mu m$, images were acquired after relaxation of the force signal. Z stacks were acquired at 20 and 12 $\mu m$ and the middle plane was acquired for the other heights.

##### H. Nuclear envelope fluctuations.

Movies of one plane every 250 ms are acquired with a 200 ms exposure at 3-5 % laser power on a Leica DMI8 microscope, equipped with CSU-X1 Yokogawa spinning disk module. The acquisition was realized with a 63x oil objective (N.A. 1.40) and collected by the Hamamatsu Orca flash 4.0 camera. The images are corrected for bleaching using the Fiji plugin Bleach Correction with the Single Ratio method. Single cells are then individualized and registered with the Fiji plugin MultiStackReg (Thevenaz et al., 1998), with the Rigid Body method that corrects for both translation and rotation. On each individual cell, 4 lines were drawn across the nuclear envelope and resliced over time to obtain kymographs of the nuclear envelop at these positions. The NE position at each timepoint is determined by fitting a parabola on an 8-pixel stretch centered on the maximal intensity and taking the abscissa of the parabola peak. The amplitude of fluctuations is calculated as the difference between the 9th and 1st deciles (Fig. 2E,F).

##### I. Nuclear volume.

In this study we used two approaches to estimate nuclear volume upon confinement.

###### *1. Volume estimation using local nuclear height measurement.*

We used this method for the static six-well confinement experiments (Fig. 1C and Supplementary Fig. 1C,D). In this approach, a z-stack of both chromatin and nuclear envelope is acquired with Z step of 0.4 $\mu$m or lower. From this stack, the projected area is measured in the medial plane, and the average nuclear height is determined from the orthogonal (side view) cross-section (Supplementary Fig. 1B). By taking advantage of the fact that nuclei under 8 microns confinement are flattened, the nuclear volume was approximated as the product of the projected area and the average height. We validated this method below 8 microns confinement using 3D reconstructions of the Hoechst nuclear staining (Supplementary Fig. 1A, bottom row). Images were acquired on a Leica DMI8 microscope, equipped with CSU-X1 Yokogawa spinning disk module. The acquisition was realized with a 63x oil objective (N.A. 1.40) and collected by the Hamamatsu Orca flash 4.0 camera. The projected area was obtained by classical thresholding on the Hoechst signal in the middle plane. The average height is obtained from a resliced orthogonal view of the nuclear envelope labeling. A homemade python script detects the position of the nuclear envelope and measures the distance between top and bottom 2 pixels by 2 pixels. It uses the derivative of the pixel intensity to position intensity peaks. An average value is computed for the cell. The volume is calculated as the product of the projected area and the average height.

For the dynamic AFM experiments (Fig. 3A-D), acquiring multiple planes at each confinement step proved too time-consuming. Therefore, we adopted the subsequent method for these dynamic experiments.

###### *2. Volume measurement from the geometrical model.*

We used our geometrical model, to estimate nuclear volume from the maximal projected area of the nucleus and the height of confinement. We first computed the approximated nuclear volume :

$$V_n^{approx}(h, B) = S^{proj}(h, B) \cdot h \tag{215}$$

This approximation is qualitatively expected to be reliable under conditions of high confinement, where nuclei are markedly compressed and adopt a pancake-like shape, and to deteriorate at lower confinement levels (see Supplementary Fig. 1A orthogonal view row). To correct this estimation we inverse Eq.215 to obtain the corresponding $B$ (Eq.9), and then compute the corrected volume $V_n^{(2)}$ using (Eq.18).

We show in Supplementary Fig. 2C-E the inferred parameter $B$, the error $\mathcal{E}^{exp}$ (Eq.216) between the approximated volume $V_n^{approx}$ (Eq.215) and the corrected volume $V_n^{(2)}$ (Eq.18), and the absolute values of both volumes for the LatA cells as an example (N = 3, n = 22 nuclei). We checked that the value at 8 $\mu$m of the corrected volume corresponded within 10 % to the population average values (Fig. 1C) as well as the volume of non-confined nuclei estimated either from 3D reconstructions or simply taking the midplane surface of non-confined nuclei assuming a spherical shape to compute an effective radius. We define the error on the measurement as :

$$\mathcal{E}^{exp}(B) = \frac{V_n^{approx} - V_n^{(2)}}{V_n^{approx}} \tag{216}$$

It is straightforward to show that $\mathcal{E}(B)$ only depends on B. This error is maximal in the case of a sphere ($B >> 1$) $\sim 30\%$ and goes to 0 as the shape converges to a pancake ($B << 1$). We see that the approximated volume is accurate in the range $5- > 3\mu$m (Supplementary Fig. 2D).

#### J. Nuclear shape.

All the measures and calculations detailed below are determined from binary images obtained from manually-controlled threshold on the NE labeling. We explain in the following paragraph the different metrics we used to quantify the amount of nuclear folds (Fig. 4A and Supplementary Fig. 4C,E).

##### 1. *Elliptic Fourier Coefficients ratio.*

Pr. Tanmay Lele and his student kindly provided the MATLAB script used in ([17]), that implements the calculation of the Elliptic Fourier Coefficients (EFC) first described in ([18]), and that we adapted. Briefly, the method relies on fitting successive Fourier ellipses to approximate the shape of the object. The EFC ratio is defined as a ratio of the contribution of the first harmonic compared to all the others:

$$EFC_{ratio} = \frac{\text{major axis}_1 + \text{minor axis}_1}{\sum_{i=2}^{N}(\text{major axis}_i + \text{minor axis}_i)} \tag{217}$$

An arbitrary number of harmonics N, typically the smallest that enables a good Fourier fit to the shapes, is chosen (N=25 in ([18]) and N=15 in ([17]). In order to define the number of harmonics to use, we manually compared the fit quality for several values of N. For very round nuclei, a few harmonics are enough to approximate the shape, while the more folded nuclei required more. We settled for N=25, as a good compromise between the fitting quality and the stability of the measure.

##### 2. *Curvature and distance from convex hull.*

Curvature and distance from convex hull are calculated using a custom Python script. The contour is obtained from binary images thanks to the contour detection function of the Python OpenCV library. From the regularly spaced contour, we calculate the Menger curvature for each point, using and adapting the library developed in ([19]). The Menger curvature is calculated by fitting a circle going through 3 points: the central point and one on each side, at a distance that is adjusted depending on the pixel size. We then plot it against the curvilinear abscissa, raw or normalized to 100 for comparison with other cells. In order to assess the depths of the folds on the contour, we determine the distance of each point on the contour to the convex hull of the nucleus mask. The convex hull and distances are obtained using the Convex Hull and Distance Map functions in Fiji. Further processing and plotting are done by a homemade Python script.

#### K. Nuclear condensates.

##### 1. *Cells transfection and confinement*

*a. Cell culture* All cellular experiments were carried out in human cervical carcinoma HeLa cells (ATCC, CCL-2, RRID:CVCL_0030). Cells were cultivated in Dulbecco's modified Eagle's medium (DMEM, with 4.5 g/L D-glucose, Corning) supplemented with 10% foetal bovine serum (FBS, Cytiva, HYCLSV30160.03) and 1% penicillin/streptomycin (P/S, Sigma P4333), at 37°C in a 5% $CO_2$ humidified atmosphere. Cells were routinely tested for mycoplasma contamination.

*b. Transfection* HeLa cells were seeded 24 h prior to transfection (350,000 cells/well on 22 × 22 mm square coverslips). Cells were transiently transfected with plasmids (2.5 µg for 6-well plates, 1 µg for Ibidi µ-slides) using Lipofectamine 2000 (Invitrogen) and OptiMEM (Gibco, 31985-062) according to the manufacturer's instructions. GFP-DAXX plasmid (500 ng) was co-transfected with a non-expressing ghost plasmid to reach a total of 2.5 µg DNA.

The day before the experiment, FluoroDishes were treated overnight with 100 µL of poly-L-lysine (Sigma-Aldrich, P8920), after being cleaned for 2 min in a plasma cleaner (Harrick Plasma, PDC-32G-2). Thirty minutes before the experiment, 300,000 cells were concentrated in 20 µL of medium and seeded in a FluoroDish (World Precision Instruments, FD35-100) containing 800 µL of medium.

Cells were confined using a dynamic confinement device, as described by [13]. Cells were imaged on a spinning-disk microscope at 60× magnification. A z-stack of images was acquired at 0.4 µm intervals over a distance of 30 µm around the focal plane of the cell. Images of the same cells were acquired before confinement and at 3, 10, and 30 min after confinement using a 485 nm laser.

###### *2. Image analysis*

The analysis was performed using a semi-automated program on Fiji. First, a maximum intensity z-projection was performed. The nuclei were then segmented using a Huang auto-threshold. The condensates were then segmented using Stardist. The number of condensates was obtained by combining this segmentation with FindMaxima for quality control. To obtain more precise information on condensate size, the intensity plot profile was determined for each condensate. The full width at half maximum (FWHM) was determined by doing a Gaussian fit of the profile. Condensate size is calculated using the FWHM. To eliminate aberrant values, the FWHMs are compared to the radius obtained with the ROIs Area, and replaced by this radius if the FWHM is 3 times greater than the radius obtained with the Area.

##### L. AFM Data Collection and Preprocessing

Atomic force microscopy (AFM) measurements were performed across several experimental conditions. The dataset consists of the following:

- Pharmacological Perturbations: Control (DMSO; N=6 independent experiments, n=25 cells), Y-27632 (N=3,n=21), Latrunculin A (LatA; N=6,n=37), and a combined LatA and Trichostatin A treatment (LatA+TSA; N=3,n=22)
- Genetic Knockdowns: Control siRNA treated with DMSO (siCTL-DMSO; N=3,n=26), Lamin A/C knockdown treated with DMSO (siLMNA-DMSO; N=4,n=15), control siRNA with Latrunculin A (siCTL-LatA; N=10,n=45), and Lamin A/C knockdown with Latrunculin A (siLMNA-LatA; N=7,n=32).

For the LatA dataset, two experimental replicates were excluded from the final analysis based on pre-defined quality control criteria. Specifically, one replicate was removed due to ineffective drug action (measured forces were indistinguishable from non-perturbed controls), and another was excluded due to negligible force readings indicative of poor cell viability.

Furthermore, where necessary, we corrected for linear force drift by fitting a first-order polynomial to the baseline and subtracting it from the raw force record.

##### M. Electron microscopy

Cells were fixed in 2% glutaraldehyde in 0.1 M cacodylate buffer (pH 7.4) for 1 h, followed by post-fixation in 1% osmium tetroxide in the same buffer for 1 h. Samples were dehydrated through a graded ethanol series and embedded in epoxy resin. Electron micrographs were acquired using a Quemesa digital camera (SIS) mounted on a Tecnai Spirit transmission electron microscope (FEI Company) operated at 80 kV.

##### N. Statistics and Reproducibility.

*a. Data Presentation* Over the paper data are shown as binned means with 95% CI whiskers. Shaded regions denote the standard deviation (s.d.), representing the population variability within each bin.

*b. Model Parameter Estimation* Parameters inferred from the dynamic model were estimated using a median-based approach to ensure robustness against outliers. Central values for these parameters represent the median of the distribution. Uncertainty in these estimates was quantified via a bootstrapping procedure. Briefly, we performed 1,0000 resamples with replacement from the dataset; Error bars for model-derived parameters represent the 95% CI obtained from the resulting bootstrap distributions. An exception was made for the rose plots (circular histograms), where a 50% CI was utilized for visibility.

Statistical analyses and figure generation were performed using the MATLAB software. Unless otherwise stated, N denotes the number of independent biological replicates, and n denotes the total number of individual nuclei measured.

---
